# Long transposon-rich centromeres in an oomycete reveal divergence of centromere features in Stramenopila-Alveolata-Rhizaria lineages

**DOI:** 10.1101/765032

**Authors:** Yufeng “Francis” Fang, Marco A. Coelho, Haidong Shu, Klaas Schotanus, Bhagya C. Thimmappa, Vikas Yadav, Han Chen, Ewa P. Malc, Jeremy Wang, Piotr A. Mieczkowski, Brent Kronmiller, Brett M. Tyler, Kaustuv Sanyal, Suomeng Dong, Minou Nowrousian, Joseph Heitman

**Affiliations:** Department of Molecular Genetics and Microbiology, Duke University Medical Center, Durham, North Carolina 27710; College of Plant Protection, Nanjing Agricultural University, Nanjing, China; Molecular Biology and Genetics Unit, Jawaharlal Nehru Centre for Advanced Scientific Research, Bangalore, India; Department of Genetics, University of North Carolina, Chapel Hill, North Carolina 27599; Center for Genome Research and Biocomputing and Department of Botany and Plant Pathology, Oregon State University, Corvallis, Oregon 97331; Lehrstuhl fuer Molekulare und Zellulaere Botanik, Ruhr-Universitaet Bochum, Bochum, Germany

**Author notes:** Department of Biochemistry, Robert-Cedergren Centre for Bioinformatics and Genomics, University of Montreal, 2900 Edouard-Montpetit, Montreal, H3T1J4, QC, Canada.

## Abstract

Centromeres are chromosomal regions that serve as platforms for kinetochore assembly and spindle attachments, ensuring accurate chromosome segregation during cell division. Despite functional conservation, centromere DNA sequences are diverse and often repetitive, making them challenging to assemble and identify. Here, we describe centromeres in an oomycete *Phytophthora sojae* by combining long-read sequencing-based genome assembly and chromatin immunoprecipitation for the centromeric histone CENP-A followed by high-throughput sequencing (ChIP-seq). *P. sojae* centromeres cluster at a single focus at different life stages and during nuclear division. We report an improved genome assembly of the *P. sojae* reference strain, which enabled identification of 15 enriched CENP-A binding regions as putative centromeres. By focusing on a subset of these regions, we demonstrate that centromeres in *P. sojae* are regional, spanning 211 to 356 kb. Most of these regions are transposon-rich, poorly transcribed, and lack the histone modification H3K4me2 but are embedded within regions with the heterochromatin marks H3K9me3 and H3K27me3. Strikingly, we discovered a *Copia*-like transposon (CoLT) that is highly enriched in the CENP-A chromatin. Similar clustered elements are also found in oomycete relatives of *P. sojae*, and may be applied as a criterion for prediction of oomycete centromeres. This work reveals a divergence of centromere features in oomycetes as compared to other organisms in the Stramenopila-Alveolata-Rhizaria (SAR) supergroup including diatoms and *Plasmodium falciparum* that have relatively short and simple regional centromeres. Identification of *P. sojae* centromeres in turn also advances the genome assembly.

**Author summary:** Oomycetes are fungal-like microorganisms that belong to the stramenopiles within the Stramenopila-Alveolata-Rhizaria (SAR) supergroup. The *Phytophthora* oomycetes are infamous as plant killers, threatening crop production worldwide. Because of the highly repetitive nature of their genomes, assembly of oomycete genomes presents challenges that impede identification of centromeres, which are chromosomal sites mediating faithful chromosome segregation. We report long-read sequencing-based genome assembly of the *Phytophthora sojae* reference strain, which facilitated the discovery of centromeres. *P. sojae* harbors large regional centromeres fully embedded in heterochromatin, and enriched for a *Copia*-like transposon that is also found in discrete clusters in other oomycetes. This study provides insight into the oomycete genome organization, broadens our knowledge of the centromere structure, function and evolution in eukaryotes, and may help elucidate the high frequency of aneuploidy during oomycete reproduction.

## Introduction

Accurate segregation of chromosomes during mitosis and meiosis is critical for the development and reproduction of all eukaryotic organisms. Centromeres are specialized regions of chromosomes that mediate kinetochore formation, spindle attachment, and sister chromatid segregation during cell division [1, 2]. The DNA coincident with functional centromeres typically consists of unusual sequence composition (e.g. AT-rich) and structure (e.g. repeats, transposable elements), low gene density, and transcription of non-coding RNA (ncRNA) as well as heterochromatic nature [3]. However, an active centromere is defined not by DNA sequences but by the deposition of a centromere-associated protein called centromere protein A (CENP-A, also known as CenH3) [1, 4]. CENP-A is a histone H3 variant, which replaces the canonical H3 in the nucleosomes at centromeres and provides the foundation for kinetochore assembly [1, 5, 6].

Despite the fact that centromere function is broadly conserved, centromeric sequences vary greatly in size and composition, ranging from “point” centromeres of 125 bp in length to “regional” centromeres consisting of up to megabases of repeated sequences to holocentromeres that extend along the entire length of the chromosome [1, 3, 7]. To date, point centromeres have been only reported in the budding yeast *Saccharomyces cerevisiae* and its close relatives, holocentromeres have been identified in some insects, plants and nematodes, represented by *Caenorhabditis elegans*, while regional centromeres are the most common type and found in nearly all eukaryotic phyla [1, 3]. Most animals and plants have large regional centromeres composed of satellite sequences that are organized into a variety of different higher order repeats [4, 8, 9]. Some plant centromeres also possess a different type of repeat called centromere-specific retroelements (CR) [10]. In comparison, all fungal centromeres identified to date do not contain satellite repeats and have diverse organizations. The size of fungal regional centromeres ranges from several kilobases, such as in *Candida albicans*, to hundreds of kilobases in *Neurospora crassa* [11, 12]. The centromeric sequences of fungal regional centromeres can be composed of active or inactive clusters of transposable elements and thus very repetitive, such as in *Cryptococcus* spp. and *N. crassa* [13, 14], or can be nonrepetitive and very short, such as in the wheat pathogen *Zymoseptoria tritici* [15] and *C. albicans* [16]. Information on centromeres is limited in other eukaryotic lineages. The malaria pathogen *Plasmodium falciparum* and the diatom *Phaeodactylum tricornutum* CENP-A binding regions are characterized by short simple AT-rich sequences [17, 18], while the parasite *Toxoplasma gondii* has a simple centromere without nucleotide bias [19].

Due to their highly repetitive nature, assembly of large regional centromeres presents a significant challenge. Emerging long-read sequencing technologies, such as Pacific Bioscience (PacBio) and Oxford Nanopore Technology (ONT), have led to substantial advances in resolution of chromosomal structures including highly repetitive sequences such as centromeres. Using these technologies, centromeres that were difficult to resolve using short-read sequencing, were defined in various organisms, from fungi [13, 20, 21] to insects [22], plants [23] and humans [24].

Oomycetes are fungal-like organisms but belong to the stramenopila kingdom within the Stramenopila-Alveolata-Rhizaria (SAR) supergroup [25, 26]. The SAR supergroup contains a high diversity of lineages that include many important photosynthetic lineages (e.g. diatoms and kelp), and important parasites of animals (e.g., *Plasmodium*, the causative agent of malaria) and plants (e.g. oomycetes, or water molds) [27]. *Phytophthora* is a large oomycete genus (>160 species found to date) and contains some of the most devastating plant pathogens that destroy a wide range of plants important in agriculture, forestry, ornamental and recreational plantings, and natural ecosystems [28]. One notorious example is *Phytophthora infestans*, which caused the Great Irish Potato Famine of the mid-1840s [29]. Today, *Phytophthora* species remain significant threats to major food crops, causing multi-billion US dollars losses annually throughout the world [28, 30]. *Phytophthora sojae* is a widespread soil-borne pathogen of soybean. Because of its economic impact, and tractable genetic manipulation [31–33], *P. sojae* has become a model species to study oomycete genetics, biology, and interactions with plants.

To date, the genomes of more than 20 *Phytophthora* species have been sequenced [34]. Their genomes are generally large and display complex features: they are diploid, highly heterozygous for heterothallic species, and very repetitive, which makes genome assembly challenging. The most contiguous oomycete genome assembly published to date is of the *P. sojae* reference genome, which was generated based on Sanger random shotgun sequencing and subsequent improvements involving gap closure and BAC sequencing [26, 35]. *P. sojae* genome assembly v3.0 (www.jgi.doe.gov) spans ∼82 Mb and contains 82 scaffolds; however, there are ∼3 Mb of unresolved gaps (N’s) persisting in the assembly. Recently, significant progress has been made in genome assemblies of oomycetes based on long-read sequencing [36, 37]; however, the identity or the nature of the DNA sequences that form essential chromosomal elements such as centromeres, remain unknown. In this study, using the evolutionarily conserved kinetochore protein CENP-A as a tool, we investigated cellular dynamics of the kinetochore complex in *P. sojae*, and uncovered the nature of the oomycete centromeres with the aid of long-read genome sequencing and ChIP-seq (chromatin immunoprecipitation followed by high-throughput sequencing) technologies. Our findings suggest that the centromeres of *P. sojae* are divergent from those reported in other SAR lineages, and their features may be used to predict centromeres in other oomycetes.

## Results

### GFP-tagging of CENP-A in *P. sojae* reveals clustered centromeres in different life stages and throughout hyphal growth

Kinetochore protein homologs have been predicted in diverse eukaryotic lineages including oomycete species [38]. To identify kinetochore proteins in *P. sojae*, we conducted BLAST searches against the existing *P. sojae* genome database using the predicted oomycete orthologs as query. Gene models of *P. sojae* kinetochore proteins were examined and corrected based on RNA-seq data when necessary. Protein sequences were verified based on the presence of corresponding motifs (S1 Fig and S1 File).

To examine centromere/kinetochore organization and localization in *P. sojae*, we selected CENP-A, the hallmark of centromere identity in most organisms. The RNA-seq data did not support the gene models of *CENP-A* that was instead verified by 3’-RACE and RT-PCR, followed by Sanger sequencing (S2 A-B Fig). *P. sojae* CENP-A (PsCENP-A) has a conserved C-terminus including the “CENP-A targeting domain” (CATD) (S2C Fig). *GFP* was fused to *CENP-A* at the N-terminus and transiently expressed in *P. sojae* transformants with a constitutive promoter derived from the *Bremia lactucae HAM34* gene (S2D Fig). Overexpressed GFP-CENP-A exhibited nuclear localization with a single fluorescent focus in the nucleus (S2D Fig), suggesting that *P. sojae* has a clustered centromere organization.

We also generated GFP labeled CENP-A expressed from the endogenous locus utilizing CRISPR/Cas9-mediated gene replacement (Fig 1A and S3 Fig). Homokaryotic GFP-CENP-A strains exhibited single GFP foci within nuclei from different *P. sojae* life stages (Fig 1B), confirming that the clustered centromere organization is a feature in *P. sojae.* In addition, we tracked the centromere dynamics during hyphal growth, and found that the clustered centromere pattern was maintained throughout *P. sojae* nuclear division (Fig 1C and S1 Movie).

**Fig 1.**
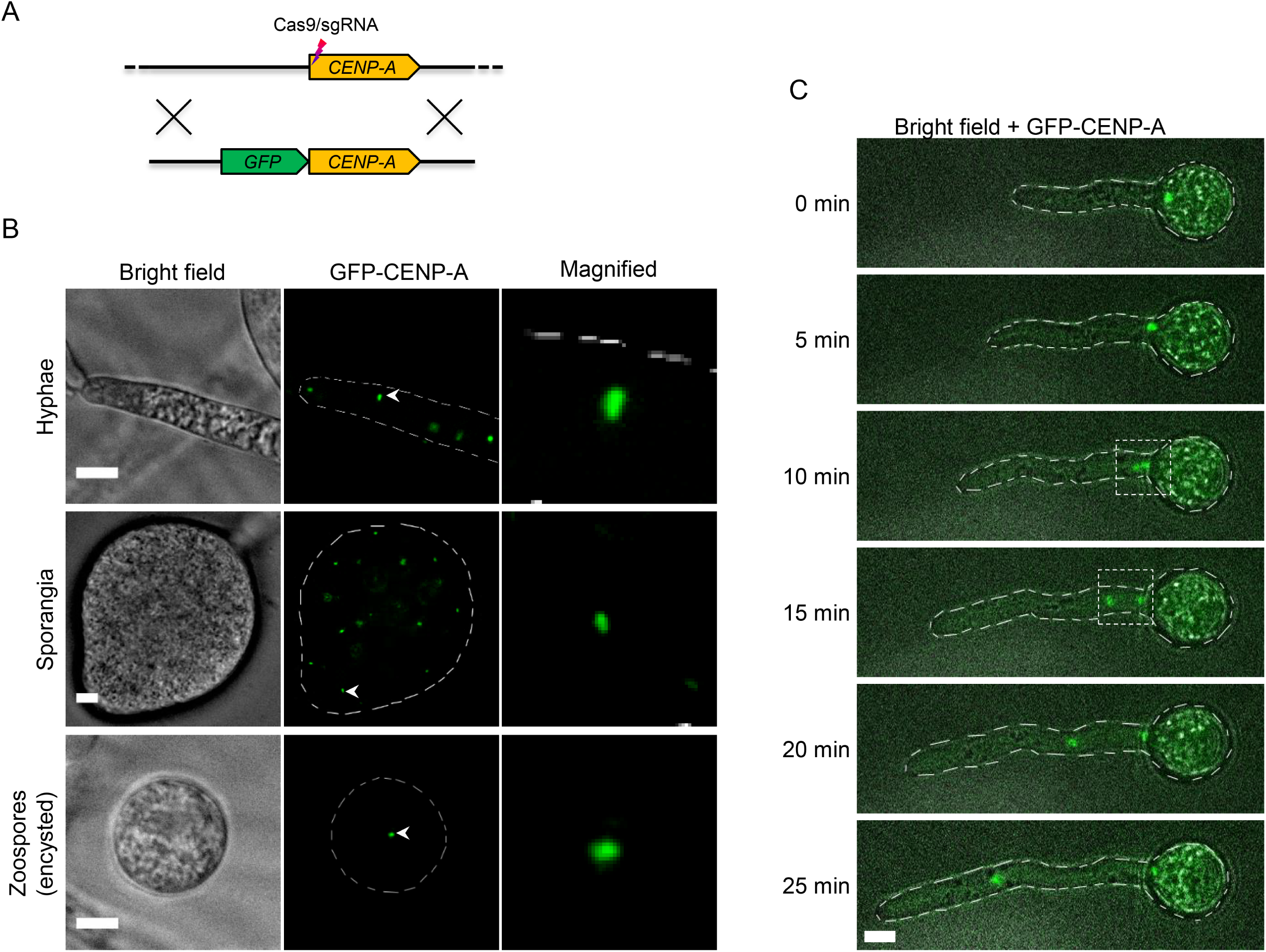
Subcellular localization of CENP-A in *P. sojae* at different life stages and during vegetative growth. (A) A schematic showing the generation of *GFP-*fused *CENP-A* utilizing CRISPR/Cas9 mediated gene replacement. (B) Subcellular localization of GFP-tagged CENP-A (expressed from the endogenous locus) in *P. sojae* hyphae, sporangia, and encysted zoospores. (C) Time-lapse images illustrating localization of GFP tagged CENP-A during hyphal growth. Dashed squares denote occurrence of nuclear division. Representative images are shown. Scale bars in all images, 5 µm.

### Identification of centromeres in a long-read Nanopore-based assembly

To identify *P. sojae* centromeres, we performed native chromatin immunoprecipitation (N-ChIP) using an anti-GFP antibody against the GFP-CENP-A fusion, followed by high-throughput Illumina DNA sequencing. ChIP-seq reads were mapped to the latest Sanger genome assembly (*P. sojae* V3 from JGI), which identified 12 scaffolds that showed relatively concentrated enrichment of CENP-A reads (S4A Fig). CENP-A peaks appeared scattered in Scaffold 1 and Scaffold 11, while more clustered in the other 10 scaffolds. However, further examination of each CENP-A binding region revealed that most of the regions were interrupted by many sequence gaps, which hampered analysis of the sequence features of the candidate centromeres. Thus, we processed to re-sequence and re-assemble the reference *P. sojae* genome.

To improve the genome assembly of *P. sojae* reference strain P6497, we applied Nanopore long-read sequencing and generated a *de novo* genome assembly with SMARTdenovo together with polishing from PacBio and Sanger reads (S5A Fig and S1 Text). The resulting assembly of the nuclear genome (Psojae2019.1) has a size of 86 Mb contained in 70 contigs, with a contig N50 of 2 Mb (S5C Fig). Comparison of Psojae2019.1 to the Sanger assembly indicated that Psojae2019.1 has more repetitive sequences and most regions were collinear (S5 B and S5C Fig, also see S1 Text for details). We also checked telomere repeats using a motif proposed for oomycetes [39], and found 13 contigs (versus 7 in Sanger) that harbor telomeric sequences at single ends (S1 Text and S2 File).

ChIP-seq reads derived from PsCENP-A were mapped to the new genome assembly Psojae2019.1 (S2 Table), which initially revealed 16 regions exhibiting CENP-A enrichment. On closer analysis, we found that the unassembled centromere in contig 20 was an artifact caused by inaccurate genome assembly, as this region was duplicated with a centromere-containing region in contig34 (S6F Fig). Of the 15 remaining CENP-A binding regions, 11 regions were assembled within contigs, whereas four regions were disrupted at the edge of contigs (Fig 2). We confirmed the integrity of the 10 centromeres assembled within contigs as they were completely covered by long reads (S7 Fig), while the CENP-A peaks in Contig 37 and three broken ones (in Contigs 9, 10, 57) lacked sufficient long-read coverage. We focused on the 10 verified CENP-A regions for the further studies (Table 1). RNA-seq analysis indicated that all of the 10 CENP-A regions exhibited low transcription, except the region in Contig 11. Contig 11 contained two adjacent CENP-A peaks, one was 18 kb and the other was 114 kb, which were interrupted by a 21 kb transcriptionally active region (Fig 4C). Here, we define the entire region (two CENP-A peaks, and the region between them) as one centromere (*CEN4*). Among the 10 CENP-A regions, five have a length of ∼190 kb, and three are ∼160 kb, while *CEN3* and *CEN6* are significant larger (>270 kb) (Table 1). All of these centromeres have a GC content comparable to the whole genome (52.16 - 58.13% vs. 54.6%) (Table 1). Taken together, our CENP-A ChIP-seq analysis utilizing the newly assembled genome indicates that *P. sojae* CENP-A prefers to bind large poorly transcribed genomic regions with no specific DNA sequence bias.

**Fig 2.**
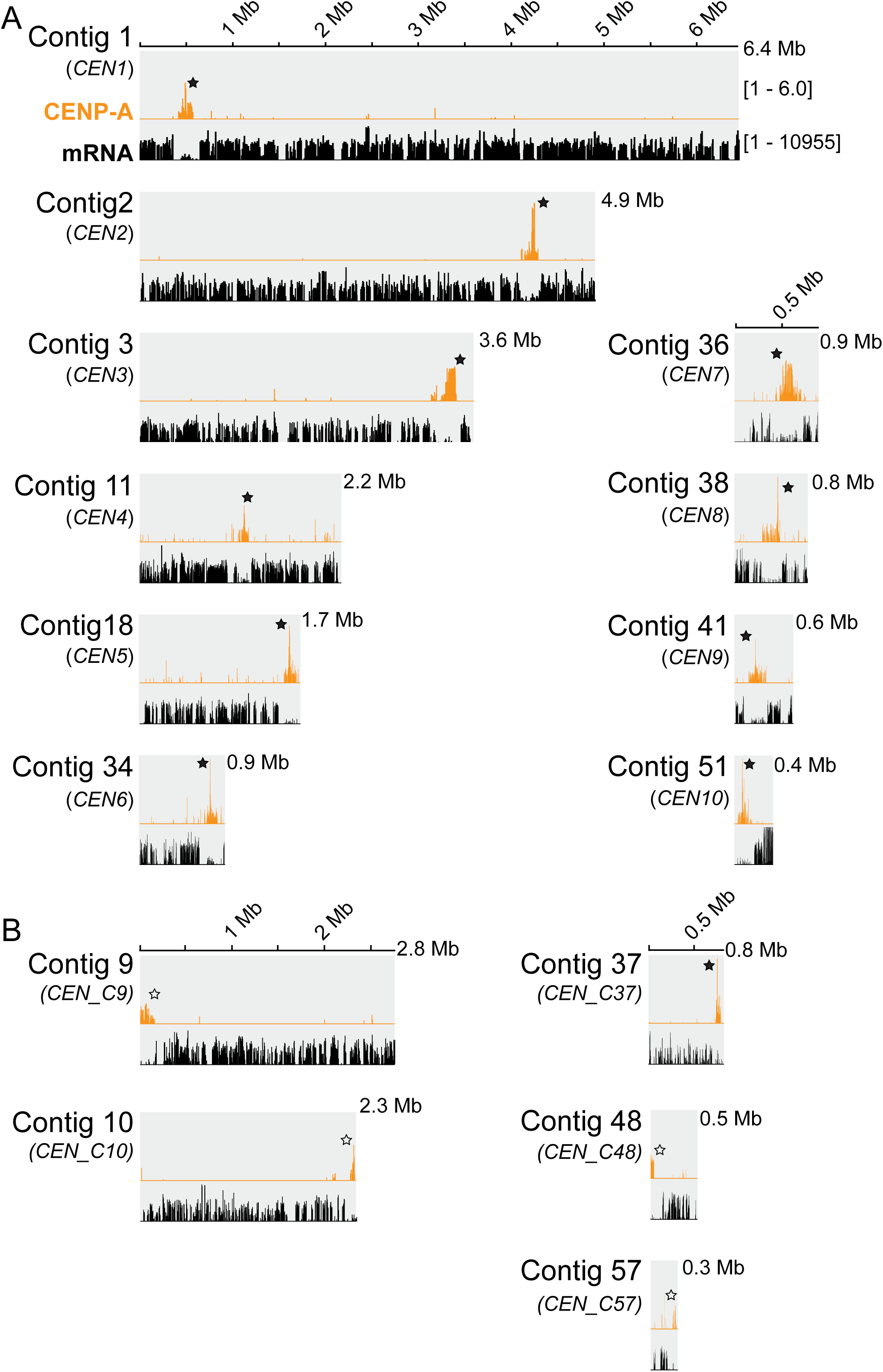
Contigs in the Psojae2019.1 assembly demonstrating CENP-A enrichment based on ChIP-seq. (A) 10 fully assembled CENP-A binding sites presented contigs. (B) 5 incompletely assembled CENP-A binding regions. All contigs are drawn to scale and the ruler indicates the length of the contigs. All CENP-A profiles shown were normalized to input DNA. mRNA profiles are shown as log-scales. Solid stars indicate the CENP-A enriched regions within contigs; hollow stars denote broken centromeres at the edge.

**Table 1.** Centromeres identified in the Psojae2019.1 assembly and their counterparts in the Sanger assembly

| Psojae2019.1 |  |  |  |  | Sanger V3 |  |
| --- | --- | --- | --- | --- | --- | --- |
| Name | Contig | Position of core centromere, kb (size, kb)* | GC% of *CEN* | Position of pericentric region, kb (size, kb) | Scaffold | Position of core centromere (kb)* |
| *CEN1* | 1 | 415-578 (163) | 56.91 | 361-415 (54)<br>579-650 (71) | 2 | 7086-7267 |
| *CEN2* | 2 | 4102-4295 (193) | 55.51 | 4094-4102 (8)<br>4296-4305 (9) | 8 | 2374-2420 |
| *CEN3* | 3 | 3138-3410 (272) | 54.81 | 3223-3138 (15)<br>3412-3560 (48) | 9 | 3075-3286 |
| *CEN4*† | 11 | 991-1174 (183) | 58.13 | 944-991 (47)<br>1175-1205 (30) | 4 | 995-1246 |
| *CEN5* | 18 | 1556-1706 (150) | 57.93 | 1492-1556 (64)<br>1709-? (>22) | 3 | 4078-4138 |
| *CEN6* | 34 | 697-880 (183) | 57.65 | 643-696 (53)<br>882-904 (22) | 6 | 1521-1726 |
| *CEN7* | 36 | 432-706 (274) | 52.16 | 376-433 (57)<br>708-732 (24) | 5 | 2049-2310 |
| *CEN8* | 38 | 302-490 (188) | 57.01 | 288-302 (14)<br>490-515 (25) | 1 | 9667-9688 |
| *CEN9* | 41 | 154-342 (188) | 57.40 | 101-153 (52)<br>342-358 (16) | 1 | 2921-3079 |
| *CEN10* | 51 | 36-151 (115) | 57.93 | ?-36 (>36)‡<br>152-216 (64) | - | - |

\* The coordinates of core centromeres in the Psojae2019.1 assembly are defined by peak calling (S7 File). The positions of centromeres in the Sanger assembly are defined by visualization of peaks.
†Contig11 contains a minor (coordinate, 991465 - 1009035, 18 kb) and a major (coordinate, 1060806 - 1174302, 114 kb) peak that are separated by a 21 kb transcriptionally active region. The core centromere is composed of the minor and major CENP-A regions, and the region between the two peaks.
‡One side of pericentric heterochromatin region is not fully assembled.

To examine the correlation between the centromere regions identified in the new genome assembly and in the Sanger assembly, we conducted synteny analysis using the genomic regions flanking the centromeres. The locations of CENP-A found in the Psojae2019.1 assembly were highly correlated with those in the Sanger assembly, except *CEN10* (Table 1, Fig 3 and S6 Fig). Contig 51 was collinear with the Sanger scaffold 23; however, no enriched CENP-A signal was detected for this scaffold, probably because the region corresponding to *CEN10* is interrupted by gaps. Notably, two CENP-A binding regions in Sanger Scaffold 1 were found to correspond to *CEN8* and *CEN9*, and the smaller one (coordinates: 9,667-9,688 kb) was expanded from 20 kb to 188 kb corresponding to *CEN8* (Table 1, S4B and S6H Figs). In addition, four contigs of the Psojae2019.1 assembly (contigs 4, 38, 23, and 58) are collinear with Sanger Scaffold 1, and telomere repeats are found at the ends of Contigs 4 and Contig 58, further suggesting that Scaffold 1 of the Sanger genome is assembled incorrectly and should be split into two scaffolds (S6H Fig). Overall, comparison of centromeres identified in the Sanger and Psojae2019.1 assemblies further confirms their authenticity and reflects some misassemblies that are present in the Sanger genome assembly.

**Fig 3.**
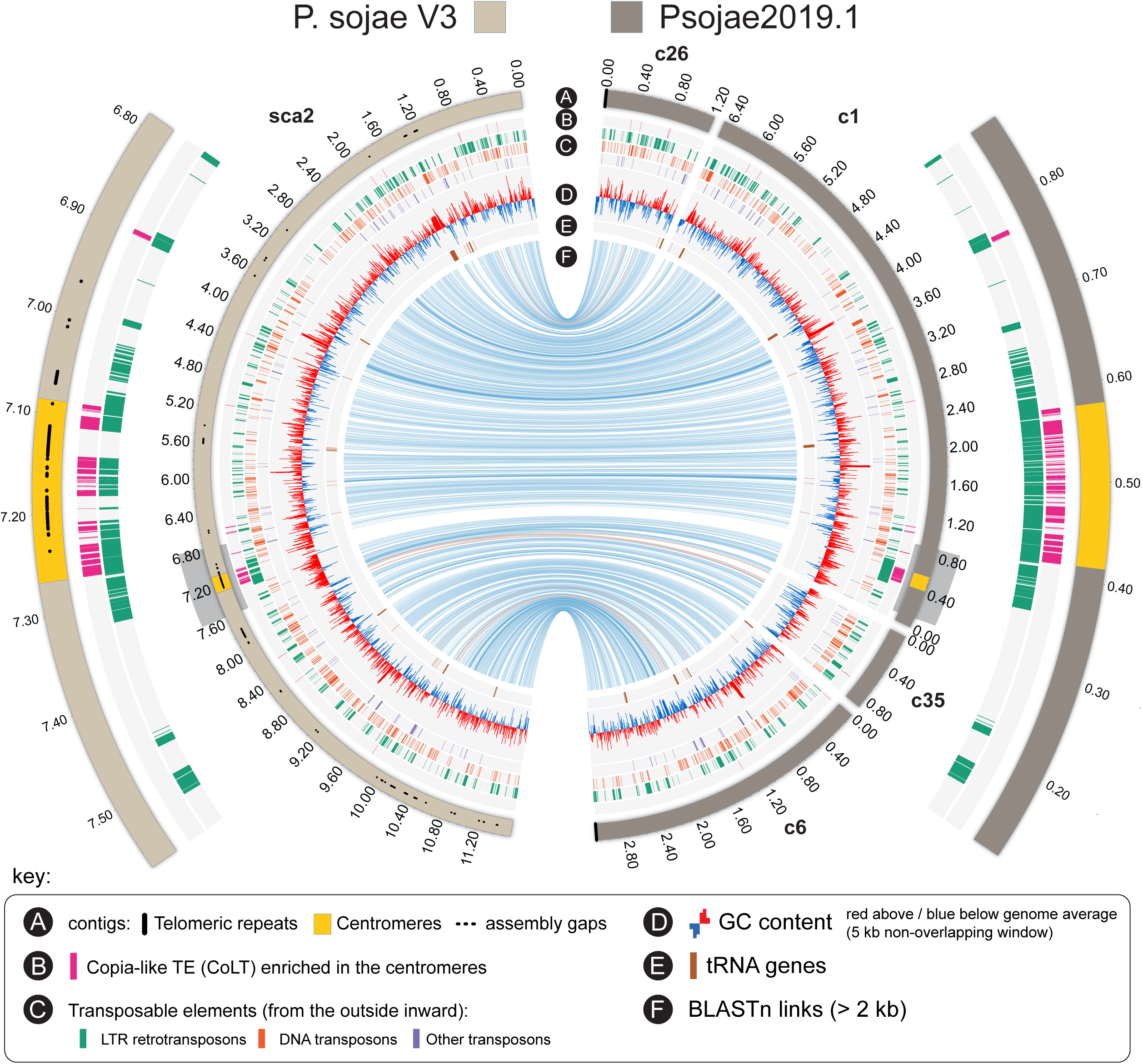
A representative Circos visualization comparing centromere-containing genomic regions between the Sanger V3 Scaffold 2 and the corresponding contigs in the Psojae2019.1 assembly. For the inner circle, track A illustrates assembled contigs (in Psojae2019.1) or scaffold (in P. sojae V3) that are color coded as shown at the top. The locations of centromeres (CENP-A binding regions) are highlighted in yellow. Tracks B-E show the location of other genomic features as given in the key on the bottom. Blue and orange lines in track F link regions with collinearity extending over 2 kb, with orange lines corresponding to inversions. Grey box-shaded centromere-containing regions are magnified for detailed visualization (the two outer arcs).

Due to large genome scales and potentially similar chromosome sizes, the karyotypes of *Phytophthora* species cannot be well resolved by pulsed-field gel electrophoresis [32, 40]. The chromosome number of *P. sojae* is not yet accurately known, but has been estimated to be between 10 and 15 based on an earlier cytological study [41]. By comparing the location of centromeres in the Sanger and Psojae2019.1 assembly, we could validate and predict the configuration of 11 centromeres, namely *CEN1*-*CEN10*, and *CEN_C9* + *CEN_C48* (Table 1 and S4 Table). Three centromeres, namely *CEN_C37*, *CEN_C10* and *CEN_C57*, are not fully assembled. Thus, *P. sojae* is estimated to harbor 12-14 chromosomes, which is in agreement with the previous cytological study.

### *P. sojae* CENP-A regions are embedded within heterochromatin

To define the epigenetic state of *P. sojae* centromeric regions, we performed ChIP-seq with antibodies against two heterochromatin marks (H3K9me3, trimethylation of lysine 9 of histone H3, and H3K27me3, trimethylation of lysine 27 of histone H3) and one euchromatin mark (H3K4me2, dimethylation of lysine 4 of histone H3). The distribution of H3K9me3 and H3K27me3 is generally coincident throughout the genome, and both were colocalized with the CENP-A binding regions (Fig 4 and S7 Fig). Intriguingly, the heterochromatic region extended 8 kb to 64 kb beyond each CENP-A binding region (Table 1 and Fig 4A), similar to pericentromeric heterochromatin regions described in other species [14, 22, 42]. In contrast, the euchromatic mark H3K4me2 was excluded from the CENP-A region and its flanking heterochromatic regions, and generally overlapped with the mRNA transcriptional profile (Figs. 4 B, 4C and S7 Fig). Thus, distribution of histone modifications suggests that the CENP-A regions are embedded in heterochromatin, and we define the heterochromatic regions adjacent to the CENP-A peaks as pericentric regions.

**Fig 4.**
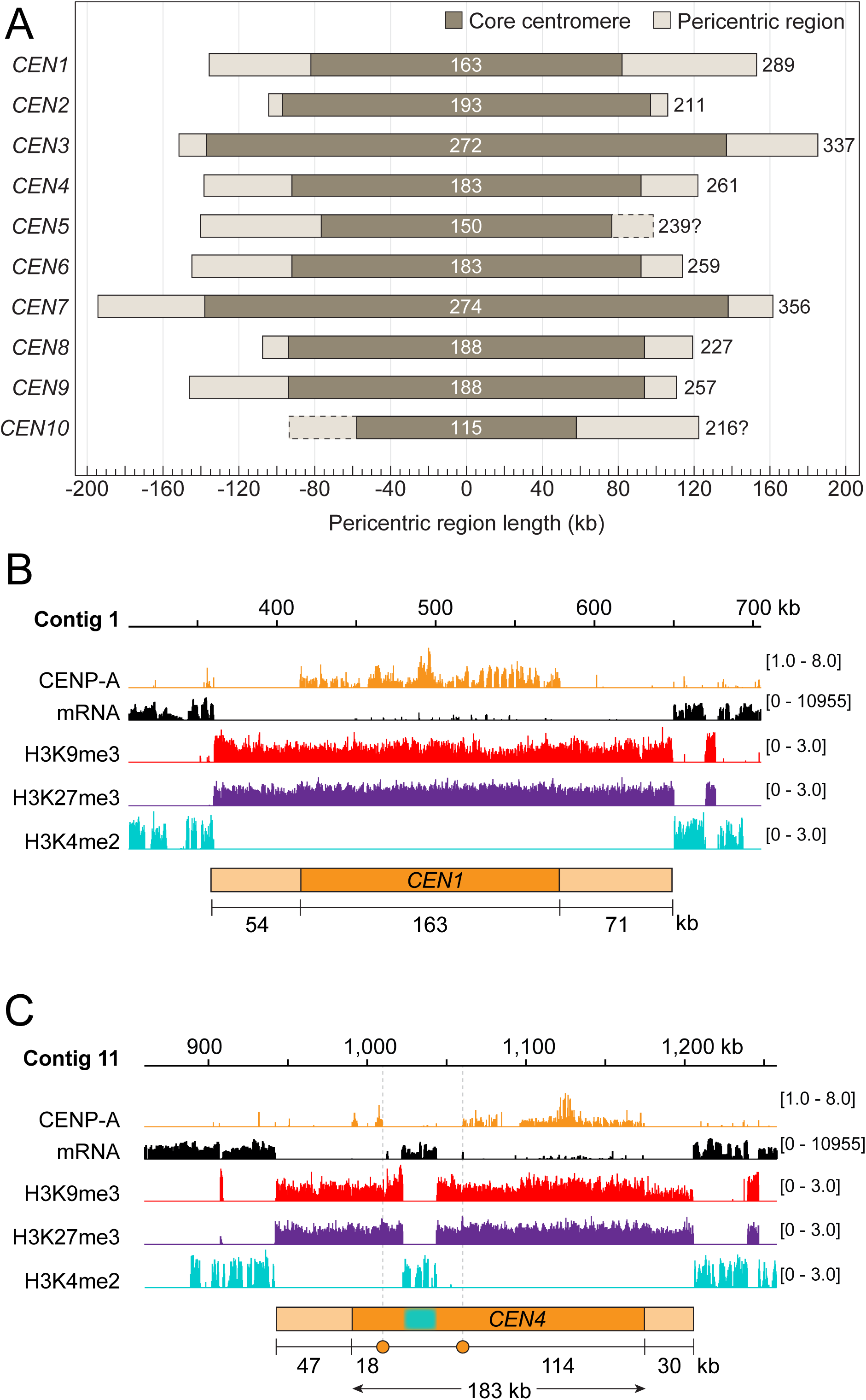
Epigenetic state of *P. sojae* centromeres. (A) Schematics showing the *P. sojae* core centromeres (CENP-A binding regions) and the pericentric regions of various lengths. Dark and light grey bars indicate core centromeric and pericentric regions. Digits at the center indicate the size of core centromeres; digits on the left denote the full-length of the centromeres (a combination of core centromere and pericentromeric region). The right pericentric region of *CEN5* and the left pericentric region of *CEN10*, indicated by dashed bars, are not fully assembled, and their lengths are labeled with question marks. (B-C) Two centromeres (*CEN1* and *CEN4*) are shown as representatives to compare CENP-A localization to the distributions of modified histones. A 400 kb region harboring the centromeric region is shown for *CEN1* and *CEN4*. Cyan block, a transcriptionally active region (21 kb) that interrupts *CEN4*. Profiles of CENP-A, H3K9me3, H3K27me3 and H3K4me2 shown were normalized to input. mRNA profiles are shown as log-scales.

### A *Copia*-like transposon (CoLT) is highly enriched in the *P. sojae* centromeres

The Psojae2019.1 genome assembly contains 31% repetitive sequences, the majority of which are transposable elements (TEs) (S5D Fig). Our analysis showed that centromeres are structurally organized differently, but all composed of many repetitive elements, mostly LTR-retrotransposons (Figs 3, 5A, S6 and S7 Figs). To identify whether the centromeres in *P. sojae* possess any common sequences or repeat elements, all identified CENP-A regions were subject to multiple sequence alignment. This analysis found an ∼5 kb sequence that is highly similar (>98%) and shared among 10 centromeres (S8 Fig and S3 File). BLAST analyses with the consensus 5 kb sequence against the genome revealed that although this element is not exclusive to centromeres, it is significantly enriched in centromeres: approximately 90% of all genomic copies of this element localized to centromeres (Fig 5B). Moreover, this element is present as clusters in centromeric regions, and only sparsely found in other regions of the genome (Fig 5B), further strengthening its association with centromeres.

**Fig 5.**
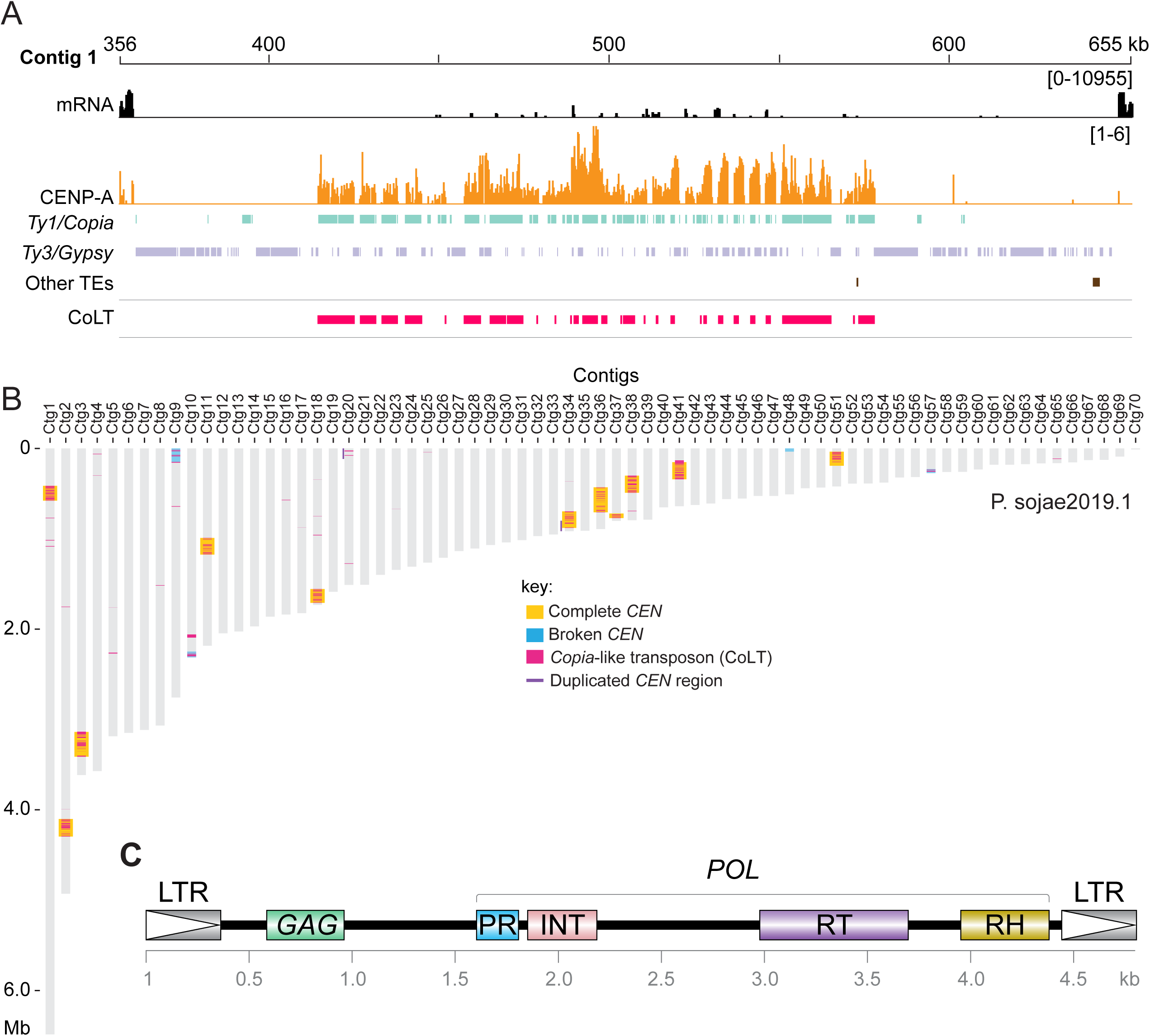
*P. sojae* centromeres are enriched for a *Copia*-like transposon (CoLT). (A) Distribution of transposable elements (TEs) in *CEN1*. TEs were annotated using a *Phytophthora* TE library from Repbase [45]. Tracks of different repeat families are color coded. The track “Other TEs” includes all types of TEs beyond *Gypsy* and *Copia*. CoLT was composed of three elements annotated in Repbase [45], namely Copia-24_PIT-I, Copia-24_PIT-LTR and Gypsy-P17-PR-I. The profile of CENP-A shown was normalized to input. The mRNA track is shown as log-scales and used to define the boundary of pericentric heterochromatin. (B) Location of CoLT elements across all Psojae2019.1 contigs. (C) Diagram showing the domain structure of a representative full-length CoLT sequence (see S3 File). The coding domain is characteristic of *Copia* superfamily of retrotransposons, which consists of capsid protein (Gag), Gag-pre-integrase (PR), integrase (INT), reverse transcriptase (RT), and RNase H (RH) domains, and diverges from the *Gypsy* superfamily in the order of the RT and INT domains in their *POL* genes [86].

In parallel, examination of the 5 kb sequence indicated that it resembled a *Copia*-like transposon that comprises a *GAG* gene, and a *POL* gene encoding PR (protease), INT (integrase), RT (reverse transcriptase), and RH (RNase H) domains in order (Fig 5C and S3 File). To identify the long terminal repeats (LTRs) of CoLT, we analyzed the best Blast hit of the 5 kb sequence. Examination of its flanking nucleotides and the LTR marks (5’ …TG-3’ and 5’ …CA-3’) enabled us to identify the LTRs that are nearly identical (Fig 5C and S3 File). Hence, we named this retroelement CoLT for *<u>C</u>opia*-<u>L</u>ike <u>T</u>ransposon. Phylogenetic analysis based on the conserved RT domains of various reported retroelements confirmed the classification of CoLT as a Ty1/*Copia* retrotransposon (Fig 6A). Notably, among the selected retroelements shown in Fig 6A, *P. sojae* CoLT is distinct from previously studied oomycete *Copia* retrotransposons [43, 44], but clusters together with *P. infestans Copia*-24_PIT-I annotated in Repbase [45], and with predicted CoLT sequences found in other two oomycetes as described below. In addition, phylogenetic tree also suggests that CoLT is close to *Copia* sequences found in plants (Fig 6A).

**Fig 6.**
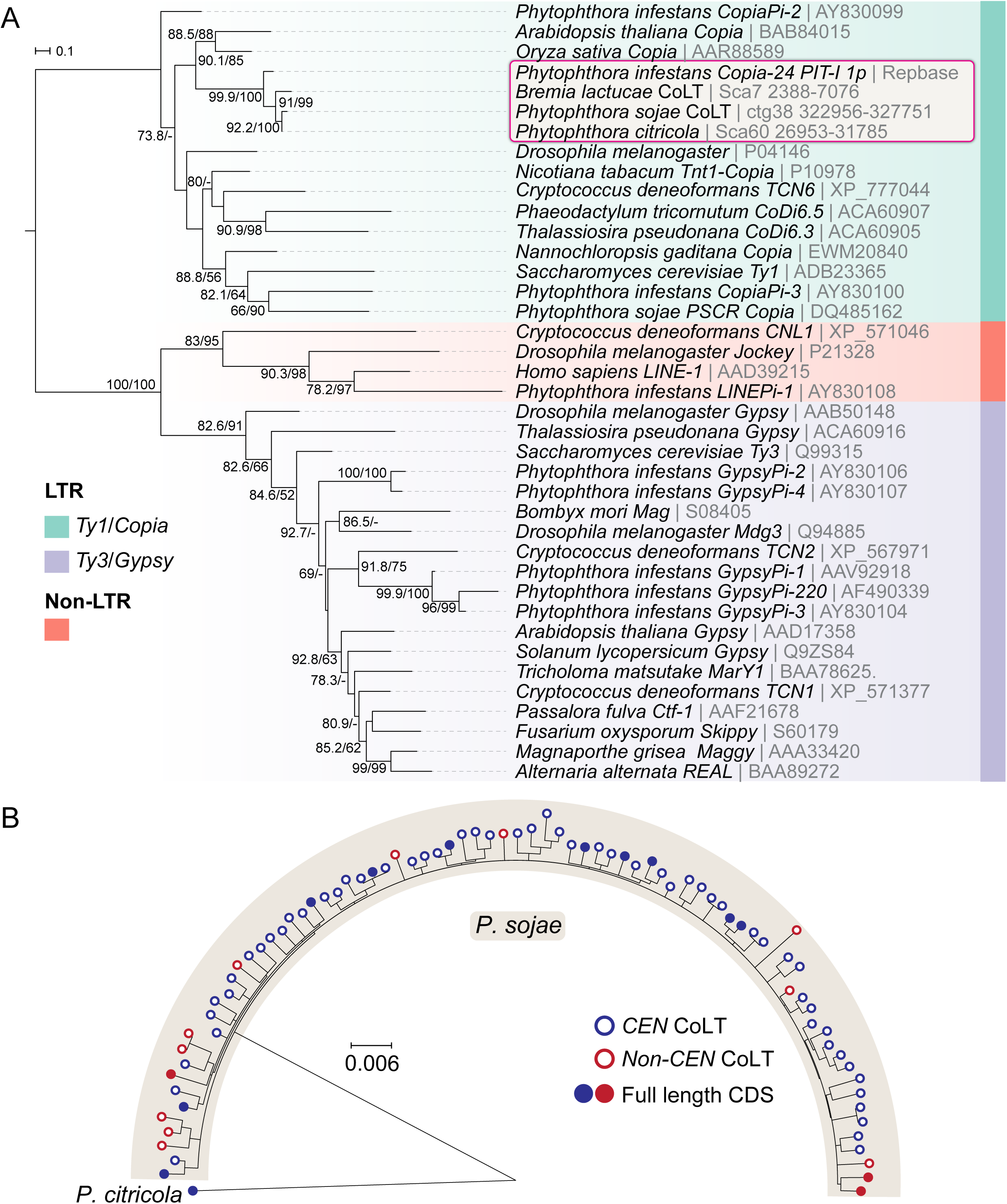
Phylogenetic analyses of CoLT. (A) Maximum-likelihood phylogeny of different retroelements. Protein sequences of the reverse transcriptase (RT) domains were used to construct the tree [87] (see S4A File for the complete DNA and protein sequences). The tree was rooted in the midpoint and branch support values (> 50%) shown at the three nodes were determined by 10,000 replicates of both ultrafast bootstrap approximation (UFBoot) and the Shimodaira-Hasegawa approximate likelihood ratio test (SH-aLRT). (B) Maximum-likelihood phylogeny of full-length CoLT elements identified in the Psojae2010.1 genome assembly. CoLT copies locate inside and outside of centromeres are denoted by blue and red circles. CoLT elements with full-length coding sequences (CDS) are depicted as filled circles. A CoLT sequence identified in *P. citricola* served as an outgroup. See S4A File for the CoLT copies used for the phylogenetic analysis.

Further BLAST analysis employing the full-length CoLT including LTRs as a query indicated that the Psojae2019.1 assembly, in total, harbors 80 CoLT elements of similar length including 11 full-length elements that possess one long ORF encoding all of the *Copia* domains, and 69 sequences that have frameshift mutations within domains (S4B File). Phylogenetic analysis based on the DNA sequences of these elements revealed that the centromeric CoLT copies cannot be distinguished from those found elsewhere in the genome, indicating that centromeric and non-centromeric CoLT copies do not evolve separately; however, it is compelling that almost all full-length CoLT copies (10 out of 11) are found in centromere (Fig 6B). Collectively, our analyses suggest that CoLT, a retrotransposon that harbors all predicted functional domains characteristic for a *Copia*-like transposon, is highly enriched in *P. sojae* centromeres, and is also the only feature shared by all of the centromeres.

### CoLT clusters are conserved in two *P. sojae* oomycete relatives and may be a hallmark of oomycete centromeres

To examine if clustered CoLT elements found in *P. sojae* centromeres are also present in other oomycete genomes, we conducted BLAST searches using the 5 kb consensus sequence derived from *P. sojae* centromeres against the genome assemblies of two *P. sojae* relatives, *Bremia lactucae* (downy mildew, lettuce pathogen) and *Phytophthora citricola* (citrus pathogen), which have relatively contiguous genome assemblies. Interestingly, similar CoLTs clusters were observed in these genomes, and usually appeared once per contig (Figs 7A and 8A). To assess if these clustered CoLTs were syntenic with the *P. sojae* centromere-containing contigs, we examined the CoLT clusters that were present within Mb-long scaffolds/contigs. Synteny analysis demonstrated that five regions in the *B. lactucae* genome that had CoLT clusters were syntenic with *P. sojae* centromeres (Figs 7A). Unexpectedly, Scaffold 2 (original name, SHOA01000004.1, see S5 File for details) contained two CoLT clusters that were syntenic with *P. sojae CEN3* and *CEN5* (Fig 7B), indicating that Scaffold 2 may be incorrectly assembled. It should be noted that the *B. lactucae* genome assembly still has a large percentage of unresolved gaps likely due to its highly heterozygous nature [37]. In comparison, all three selected regions that had clustered CoLT clusters within *P. citricola* contigs (PcContigs) were syntenic with *P. sojae* centromeres (*CEN3*/PcContig2, *CEN9*/PcContig1, *CEN5*/PcContig26) (Fig 8). However, a large number of the CoLT clusters localized at contig ends, or were distributed across the length of short contigs (Fig 8A). This suggests that many of the centromeric regions in *P. citricola* were not fully assembled. Taken together, we propose that the clustered CoLT elements may be used as criteria to predict centromere regions in other oomycete species.

**Fig 7.**
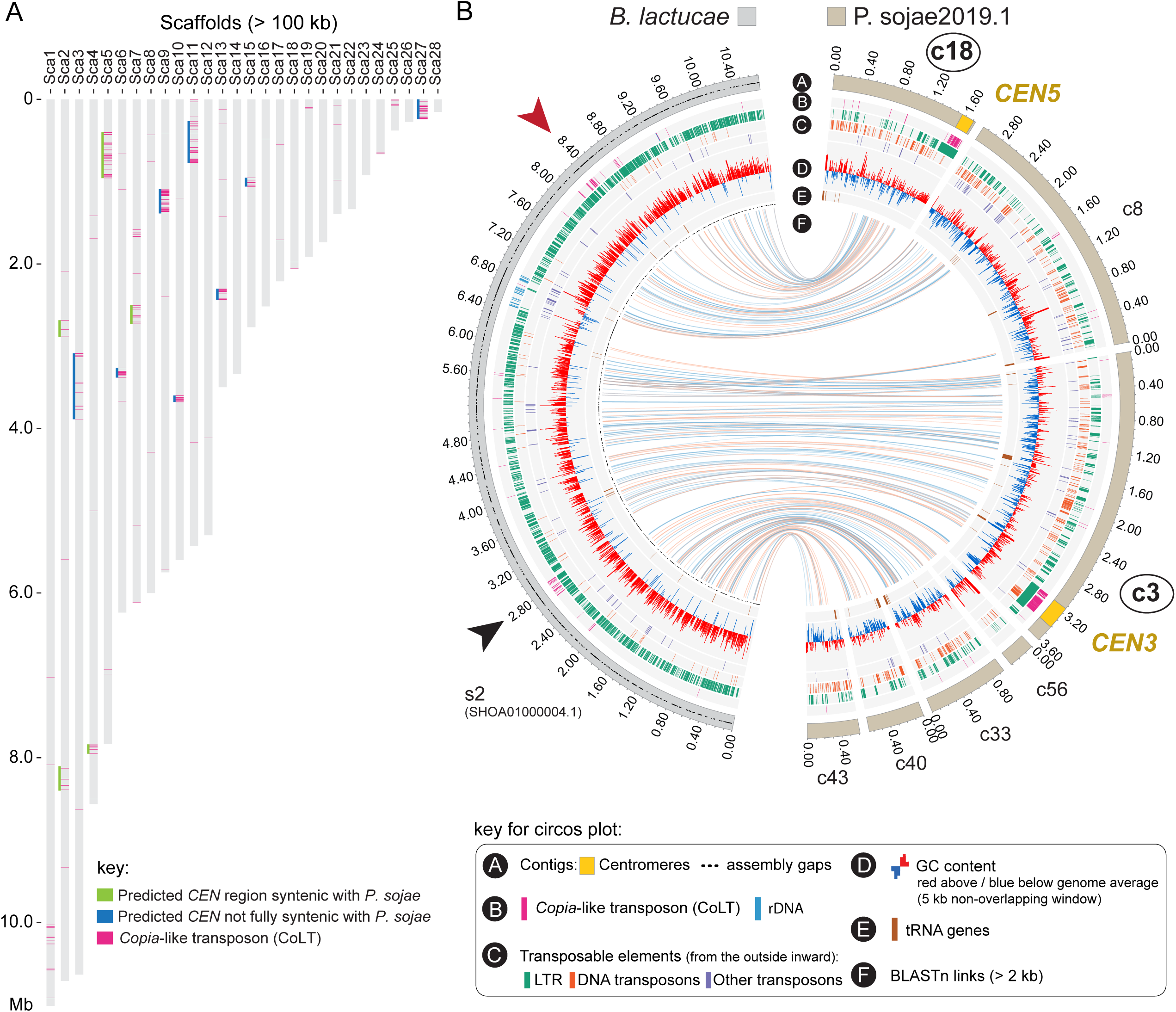
Genomic distribution of CoLT in the *Bremia lactucae* genome. (A) Location of CoLT elements across all *B. lactucae* scaffolds >100 kb. For ease of analysis, scaffolds in *B. lactucae* assembly were sorted and re-named based on sizes (large to small). See S5 File for the original scaffold names. Regions underlined by green lines indicate both sides of the CoLT clusters were syntenic with the regions surrounding *P. sojae* centromeres. Regions underlined by blue indicate only one sides of the CoLT clusters were found to be syntenic with *P. sojae* centromere flanking sequences. (B) A representative Circos plot comparing *B. lactucae* Scaffold 2 that has clustered CoLTs with the corresponding Psojae2019 contigs. The outer track illustrates the assembled scaffold (in the sorted *B. lactucae* assembly) or contigs (in Psojae2019.1), which is color coded as shown at the top. Names of contigs possessing *P. sojae* centromeres are enclosed in circles. Yellow regions on the outer tracks indicate the locations of centromeres (CENP-A binding regions). Blue and orange lines in track F link regions with synteny extending over 2 kb, with orange lines corresponding to inversions. Two CoLT clusters are identified in the *B. lactucae* Scaffold 2 (s2; original scaffold, SHOA01000004.1), which are indicated by arrowheads.

**Fig 8.**
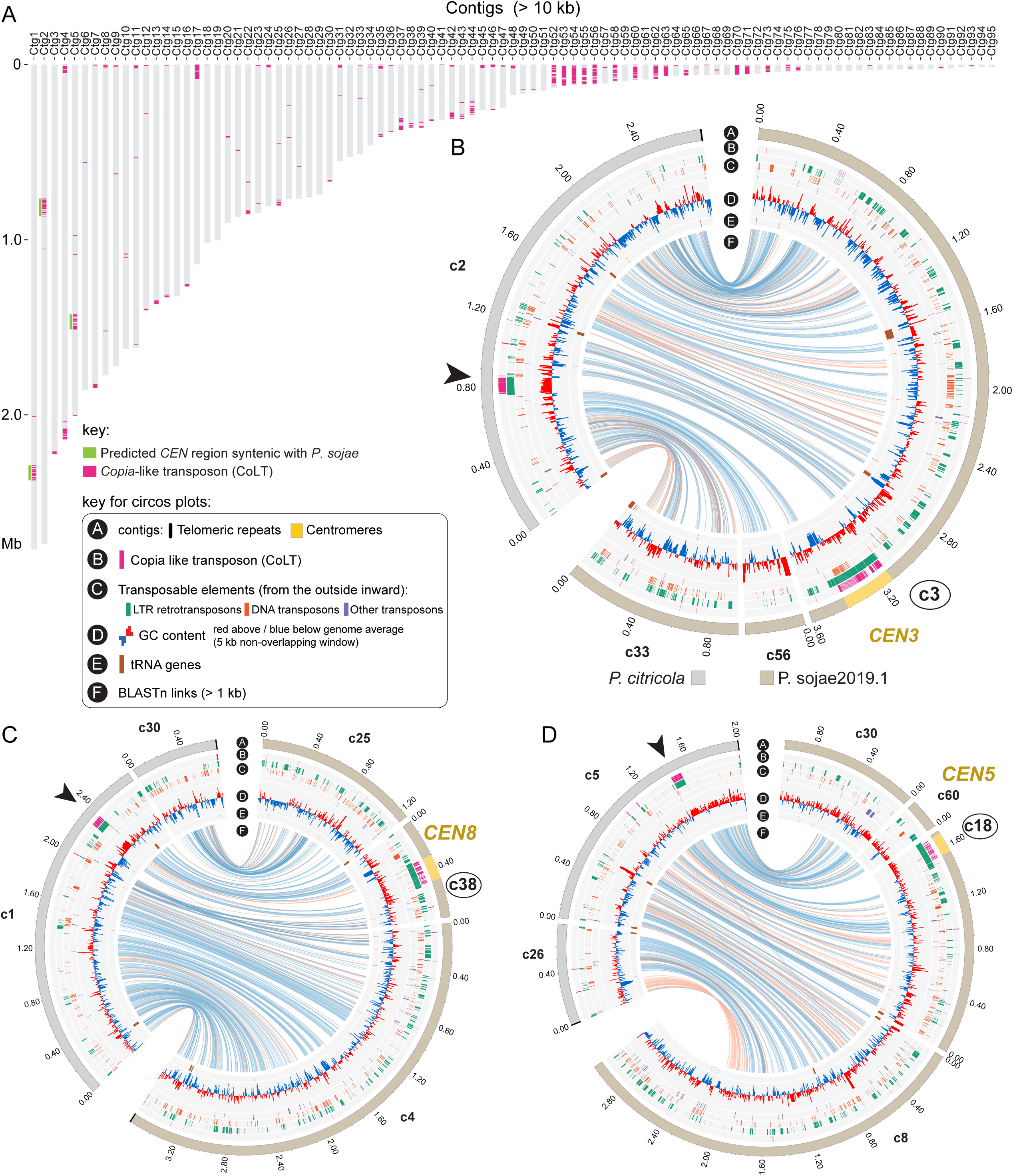
Genomic distribution of CoLT in the *Phytophthora citricola* genome. (A) Location of CoLT across all *P. citricola* contigs >100 kb. (B-D) Circos plots comparing three *P. citricola* contigs that have CoLT clusters with the corresponding Psojae2019 contigs. In each panel, the outer track (bars) illustrates the contigs in the *P. citricola* or the Psojae2019.1 genome assemblies, which is color-coded as shown below the Circos plot in panel B. Yellow regions indicate the locations of centromeres (CENP-A binding regions). Names of contigs harboring *P. sojae* centromeres are enclosed in circles. Blue and orange lines in track F link regions with synteny extending over 2 kb, with orange lines corresponding to inversions. CoLT clusters present in the *P. citricola* contigs are indicated by arrowheads. The flanking sequences of each CoLT cluster are syntenic with the regions surrounding *P. sojae* centromere.

## Discussion

In this study, we identified centromeres in the oomycete plant pathogen *P. sojae* by combining long-read sequencing and ChIP-seq with the GFP tagged kinetochore protein CENP-A. Cellular dynamics analysis revealed that *P. sojae* centromeres were clustered within nuclei in different life stages and during vegetative growth. 10 fully assembled and five incompletely assembled CENP-A binding regions were identified. The common features shared by these regions include: a) a low level of transcription; b) a GC content similar to that of the whole genome; c) repetitive sequences; d) enrichment for a specific *Copia*-like transposon; e) overlapping and surrounding heterochromatin; and f) lack of H3K4me2.

While CENP-A is conserved among different organisms, centromere sequences evolve rapidly [1, 46]. Although the filamentous fungal-like oomycetes are classified in the stramenopiles of the SAR supergroup, it is intriguing to observe that the centromeres that we identified in *P. sojae* are much larger and more complex, comparing to those reported in its stramenopile relative, the diatom *P. tricornutum*, and those found in the parasites (*P. falciparum* and *T. gondii*) of the alveolates (Fig 9). In the latter three cases, all centromeres are composed of non-repetitive sequences. Surprisingly, *P. sojae* centromeres show structural similarity to several, only distantly related, fungal species, such as *N*. *crassa* [14] and *Cryptococcus neoformans* [13]. These features include an enrichment of transposons (or their remnants), and overlap with the constitutive heterochromatin mark H3K9me2/3. Remarkably, the euchromatin mark H3K4me2 has been shown to be associated with centromeres in humans, mouse, *Drosophila*, *S. pombe*, and rice [42, 47–49], but is excluded from other fungal regional centromeres reported to date and in *P. sojae*. In humans and *D. melanogaster*, the CENP-A and pericentromeric heterochromatin domains are spatially distinct, and the CENP-A domain is flanked by but does not overlap with heterochromatin [42, 49, 50]. In contrast, the entire centromere of *P. sojae* is embedded in heterochromatin. It is unknown if the distribution of heterochromatin regions affects centromere distribution in *P. sojae*, but heterochromatin has been shown to be important for centromere function and kinetochore assembly in *N. crassa* and *S. pombe* [14, 51, 52]. In addition, it is of interest that *P. sojae* H3K9me3 and H3K27me3 fully overlap with the centromeric regions, which have not been observed in centromeres of other species thus far, but was shown in human and mouse pericentromeres [9, 53], and in the centromeres of *Z*. *tritici* accessory chromosomes (as their whole accessory chromosomes are enriched for H3K27) [15]. On the other hand, these two epigenetic marks generally coexist throughout the entire genome, suggesting it might be just a general profile of H3K27me3 and H3K9me3 in *P. sojae*.

**Fig 9.**
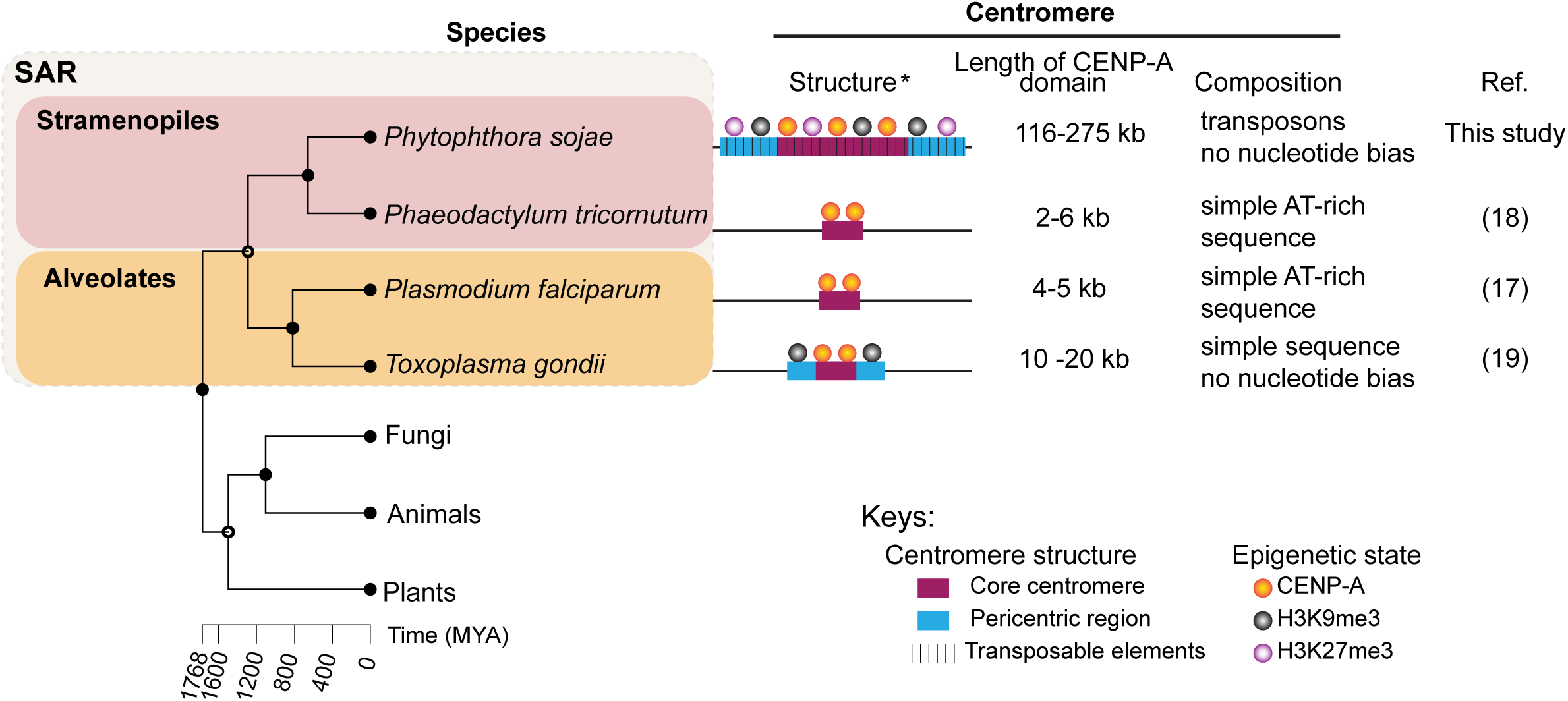
Diversity of centromere features within the SAR supergroup. Simplified schematics (not to scale) showing the structure, epigenetic modifications, size and composition of centromeres across SAR lineages. Asterisk, epigenetic state was not examined in the diatom centromeric regions; however, several AT-rich DNA sequence can be employed for episome maintenance, suggesting diatom centromere might not be epigenetically dependent. Histone modification H3K27me2 was only tested in *P. sojae*. The phylogenetic tree was constructed using TimeTree [88]. *Homo sapiens*, *Arabidopsis thaliana* and *Neurospora crassa* were used as representatives of animals, plants and fungi for the phylogenetic analysis, and are used as outgroups illustrating the evolutionary status of the SAR supergroup.

Transposable elements (and their relics) have been known as residents of the centromeres and pericentromeres of many animals, plants, and fungi [54]. While animal centromeres are associated with both satellite DNA and retroelements, satellite DNA is usually regarded as the main sequence components [55]. Centromeres of many plants, such as maize and rice, are built on centromere-specific retrotransposons (CR), and a certain CR is usually unique to a particular chromosome [8]. Centromeres of *N. crassa* [14] and *C. neoformans* [13] are composed of retrotransposons, and the retroelements in *C. neoformans* are centromere-specific [13]. In comparison, although *P. sojae* regional centromeres include various transposons, many of these elements are not limited to this region and can also be found in other genomic areas. Our study shows that a specific *Copia*-like transposon (CoLT) is highly enriched in the *P. sojae* centromeric regions and confines the CENP-A binding regions (Figs. 5A and S7 Fig). We identified 11 CoLT homologs possessing intact domains that are typical for an active *Ty1*/*Copia* retrotransposon (Fig 5B), indicating that they may be still active. High similarity of LTRs of each CoLT indicate the transposition may occur recently (S3 File). Interestingly, two independent studies employed the central and the 3’-terminal part of CoLT as probes (S3 File) for DNA fingerprinting of *P. sojae* isolates, and polymorphisms were observed by Southern blotting among strains isolated from different geographic locations [56, 57]. This suggest that CoLT may be mobile. In contrast, RNA-seq and histone modification derived from the mycelial stage demonstrated that all of the 11 copies were poorly transcribed and were heterochromatic (Fig 4B and S7 Fig), indicating that the element may be inactive at least during mycelial growth. Therefore, it remains to be determined in which life stages or conditions CoLT becomes active. A similar distribution pattern of centromere-associated retrotransposons was recently found in *Drosophila melanogaster* [22]. In *D. melanogaster*, a non-LTR retroelement named *G2*/*Jockey-3* was found to be enriched in CENP-A chromatin, and this element is also associated with centromeres in its sister species *D. simulans* [22]. Strikingly, the CoLT elements were found to be clustered in the genomes of *P. sojae* oomycete relatives, and some of those regions were syntenic with *P. sojae* centromeric regions. As most of the oomycete genome assemblies were not based on long-read sequencing technology, and thus are very fragmented, it remains to be seen if the CoLT elements have evolved to be widely utilized by oomycetes as a platform for CENP-A loading.

N-ChIP was implemented for this study, because several attempts to perform ChIP analysis based on traditional formaldehyde-cross-linking strategies were unsuccessful. Cross-linking with 1% formaldehyde caused degradation of DNA and failure of ChIP. *P. sojae* transformants expressing GFP tagged CENP-A and CENP-C were both used for N-ChIP-seq. However, only the GFP-CENP-A transformant produced significant enrichment, indicating that the binding of CENP-C to chromosomes may be too weak to recover target DNA under native conditions without cross-linking. Despite most of the ChIP-seq reads of CENP-A are clustered and define the centromeres, we found that additional reads distributed sparsely out of centromeres. This could be caused by mapping, because elements present in the centromeres may also be found in non-centromeric regions, for instance, CoLT. In fact, we observed that all of the CoLT elements including the truncated copies that are present outside of centromeres have peaks of their chromatin marks, and thus are indistinguishable from the centromere-associated ones. Future studies independent of genome assembly, such as fluorescence *in situ* hybridization (FISH) and Hi-C will probably overcome these technical challenges and confirm the identity of confirm the centromere regions mapped by ChIP-seq.

Our analysis showed that having an improved reference genome assembly based on long-read sequencing technologies was crucial to the identification and characterization of centromeres in *P. sojae*. Our attempt to characterize centromere sequences using the classical Sanger assembly was not successful because most of the non-coding repetitive regions were not assembled. While the N50 of the new genome assembly Psojae2019.1 is lower than that of the Sanger assembly, the contigs do not contain gaps and many of the gaps present in the Sanger assembly have been closed (S6 Fig). We tried to scaffold the assembly with different scaffolding programs such as npScarf [58], SSPACE [59], LINKS [60] and the optical BioNano mapping (S1 Text and S9 Fig). Although these scaffolders improved the contiguity (up to 35 scaffolds using SSPACE), they also generated multiple conflicts with the Sanger assembly, and most of the joins could not be supported by evidence such as long read coverage (S3 Table and S10 Fig). Thus, we opted to retain the contig-level assembly in our study. However, identification of centromeres helped to resolve several structural problems present in the “classical” *P. sojae* Sanger assembly, and revealed potential structural problems in other oomycete genome assemblies. On basis of the presence of centromeres and predicted telomeres together with synteny analyses, we found that three Sanger scaffolds/Psojae2019.1 contigs may represent full-length chromosomes, namely Scaffold 2/Contigs [26+1+35+6] (S6A Fig), Scaffold 5/Contigs [17+36+7+49+45] (S6G Fig); and partial Scaffold 1/Contigs [58+38+4] (S6H Fig). Notably, telomeres appear on the both ends of Sanger Scaffold 5 and its syntenic contigs in Psojae2019 (S6G Fig). There are five *P. sojae* centromeres that are not fully assembled. With the development of sequencing and assembly technologies, a finalized chromosome-level genome assembly could help to assemble those broken centromeres, and refine the centromere sequences that we identified.

Centromeres and their associated kinetochore network serve critical functions in genome stability and replication. Failures in kinetochore assembly and attachment increase the probability of chromosome mis-segregation leading to aneuploidy [61]. While these drastic genome changes can be detrimental to the organism, formation of aneuploidy and polyploidy is an important strategy orchestrated by pathogens to adapt to the environment during periods of stress [62]. Polyploidy and aneuploidy are prevalent in *Phytophthora* natural isolates and in progeny from sexual reproduction [36, 63–66]. Interestingly, plant hosts can induce aneuploidy of the sudden oak death pathogen *P. ramorum*, which enhances its phenotypic diversity and increases its adaption to the environment [65]. Recently, a phenomenon termed dynamic extreme aneuploidy (DEA) was described in a vegetable oomycete pathogen, *P. capsici*, in which high variability among progeny produced by asexual spores was caused by ploidy variation [67]. However, the mechanisms resulting in oomycete aneuploidy and/or polyploidy are understudied. As centromeres are the functional and structural foundation for kinetochore assembly and proper chromosome segregation, identification of centromeres and kinetochore proteins in *P. sojae* may help to illuminate the mechanisms underlying oomycete genetic, genomic, and phenotypic diversification.

## Materials and Methods

### *P. sojae* culture and transformation

All the strains used in this study are listed in S5 Table. The reference *P. sojae* isolate P6497 (race 2) used in this study was routinely grown and maintained in cleared V8 media at 25 °C in the dark. Transient gene expression assays based on an optimized polyethylene glycol (PEG) mediated protoplast transformation protocol [31] was applied to examine the nuclear localization of CENP-A. Stable and homokaryotic transformants expressing GFP tagged CENP-A (driven by the strong promoter derived from *HAM34* gene) were chosen for ChIP-seq, which were generated by passaging on V8 supplemented with 50 μg/mL G418 (Geneticin, AG Scientific, San Diego, California, USA) for at least 5 times followed by zoospore isolation. Co-transformation was employed to generate strains expressing both H2B-mCherry and GFP-CENP-A. Transformation was performed as previously described [31]. Sporangia and zoospores were induced by water flooding according to a method described previously [68].

### Construction of plasmids

All the primers used in this study are listed in S6 Table. All GFP fusion constructs were generated based on the plasmid backbone pYF3-GFP [69], in which StuI was used for the N-terminal fusions, and HpaI was used for the C-terminal fusions.

3’-RACE was conducted to validate the gene model of CENP-A, according to the manufacturer instruction (Invitrogen, 18373-019). All PCR-amplifications were performed using Phusion High-Fidelity DNA Polymerase (NEB, M0530S).

### CRISPR-mediated gene replacement

A sgRNA guide sequence whose PAM sequence overlapped with the start codon of CENP-A was selected as the CRISPR/Cas9 targets. An oligo annealing strategy was used for assembly of the sgRNA expression cassettes according to previously described methods [31]. HDR templates for CENP-A was assembled using NEBuilder® HiFi DNA Assembly (NEB, E2621S). 5’-junction, 3’-junction and spanning diagnostic PCR were performed to genotype mutants, utilizing the primers listed in S6 Table.

### Microscopy imaging of *P. sojae* transformants

A Zeiss 780 inverted confocal microscope was adopted to examine the subcellular localization of GFP tagged CENP-A driven by strong promoters. Images were captured using a 63 X oil objective with excitation/emission settings (in nm) 488/504-550 for GFP, and 561/605- 650 for mCherry. DeltaVision elite deconvolution microscope (Olympus IX-71 base) equipped with Coolsnap HQ2 high resolution CCD camera was employed to examine the subcellular localization of GFP tagged CENP-A produced from the native loci. Images were captured using a 100 X oil objective (100x/1.40 oil UPLSAPO100X0 1-U2B836 WD 120 micron DIC ∞/0.17/FN26.5, UIS2) with an excitation filter, 475/28 and an emission filter, 525/50 for GFP. Time-lapse experiments were performed with 40 X oil objective (40x/0.65-1.35 oil UAPO40XOI3/340 1-UB768R WD 100 micron DIC ∞/0.17/FN22, UIS2, BFP1), with the same filters. Confocal images were edited using microscope’s built-in Zen 2012 software (Blue and/or Black edition according to different purposes). DeltaVision images were edited using Fiji-ImageJ and Photoshop.

### High molecular weight genomic DNA extraction and ONT sequencing

High molecular weight (HMW) genomic DNA (gDNA) from *P. sojae* was isolated by the CTAB DNA extraction method. 1 g 3-day old fresh *P. sojae* liquid cultures were collected by filtration and washed twice with sterile water. The resulting damp mycelial pads were frozen immediately in liquid nitrogen in a pre-cooled mortar, then ground by a pestle. Mycelial powder was transferred to a 50 ml Falcon tube and mixed gently with 10 ml room temperature *P. sojae* CTAB extraction buffer (200 mM Tris·HCl pH=8.5, 250 mM NaCl, 25 mM EDTA pH=8.0, 2% SDS, 1% CTAB). The suspension was incubated in 65°C for 15 minutes with mixing every 5 minutes. An equal volume of phenol/chloroform/isoamyl alcohol (25:24:1, saturated with 10 mM Tris, pH=8.0 and 1 mM EDTA) was added to the suspension and mixed gently by inverting the tube, then centrifuge at 4°C, 5000 g for 15min. The supernatant was transferred to a new 50 ml tube and treated with RNase A (final concentration, 100 μg/ml) at 37°C for about 1 hour, followed by proteinase K treatment (final concentration 200 µg/ml) at 50°C for 2 hours. An equal volume of chloroform was added to the solution and mixed gently by inverting the tube, then centrifuge, 4°C, 5000 g for 15min. The supernatant was transferred to a new 50 ml Falcon tube and DNA precipitated by addition of an equal volume of isopropanol. The tube was mixed gently and incubated on ice for 6 hours. The resulting white clump of DNA was spooled by a pipette tip and washed once with 70% ethanol. The gDNA was air-dried for 15 minutes at room temperature and dissolved in 100 µl sterile water. The quantity of DNA was examined by Qubit and the quality was checked by pulsed field gel electrophoresis (PFGE).

1D Genomic DNA by Ligation kits (SQK-LSK108, for MinION); SQK-LSK109, for GridION) were used to prepare the Oxford Nanopore library. Oxford Nanopore sequencing runs was performed on SpotON R9.4 flow cells with MinKNOW V1.11.5 using MinION or SpotON R9.4.1 flow cells with MinKNOW V3.1.20 using GridION. All of the GridION sequence were basecalled (on GridION, in real time) using Guppy v2.0.5.

### Native ChIP-seq

Native ChIP was performed according to the ChIP protocol accompanying Gent, Wang (70) with modifications. Briefly, 1-3 mg mycelia were collected from 1-1.5 L of ∼3-day culture by filtration system, and ground into fine powder in liquid nitrogen with pre-chilled mortars and pestles. Nuclei were isolated and digested by micrococcal nuclease (M0247S, NEB) at 37 for 6 min. An antibody against GFP (Abcam, ab290) was used to immunoprecipitate single nucleosomes containing the GFP-CENP-A fusion. Antibodies H3K9me3 (Abcam, ab8898), H3K27me3 (Active Motif, 39157), and H3K4me2 (Millipore, 07-030) were used to immunoprecipitate nucleosomes with relevant modifications. ChIP-seq of GFP-CENP-A and H3K27me3 were performed by Genewiz using Illumina NextSeq500 that generated 150 nucleotide paired-end reads (80%-85% mappability, S2 Table); ChIP-seq of H3K9me3 and H3K4me2 were conducted by BGI using Illumina Hiseq 4000 that produced 50 nucleotide single-end reads (∼98% mappability, S2 Table). Numbers of reads for each sample are listed in S2 Table.

### Analysis of ChIP-seq and RNA-seq

To map ChIP-seq reads to the genomes, the quality of raw ChIP-seq reads were first assessed by FastQC (v0.11.6). For ChIP-seq of CENP-A, H3K27me3, the resulting reads were trimmed with fastx-clipper and mapped with Bowtie2 with default parameters [71] and aligned to the genome assemblies. For H3K9me3 and H3K4me2, the ChIP-seq reads were polished by BGI prior to be released and thus mapped to the genomes directly using the same Bowtie2 setup. In case of replicates, the ChIP-seq reads were mapped randomly among all replicates. The aligned file (.bam) was sorted and indexed by samtools (version 1.9). Subsequently the ChIP-ed and input samples were analyzed with DeepTools (v3.2.0) ‘‘bamCompare’’ to calculate normalized ChIP signal (log2[ChIP_RPKM_/Input_RPKM_]) and bigwig files were generated. Then .bw files were visualized using the Integrative Genome Viewer (IGV). (https://software.broadinstitute.org/software/igv/). Core centromeres (CENP-A binding regions) were defined by one continuous stretch of ChIP-seq peaks (> 5 kb) in scaffolds or contigs in the Sanger or Psojae2019.1 genome assembly. The boundaries of the 10 fully assembled CENP-A regions were defined by peaking calling employing MACS v2.2.5 (–broad -q 0.00001 --broad-cutoff 0.00001). Peaks with enrichment fold smaller than 2 were filtered out (S7 File). The boundaries of pericentric heterochromatin were defined by the start of both RNA-seq and H3K4m2 peaks. To get profile mRNA, the existing RNA-seq reads (SRA: SRR10283202) derived from the mycelial stage were aligned to the genomes using HISAT2 (version 2.1.0), and the resulting files (.bam) were sorted and indexed by samtools (version 1.9). The .bam file was converted to .tdf for visualization using IGV.

### Genome assembly, analysis of genomic features and synteny comparison

Details of the *de novo* genome assembly is described in SI Text. To predict gene models, first, the assembly Psojae2019.1 was subjected to repeat masking utilizing RepeatMasker [72] based on a library of *de novo*-identified repeat consensus sequences that was generated by RepeatModeler (www.repeatmasker.org/RepeatModeler.html). Next, the repeat-masked assembly was used to predict gene models *ab initio* based on MAKER (v2.31.18) [73] with predicted proteins from available *P. sojae* and *P. infestans* genome annotations as input [26, 74]. GC content was calculated in non-overlapping 5 kb windows using a modified Perl script (gcSkew.pl, https://github.com/Geo-omics/scripts/blob/master/AssemblyTools/gcSkew.pl) and plotted as the deviation from the genome average for each contig. Genes encoding ribosomal RNA (18S, 5.8S, 25S, and 5S) and tRNA were inferred and annotated based on RNAmmer (v1.2) [75] and tRNAscan-SE (v2.0) [76], respectively. To find telomeres, a custom-made Perl script was used to search for the sequence “TTTAGGG” that was proposed for oomycetes telomeric sequences [39]. Pairwise synteny comparison between the two *P. sojae* genome assemblies (i.e. P. sojae V3 and Psojae2019.1) or between different oomycete species was conducted using BLASTn. BLASTn hits and other genomic features were plotted using Circos (v0.69-6) [77]. Whole-genome alignment was computed with MashMap (https://github.com/marbl/MashMap) employing default settings, and was visualized as a dot plot [78].

### Bionano mapping

*P. sojae* protoplasts were generated from 2.5-day old mycelial and were embedded into agarose. Bionano Prep Cell Culture DNA Isolation Protocol was employed for extracting the high molecular weight DNA. DNA labelling with DLE-1 was performed according to the standard protocols provided by Bionano Genomics (Document number 30206, version F). Labelled DNA samples were loaded into two flow cells and run on a Saphyr system (Bionano Genomics). The *de novo* assembly was performed using Bionano Solve 3.3. Standard parameters for Saphyr data were used without “extend and split” and without haplotype refinement in order to create a single map for each allele (“optArguments_nonhaplotype_noES_DLE1_saphyr.xml”). In the process of *de novo* assembly, data generated from two flow cells were merged. An assembly graph was generated during a pairwise comparison of all of the molecules with a p value threshold of 1e-11, and was refined based on molecules aligned to the assembled maps with a p value threshold of 1e-12. After five rounds of extension and refinement, a final refinement was conducted with a p value threshold of 1e-16. Then, the *de novo* assembled map was used to scaffold the sequence assembly. When using the hybrid scaffold module of Bionano Solve 3.3 pipeline, the option of “resolve conflicts” for sequence contigs and Bionano maps was selected. The standard hybrid scaffold settings with a modified parameter (-E 0) was applied to remove discrepancies between sequence assembly and Bionano *de novo* assembly. Sequence contigs were *in silico* digested, based on the recognition sequence (CTTAAG) of DLE-1. Conflicts detection was accomplished by aligning contig maps to Bionano maps with p value threshold of 1e-10. When divergence was identified, the conflicts were resolved by cutting either the contig or the map, depending on the quality of the genome map at the divergent position.

### Analysis of transposable elements and identification of CoLT

To identify transposable elements in *P. sojae*, the new genome assembly was subjected to RepeatMasker (Repbase v23.09) analysis and hits were mapped to this genome assembly. The *Copia*-like transposon (CoLT) element was identified in a stepwise way by multiple sequence alignments followed by extraction of a consensus sequence and BLASTn analyses. Specifically, an approximately 5 kb consensus sequence was identified in the alignment of centromere sequences (including incompletely assembled ones) utilizing the multiple alignment program MAFFT, a plug-in in the Geneious R9 software (http://www.geneious.com), with default parameters. Then the consensus sequence was used as a query to perform a BLASTn search against the Psojae2019.1 genome assembly. The resulting sequence hits were used to map against the genome, and hits longer than 500 bp were used for representing in the figures. The longest sequence hit with highest identity was retrieved, and was used as a query to execute a second round of BLASTn search against the NCBI database to further characterize the sequence. The results of BLASTn analysis indicated that that the sequence was highly similar to a *Copia*-like transposable element. To define the domains of the CoLT, this sequence was further analyzed by repeat identification (utilizing a bioinformatics software Unipro UGENE [79]), and by searches utilizing the Repbase database (https://www.girinst.org/) and NCBI CD-search (https://www.ncbi.nlm.nih.gov/Structure/cdd/wrpsb.cgi).

To validate the classification of CoLT as a *Copia* retrotransposon, phylogenetic analyses were performed based on the alignment of the reverse transcriptase (RT) domain of the identified *P. sojae* CoLT, and of LTR (*Gypsy* and *Copia*) and LINE retrotransposons previously characterized in other organisms. NCBI accession number of each TE sequence is shown in Fig. 6A. CoLT sequences and the deduced RT domains of *P. sojae*, *B. lactucae* and *P. citricola* are shown in S3 File. A total of 39 protein sequences were aligned with MAFFT v7.310 [80] employing the L-INS-i strategy (--localpair --maxiterate 1000) and poorly aligned regions were removed with TrimAl (-gappyout) [81]. A maximum-likelihood phylogeny was inferred using the LG+I+G4 model of amino acid substitution in IQ-TREE v1.6.5 [82]. Branch support values were obtained from 10,000 replicates of both the ultrafast bootstrap approximation (UFboot) [83] and the SH-like approximate likelihood ratio test (SH-aLRT) [84]. Additionally, to investigate if the centromeric CoLT sequences have a common origin, BLASTn was conducted using one full-length CoLT (See S3 File for the sequence). All of the full-length CoLT copies were extracted, and hits with less than 95% sequence identity and shorter than 95% of the query length were removed. The obtained genomic coordinates were sorted, merged using ‘bedtools merge’, and were employed to retrieve the corresponding nucleotide sequences with the aid of ‘bedtools getfasta’ from the bedtools package (https://bedtools.readthedocs.io/en/latest/) [71]. CoLT nucleotide sequences were subsequently aligned using MAFFT, as described above, and the alignments were manually inspected. A ML phylogenetic tree was constructed based on 80 *P. sojae* CoLT sequences (66 located within the defined centromeric regions and 14 located elsewhere) using IQ-TREE and the TN+F+R2 model of DNA substitution. A full-length CoLT sequence retrieved from the *P. citricola* genome using a similar procedure was used as outgroup (S4 File). Truncated CoLT elements (i.e. without a full-length coding sequences) were inferred upon automatic translation of the predicted coding region (see S4 File). All trees were plotted with iTOL v4.3.3 [85].

### Prediction of centromeric regions in *P. sojae* closely related species

To predict centromeres of the two oomycete species, namely *Phytophthora citricola* P0716, (Genbank: GCA_007655245.1, with permission of the author) and *Bremia lactucae* SF5, (GenBank: GCA_004359215.1) [37], BLASTn searches were conducted utilizing the *P. sojae Copia*-like transposon (CoLT) as a query. Significant hits (>90% identity and > 500 bp) were retrieved, and were plotted to all scaffolds of the *B. lactucae* assembly and to contigs > 10 kb of the *P. citricola* assembly. For CoLT clusters that were localized within scaffolds or contigs, their collinearities with the Psojae2019.1 assembly were further examined with BLASTn, and visualized by Circos.

### Data availability

The Nanopore sequencing raw reads are available in NCBI under the BioProject PRJNA563922; All raw ChIP-seq reads can be accessed under NCBI’s SRA listed in S2 Table. The Psojae2019.1 genome assembly has been deposited at DDBJ/ENA/GenBank under the accession WWEI00000000.

## Acknowledgments

We thank BioNano Genomics Support, in particular, Yuanyuan Chang for technical support of BioNano mapping, Beth Sullivan at Duke University for critical reading and comments on the manuscript, and members of the Heitman, Sanyal and Garre labs for helpful discussions. M.N. would like to thank Ulrich Kück and Christopher Grefen for support by the Department of General and Molecular Botany/Molecular and Cellular Botany of Ruhr University. B.M.T. and B.K. were supported by Oregon State University. These studies were supported by NIH/NIAID grants R01 AI050113-15 and R37 MERIT award AI039115-21 to J.H and by the German Research Foundation (DFG, grant NO407/7-1 to MN). J.H. is also a Co-Director and Fellow of the CIFAR program, Fungal Kingdom: Threats & Opportunities.

## Supporting information

**S1 Fig. Summary of the presence and absence of putative core kinetochore proteins identified in *P. sojae.*** Kinetochore orthologs were identified based on BLAST searches. *P. sojae* CENP-A (in bold) was selected to track subcellular localization of centromere/kinetochore and profile centromere sequences. Sequences of *P. sojae* putative core kinetochore proteins are listed in S1 File.

**S2 Fig. Identification and expression of CENP-A in *P. sojae*. (A) Image of *CENP-A* model and RNA-seq log scale coverage taken from FungiDB.** Arrows and triangle denote an erroneous *P. sojae* CENP-A gene model caused by an intron that was missed in the gene model prediction. (B) Electrophoresis image showing 3’-RACE result of *CENP-A*. 5’-primer JOHE45057 (not shown in scale) served as a gene-specific primer for 3’-RACE (See S6 Table). (C) Alignment of *P. sojae* CENP-A with orthologs from different organisms. Ps, *P. sojae*; Pt, diatom (*Phaeodactylum tricornutum)*; Sc, *Saccharomyces cerevisiae;* Cn, *Cryptococcus neoformans; Hs, Homo sapiens;* At, *Arabidopsis thaliana*; Dm, *Drosophila melanogaster.* (D) Transient expression of GFP tagged CENP-A in *P. sojae* transformants. Upper panel, a plasmid constructed for transient expression of *CENP-A*. Expression of *P. sojae* CENP-A is driven by a constitutive promoter derived from the *B. lactucae HAM34* gene. Lower panel, subcellular localization of GFP-tagged CENP-A in the *P. sojae* transformants based on the constructs in (A).

**S3 Fig. Generation of *P. sojae* strains expressing *GFP* tagged *CENP-A* utilizing CRIPSR/Cas9 mediated genome editing.** (A) Schematic of gene replacement of the endogenous *CENP-A* with *GFP-CENP-A*. Lightning bolts, sgRNA guideline sequences were designed overlapping the start codon of *CENP-A*. Primer pairs, JOHE50062/JOHE45358, JOHE45420/JOHE50063, JOHE50062/JOHE50063 were used for 5’-junction, 3’-junction, and spanning diagnostic PCR screening *GFP-CENP-A* mutants. See S6 Table for the primer information. (B) Representative genotyping results of zoospore isolated (homokaryotic) *GFP-CENP-A* strains (YFP10a1 and YFP10b1, see S5 Table for their genetic background). Products observed in the 5’- and 3’-junction PCR of wild type (WT) are non-specific amplicons.

**S4 Fig. Scaffolds in the Sanger assembly that are suggested to harbor putative centromeres. (A) CENP-A enrichment identified in 12 scaffolds.** Solid and hollow stars denote CENP-A enrichment regions that are sequence-gap free or contain gaps, respectively. All CENP-A profiles shown have been normalized to input. mRNA profiles are shown as log-scales. (B) Two putative centromeric regions in Scaffold 1 were identified by poor transcription and the syntenic regions in the Psojae2019.1 assembly.

**S5 Fig. Pipeline used for *de novo* assembly and metrics of the *P. sojae* genome assembly Psojae2019.1.** (A) Pipeline used for *de novo* assembly of Psojae2019.1. Box in grey, different scaffolding programs were employed to enhance the contiguity of the assembly (See more details in S10 Fig and S3 Table). As some of them generated conflict and sequence gaps, we opted to use the contig-level assembly for the centromere study. (B) Dotplot comparison of the *de novo* assembly Psojae2019.1 against the Sanger assembly. (C) Genome assembly metrics. *Statistics for both the Psojae2019.1 and Sanger V3 assemblies were calculated by QUAST [89]; †Annotation was based on the repeat-masked assembly (See method). ‡Annotation was obtained from FungiDB release 33 (https://fungidb.org/fungidb/). §Measured by RepeatModeler (See methods). (D) Pie chart summarizing retroelements (LINE, SINE and LTR), DNA transposons and other repeat sequences predicted in the Psoaje2019.1 assembly.

**S6 Fig. Comparison of centromere-containing genomic regions between the Sanger (P. sojae V3) and the Psojae2019.1 assemblies.** In all panels (A-H), the outer tracks illustrate assembled contigs (in Psojae2019.1) or scaffolds (in P. sojae V3), and are color coded as given in key on the bottom. Yellow regions on the outer tracks indicate the location of centromeres (CENP-A binding regions). Blue and orange lines in track F link regions with synteny extending over 2 kb, with orange lines corresponding to inversions. Names of contigs that contain *P. sojae* centromeres are enclosed in circles. Arrowheads indicate the shrunk centromeres present in the Sanger scaffolds. (A) Comparison of Sanger Scaffold 2 (sca2) and its syntenic contigs in the Psojae2019 assembly. (B) Comparison of Sanger Scaffold 8 (sca8) and its syntenic contigs in the Psojae2019 assembly. (C) Comparison of Sanger Scaffold 9 (sca9) and its syntenic contigs in the Psojae2019 assembly. (D) Comparison of Sanger Scaffold 4 (sca4) and its syntenic contigs in the Psojae2019 assembly. Dots under *CEN4* indicate the regions showing CENP-A peaks, as a transcriptionally active region is found in *CEN4*. (E) Comparison of Sanger Scaffold 3 (sca3) and its syntenic contigs in the Psojae2019 assembly. (F) Comparison of Sanger Scaffold 6 (sca6) and its syntenic contigs in the Psojae2019 assembly. Dashed lines under Psojae2019.1 Contigs 20 and Contig 34 indicate duplicated regions. (G) Comparison of Sanger Scaffold 5 (sca5) and its syntenic contigs in the Psojae2019 assembly. (H) Comparison of Sanger Scaffold 1 (sca1) and its syntenic contigs in the Psojae2019 assembly. The two centromeric regions in Scaffold 1 of the Sanger assembly (P. sojae V3) suggested by CENP-A ChIP-seq are indicated by arrows. Part of the Contig4 is inverted in the Psojae2019.1 assembly compared to the Sanger assembly (inversion breakpoint indicated by an asterisk). Intriguingly, both versions can be supported by some of the mapped reads indicating that both ends of the inverted region are repeat-rich and composed of the same types of repeats so that reads map to both versions. Alternatively, this may represent a structural rearrangement between the two haplotypes of the diploid genome, or it could also represent a real difference between the P6497 culture used for the Sanger sequence and the one used for the ONT assembly.

**S7 Fig. Summary of features of each intact centromere and read coverage analysis of centromere.** For each panel (A-J), upper, an overview of the contig exhibiting enrichments of both CENP-A and histone modifications (H3K9me3, H3K27me3 and H3K4me2), and distribution of transposable elements. TE, transposable elements. CoLT, *Copia* like transposon. Middle, a magnified view of the region shaded in the contig shown above; a diagram displaying the correlation of core centromere (CENP-A) and heterochromatic regions. A 400 kb region is shown for each centromere that has assembled pericentromeric regions on both sides. A 270 kb region is shown for *CEN5*, as it is missing one side of the intact pericentric region. The length of Contig51 is close to 400 kb, only the entire contig is shown (J). Bottom, an image showing long-read coverage. Canu-corrected Nanopore reads were mapped to all centromeres using Minimap2 [90], except *CEN7*, which was verified by all Nanopore and PacBio reads (without Canu-correction) using GraphMap [91]. For better visualization, indels of reads were masked. Asterisk in (B), regions (underlined) that have high CENP-A enrichment but no CoLT and very low TE density contains unknown repetitive sequences unique to *CEN2*.

**S8 Fig. MAFFT-based alignment of CENP-A binding regions reveals a 5 kb sequence that are highly similar among *P. sojae* centromeres.** Upper panel, alignment of CENP-A binding regions. Lower panel, magnified view of the conserved 5 kb region. Images were adapted from the alignment result generated by Geneious.

**S9 Fig. Representative contigs that are anchored by BioNano mapping and contigs that are suggested to be joined.** (A) Contig 2 representing most of the cases that only parts of contigs were anchored by Bionano molecules. (B) One case that the two contigs were not anchored by BioNano but suggested to be combined. (C-E) Three contigs that were anchored by BioNano and suggested to be combined. (E, F) Two cases showing that contigs can be fully covered by BioNano molecules. Map102 was a fragment trimmed from Contig 32. NGS, next generation sequencing, i.e. Psojae2019.1; BNG, BioNano *de novo* assembly; HYBRID, hybrid scaffold.

**S10 Fig. Dot plot comparison of scaffolded assemblies against the original Psojae2019.1 assembly and the Sanger assembly.** (A-D) Assemblies generated by the scaffolding programs npSarf, SSPACE and LINKS, and BioNano mapping were aligned trespectively to the original SMARTdenovo assembly (Psojae2019.1), as well as the Sanger genome, and plotted utilizing the MUMmer package [92].

S1 Table. Metrics of ONT sequencing

S2 Table. Statistics of ChIP-seq samples

S3 Table. Metrics of scaffolded assemblies and their comparison to the Sanger V3 and Psojae2019.1 assembly.

S4 Table. Five incompletely assembled centromeres in the Psojae2019 assembly and their corresponding CENP-A regions mapped in the Sanger assembly

S5 Table. *P. sojae* strains used in the study S6 Table. Primers used in this study

S1 Text. Nanopore sequencing and *de novo* assembly of the reference *P. sojae* genome

S1 Movie. Time-lapse experiment showing cellular dynamics of CENP-A during *P. sojae* vegetative growth.

S1 File (Separate file). Sequences of kinetochore orthologs identified in *P. sojae*.

S2 File (Separate file). 13 telomeres predicted in the Psojae2019.1 assembly.

S3 File (Separate file). The 5 kb consensus sequence shared by *P. sojae* centromeres, and representative full-length CoLT homologs in *P. sojae*, *B. lactucae* and *P. citricola*.

S4 File (Separate file). Sequences used for building phylogenetic trees in Fig 6. (A) Sequences employed for the phylogenetic analysis in Figure 6A. (B) Genomic coordinates of the sequences used for constructing phylogenetic tree in Figure 6B.

S5 File (Separate file). Original names of the sorted *B. lactucae* scaffolds.

S6 File (Separate file). Bionano mapping report.

S7 File (Separate file). CENP-A peaks called by MACS in the 10 contigs that harbor fully assembled centromeres

