## Supplementary figures and images for "Long transposon-rich centromeres in an oomycete reveal divergence of centromere features in Stramenopila-Alveolata-Rhizaria lineages"

### S1 Fig

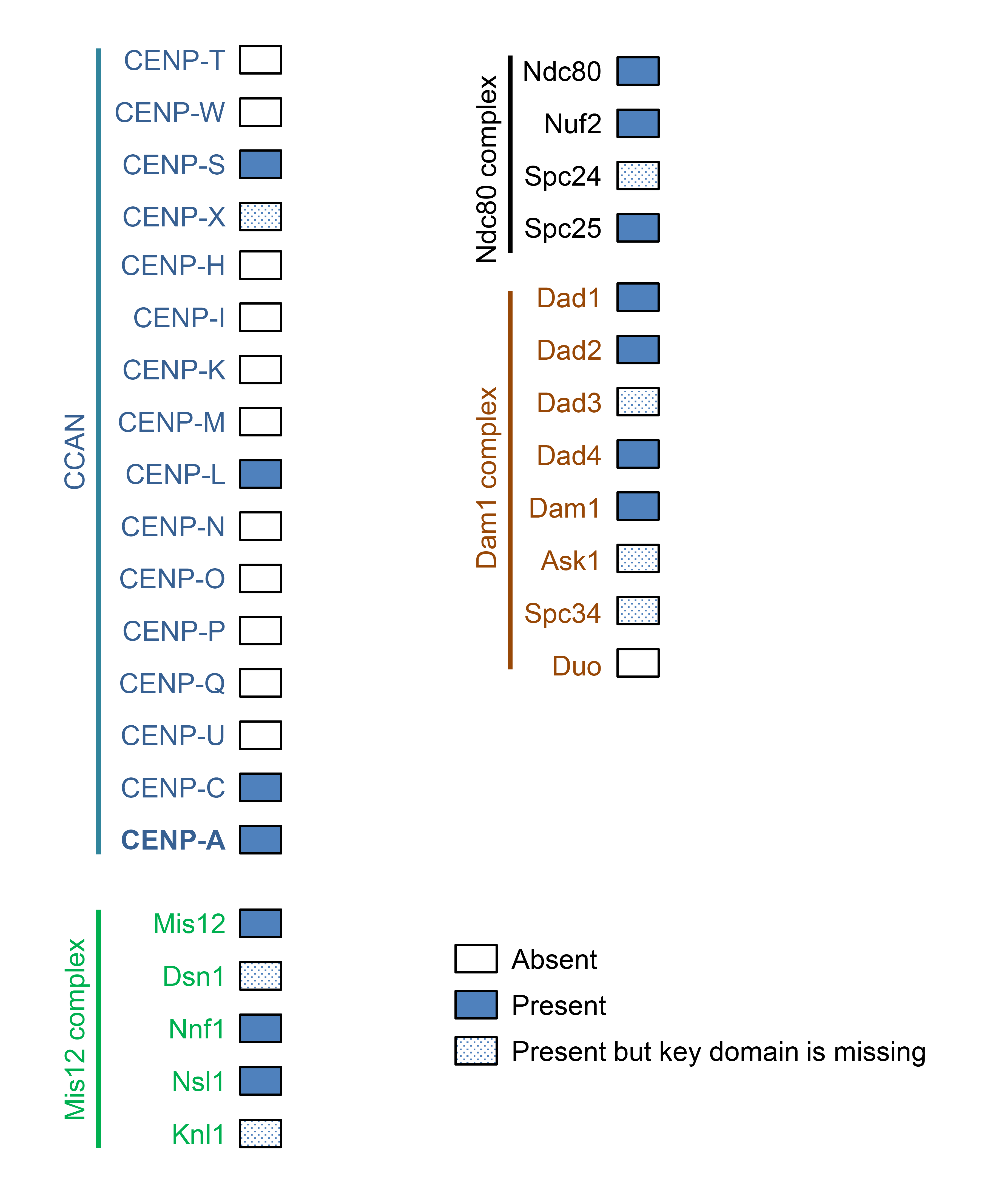

### S2 Fig

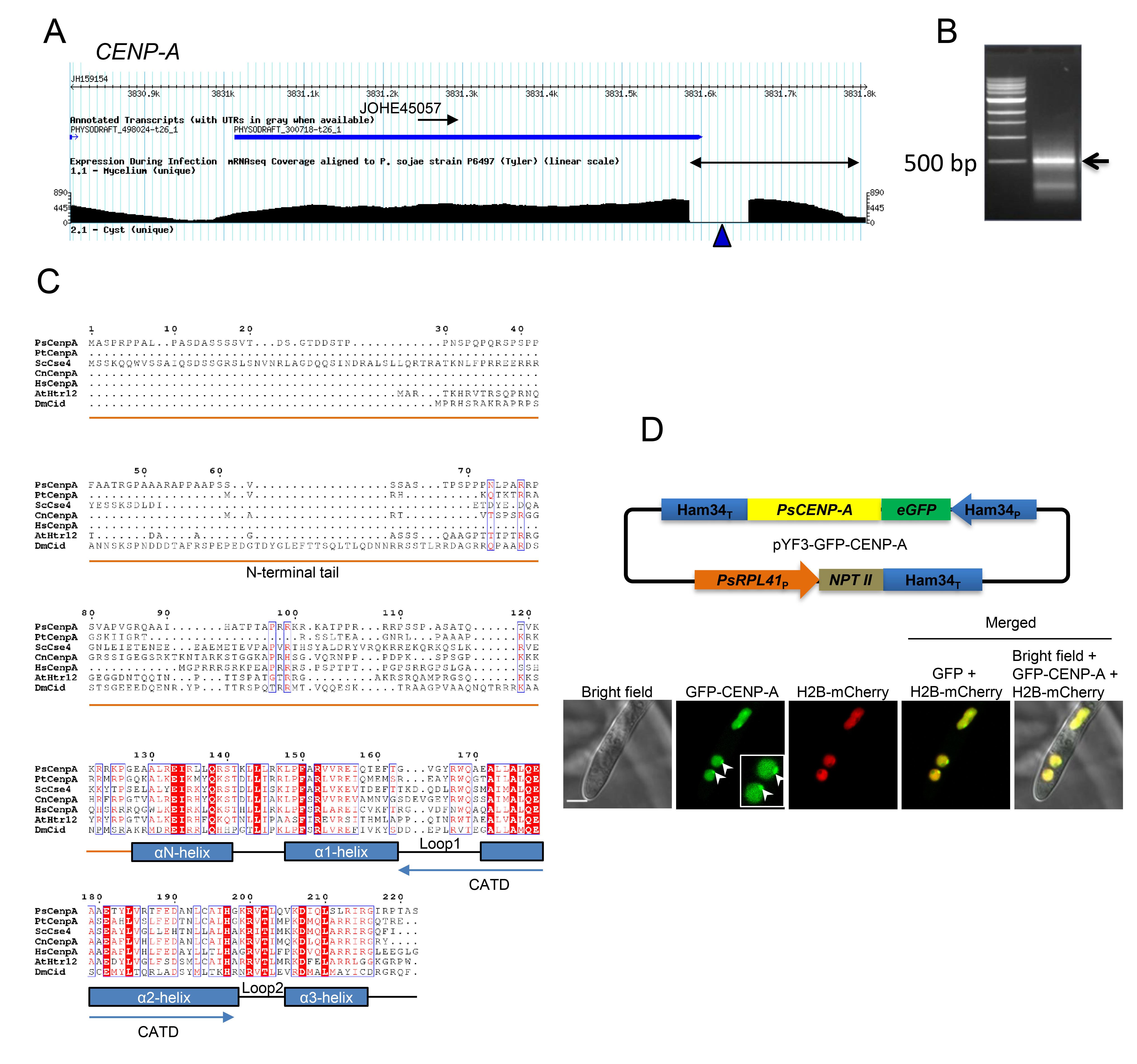

### S3 Fig

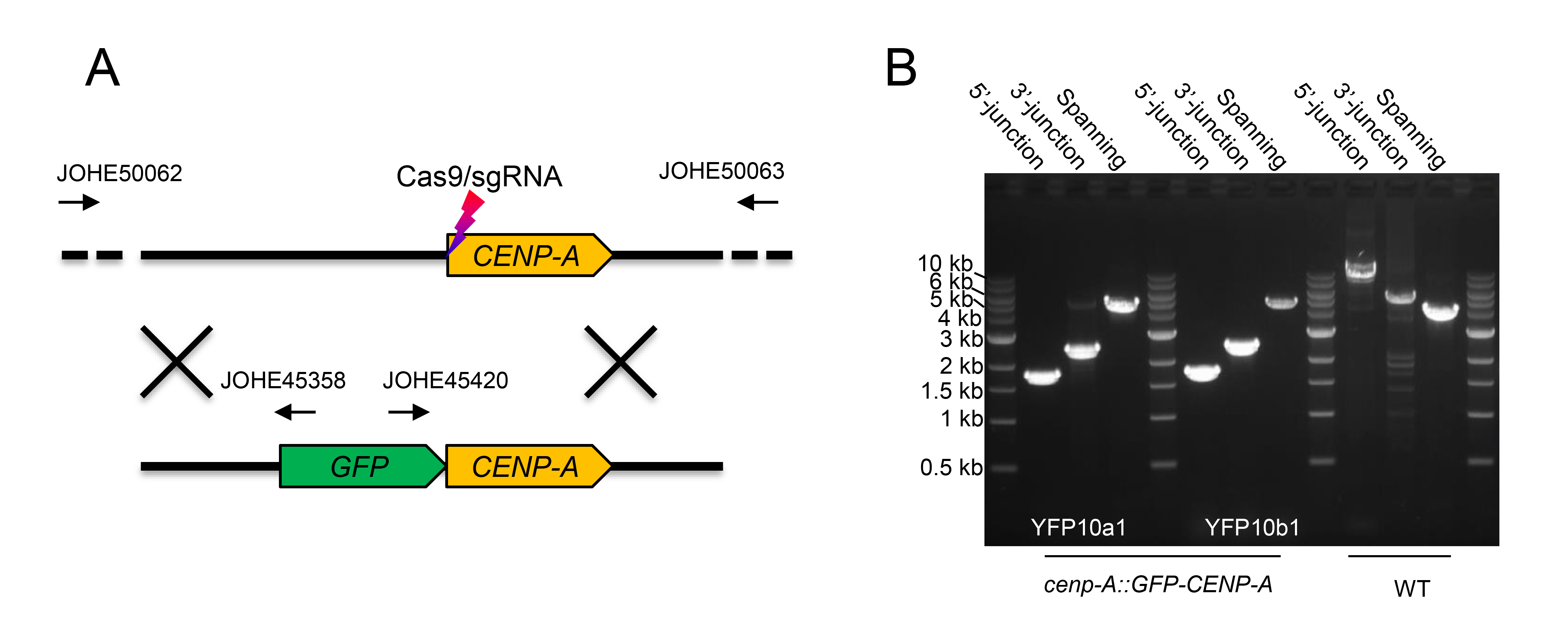

### S4 Fig

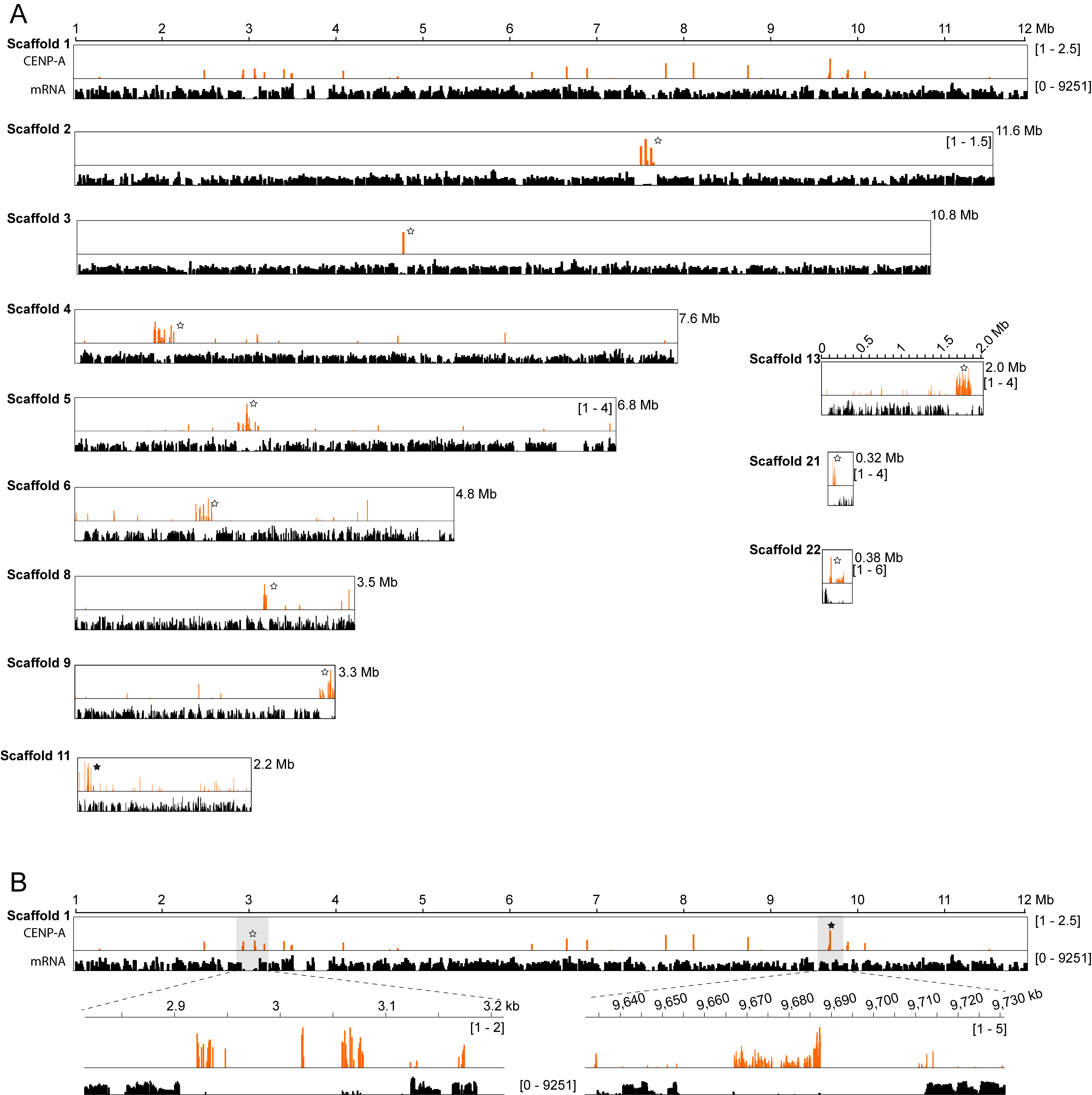

### S5 Fig

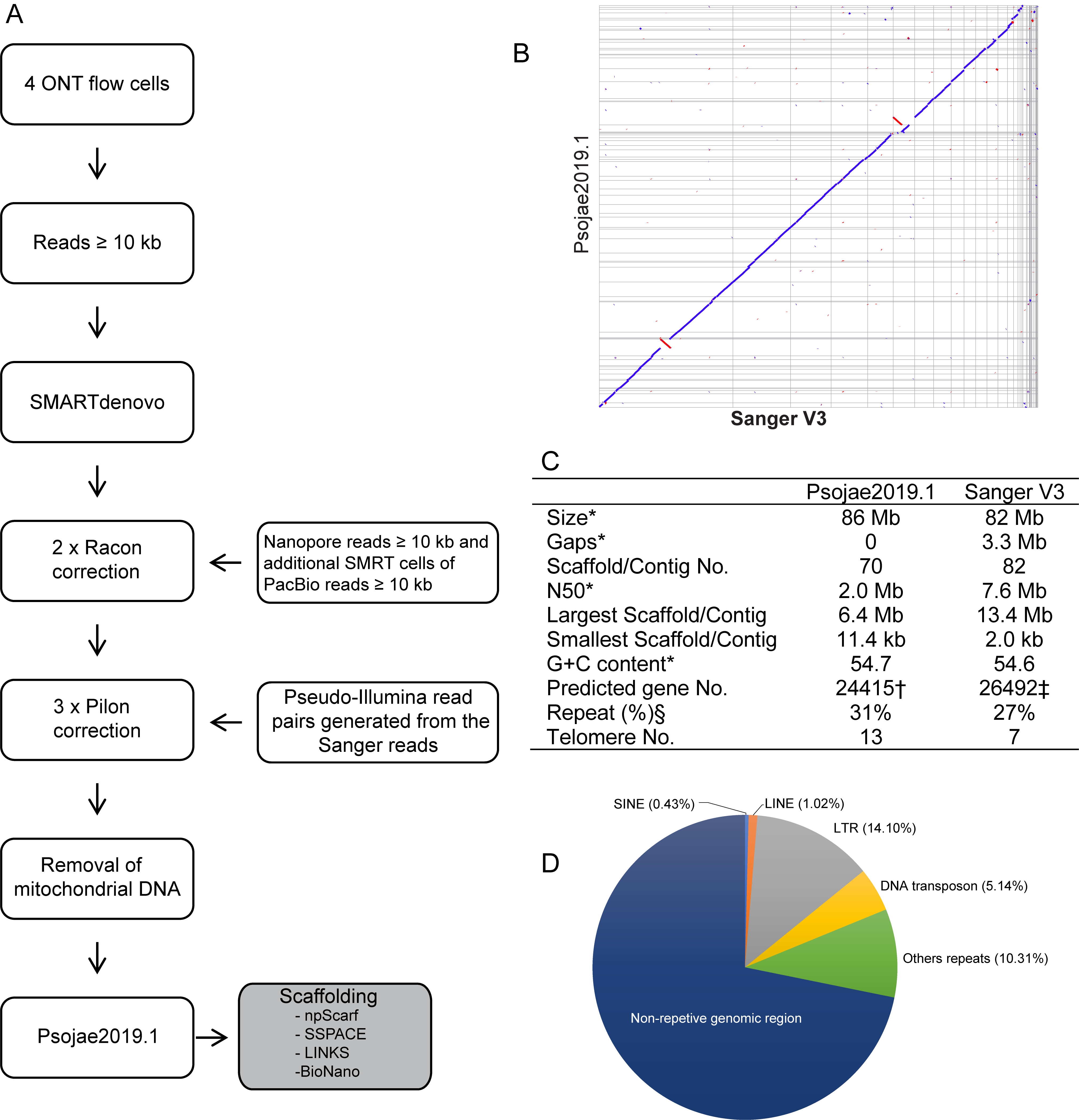

### S6 Fig

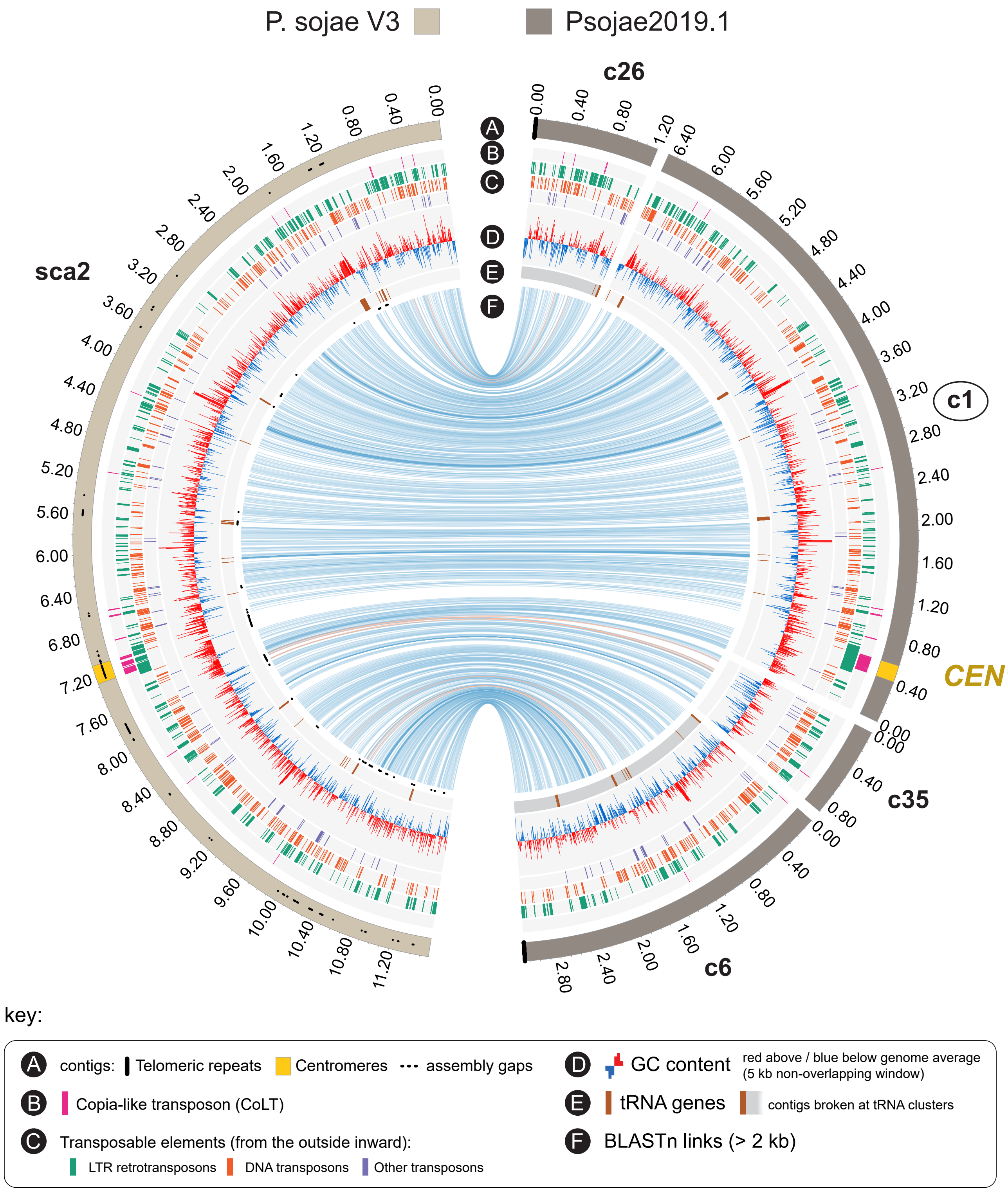

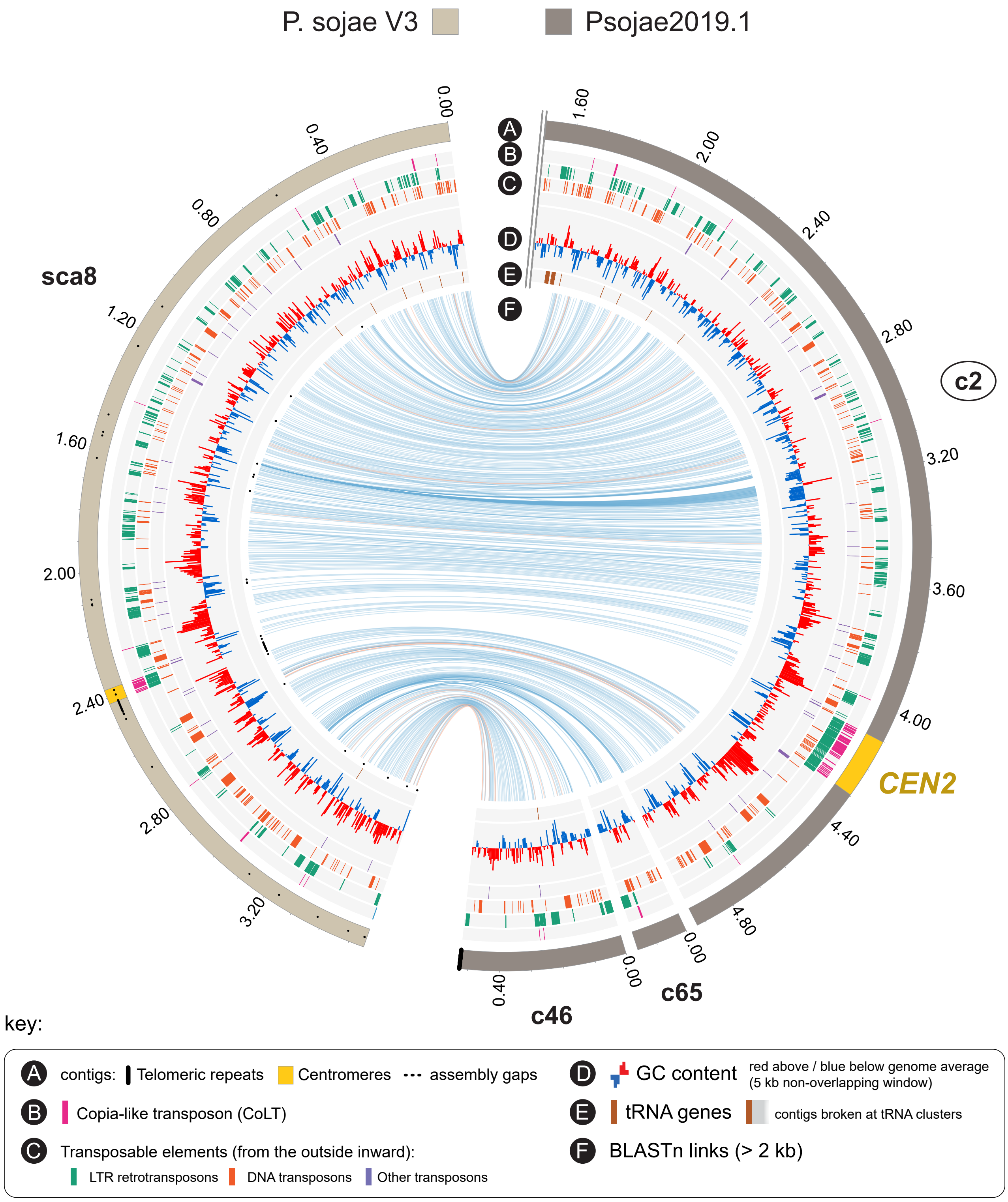

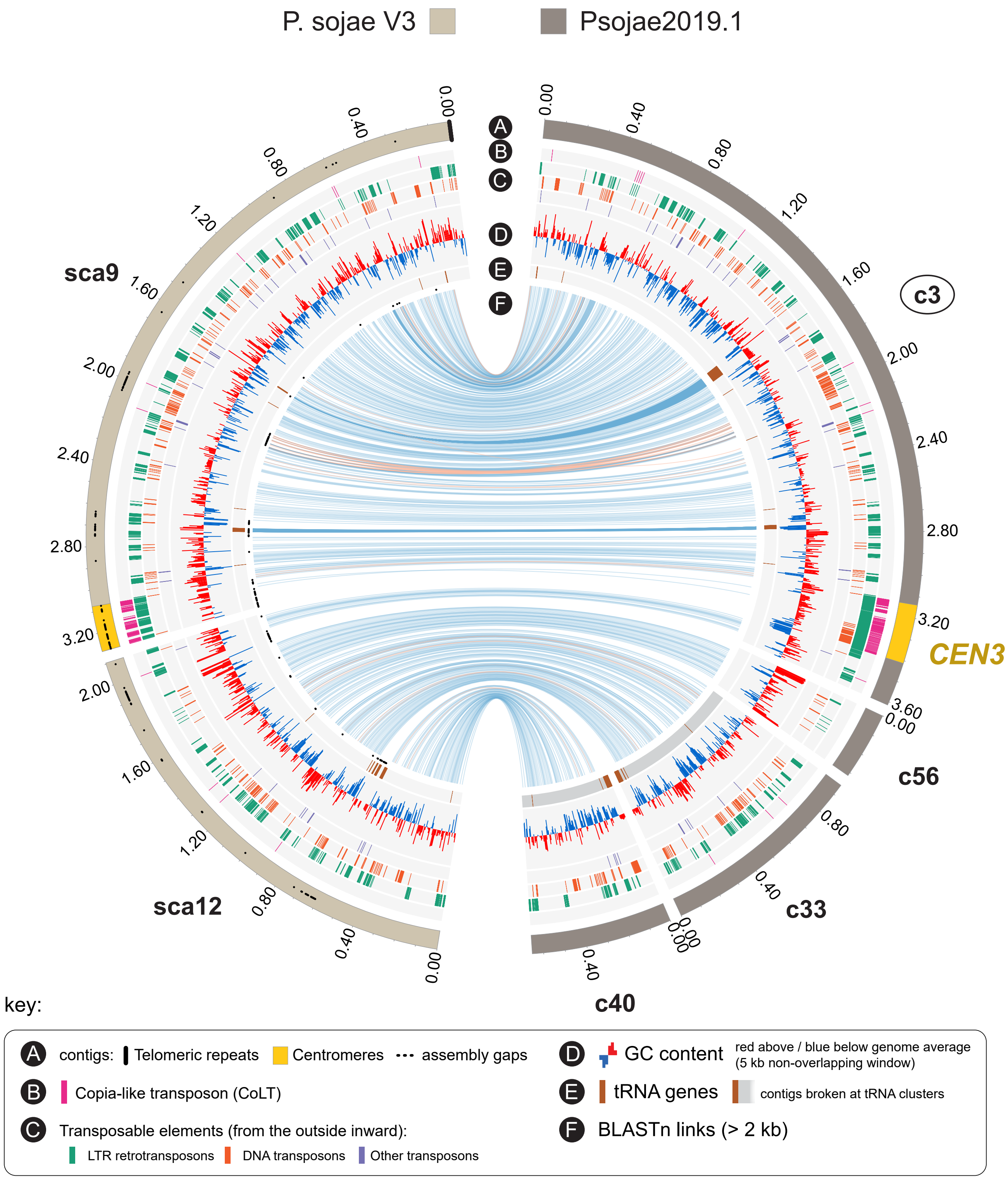

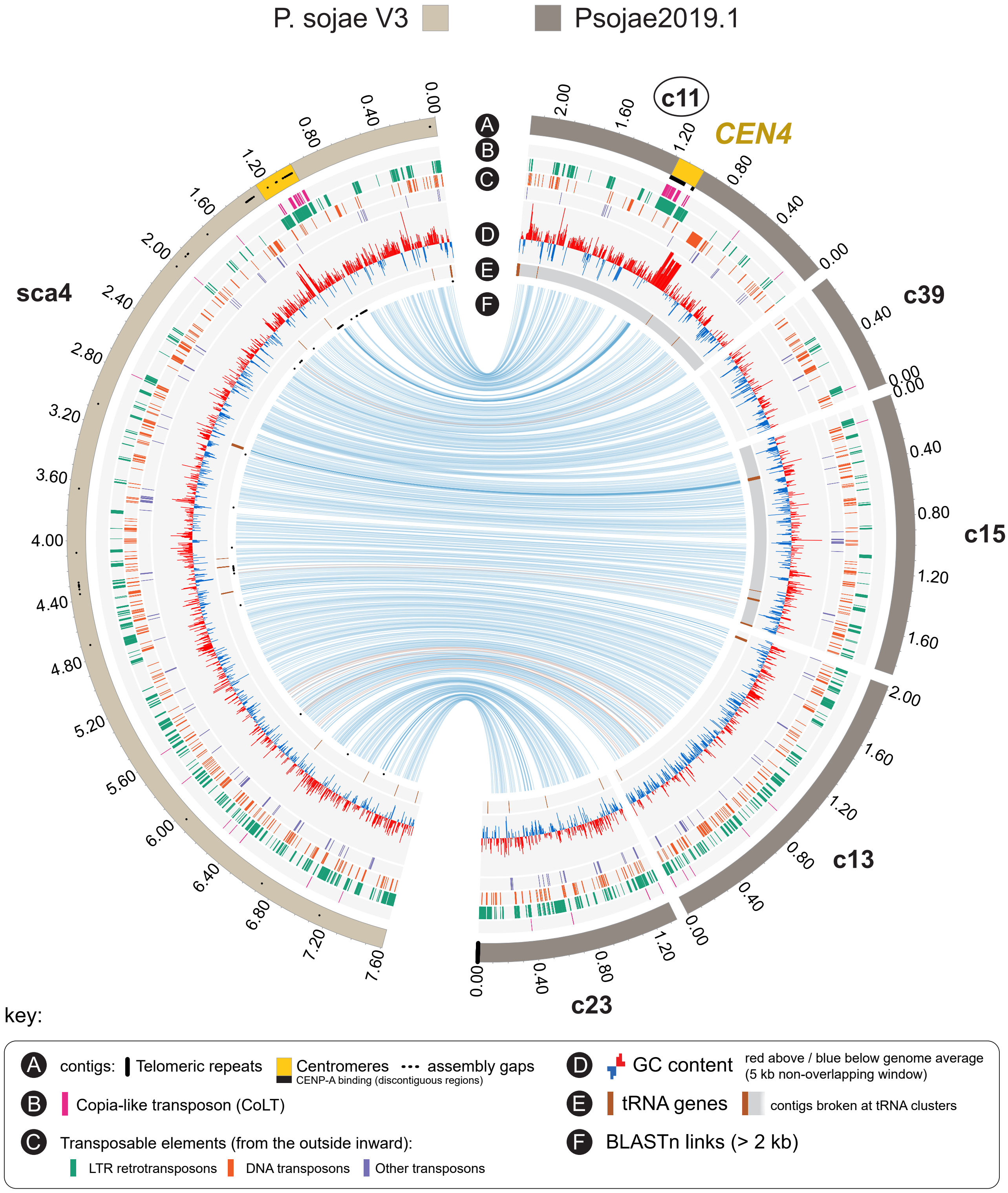

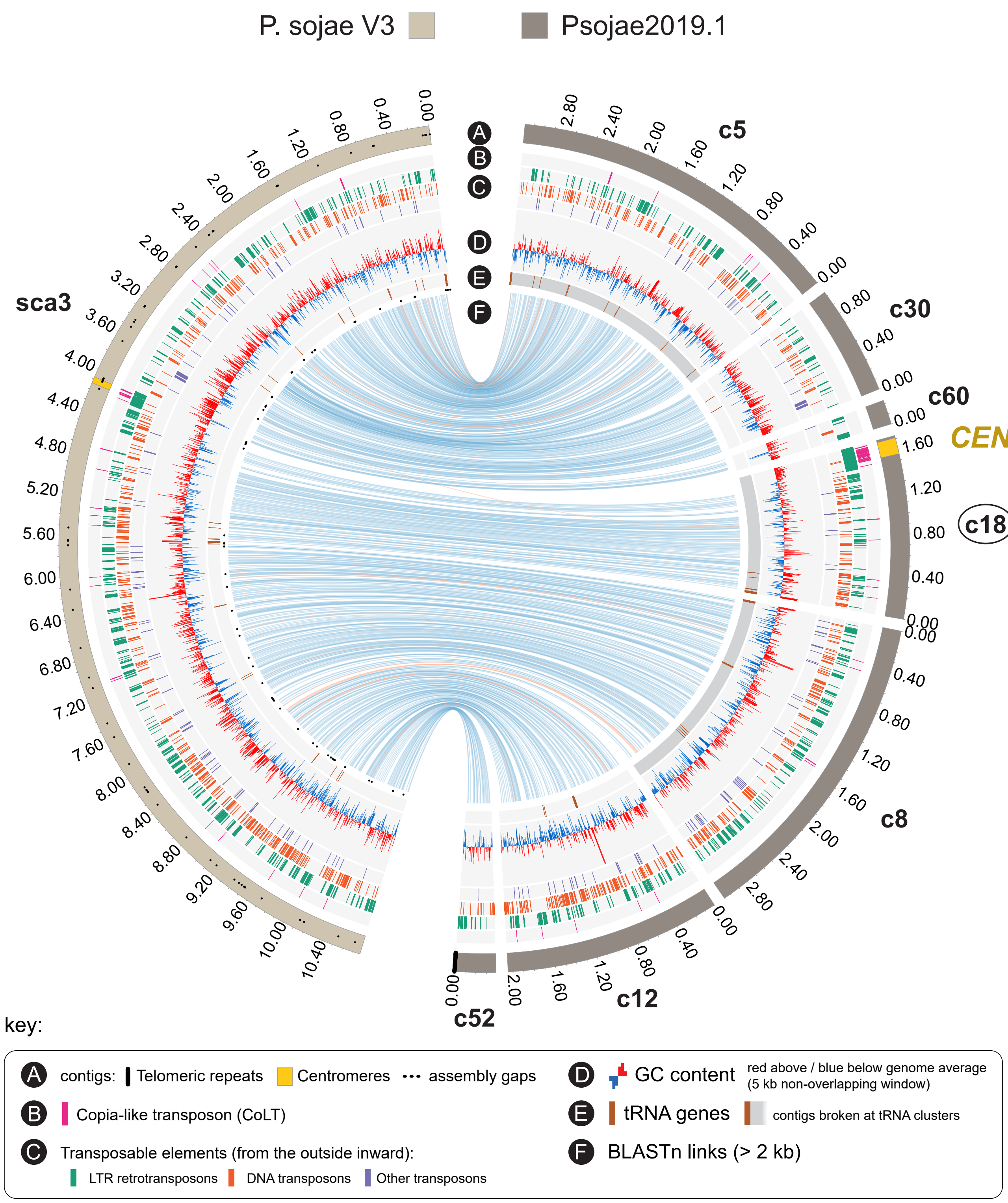

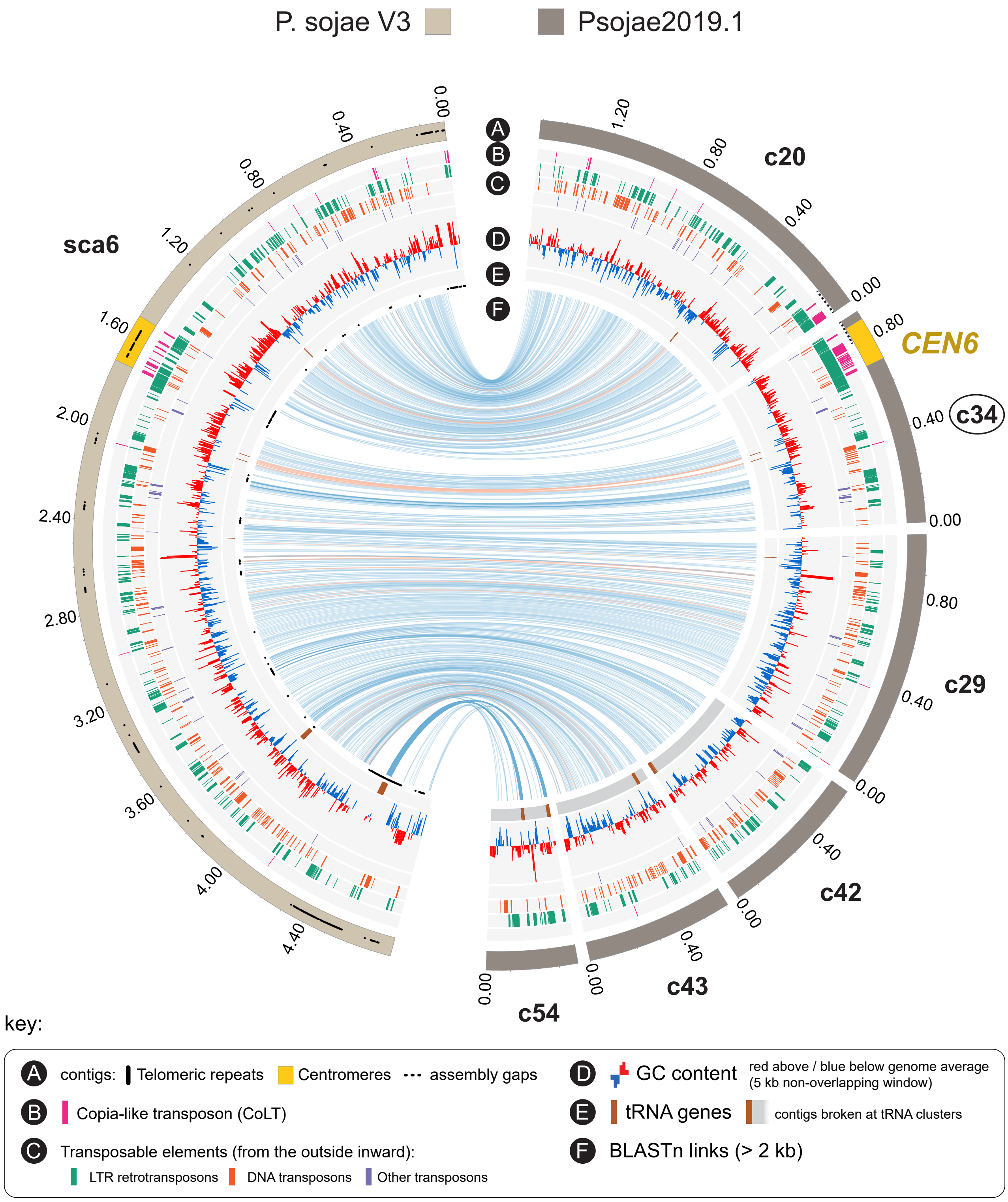

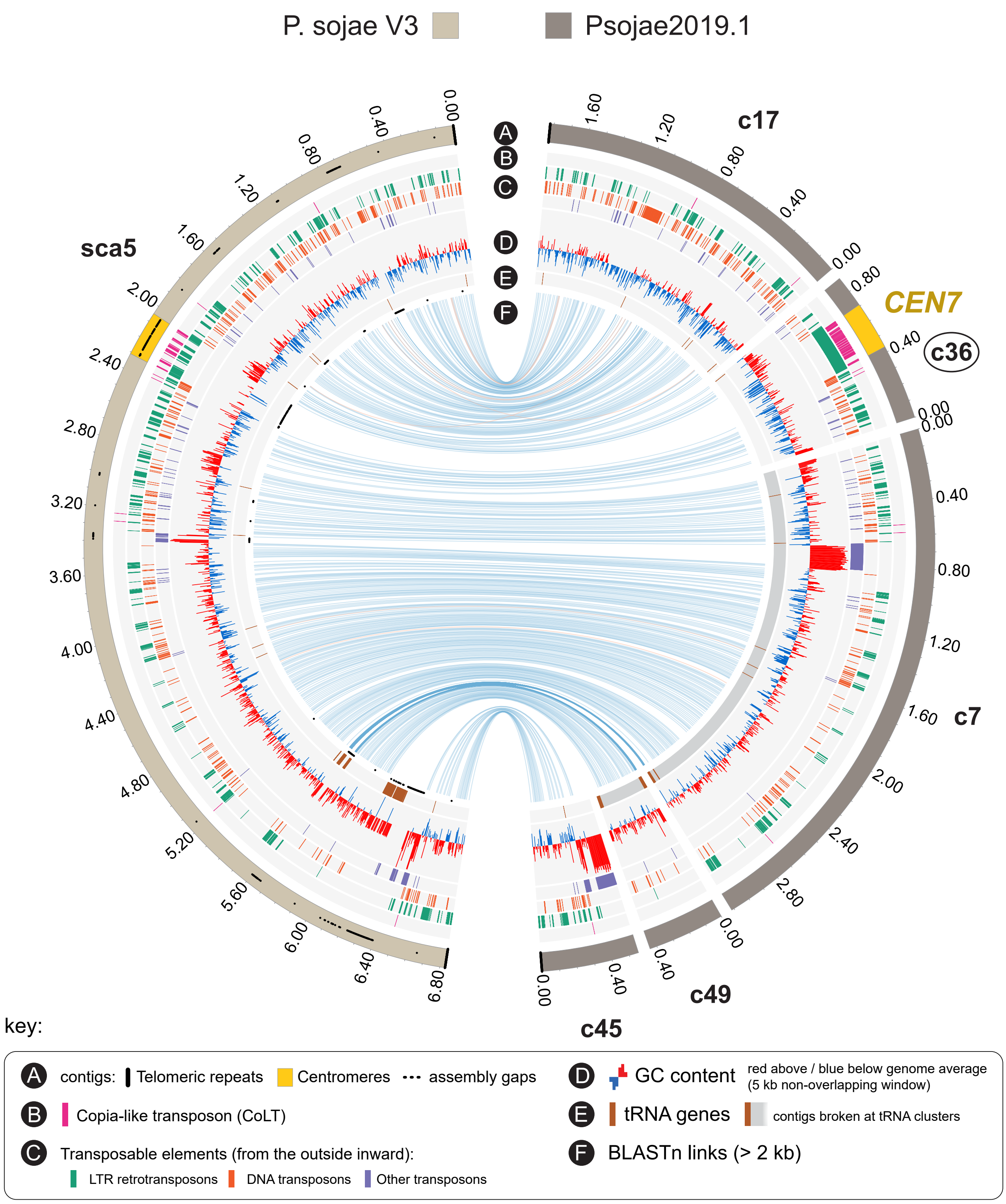

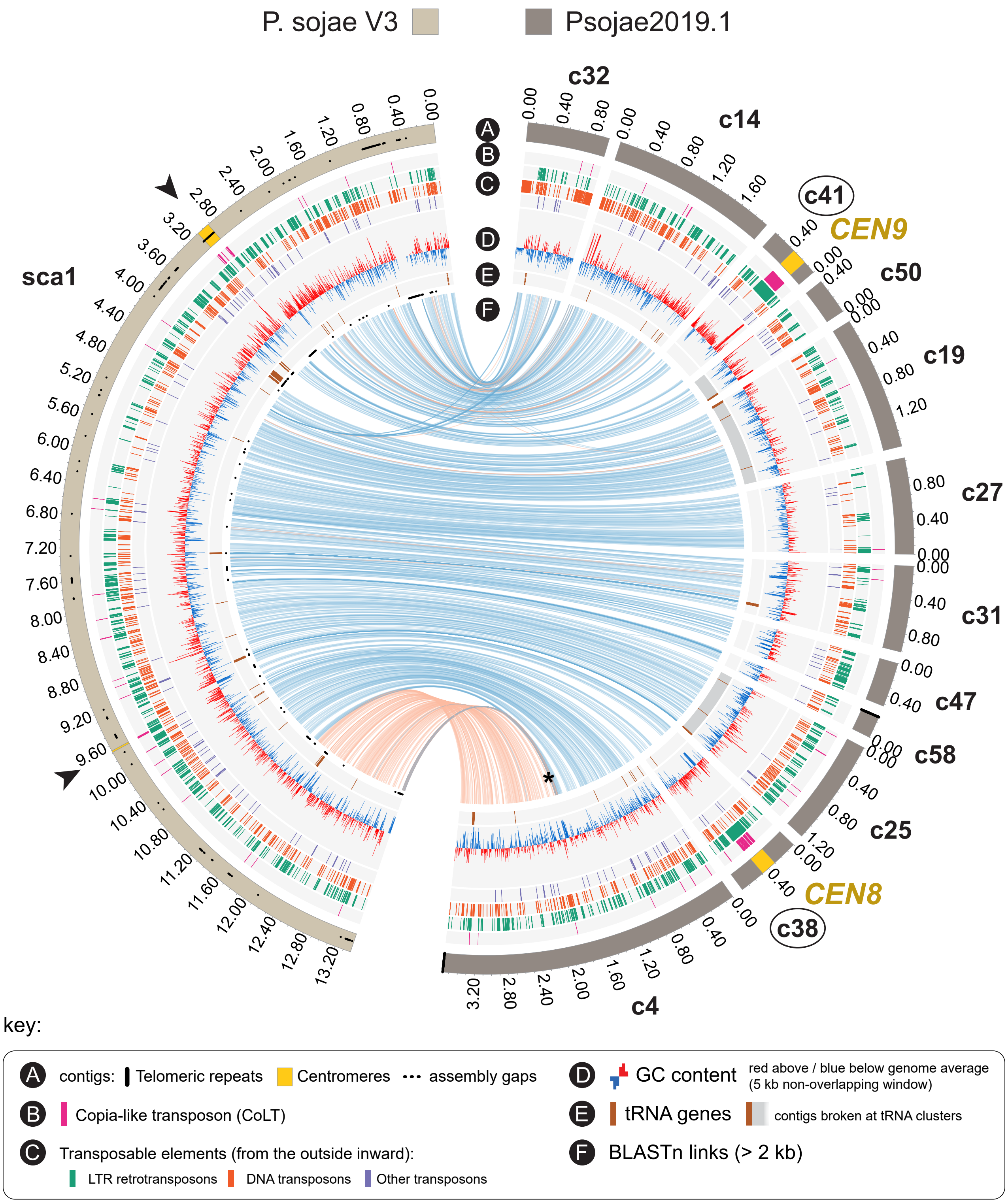

### S7 Fig

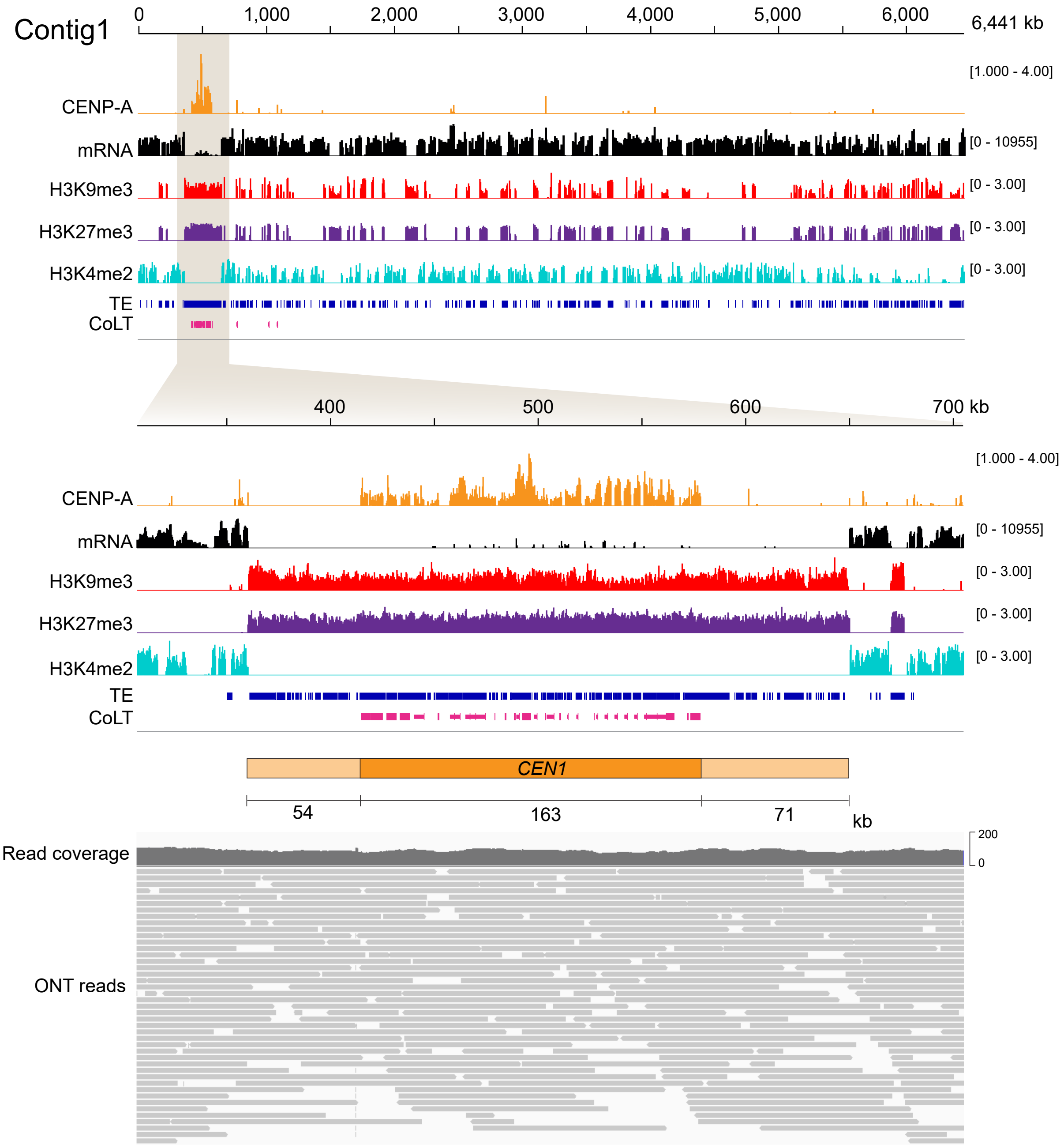

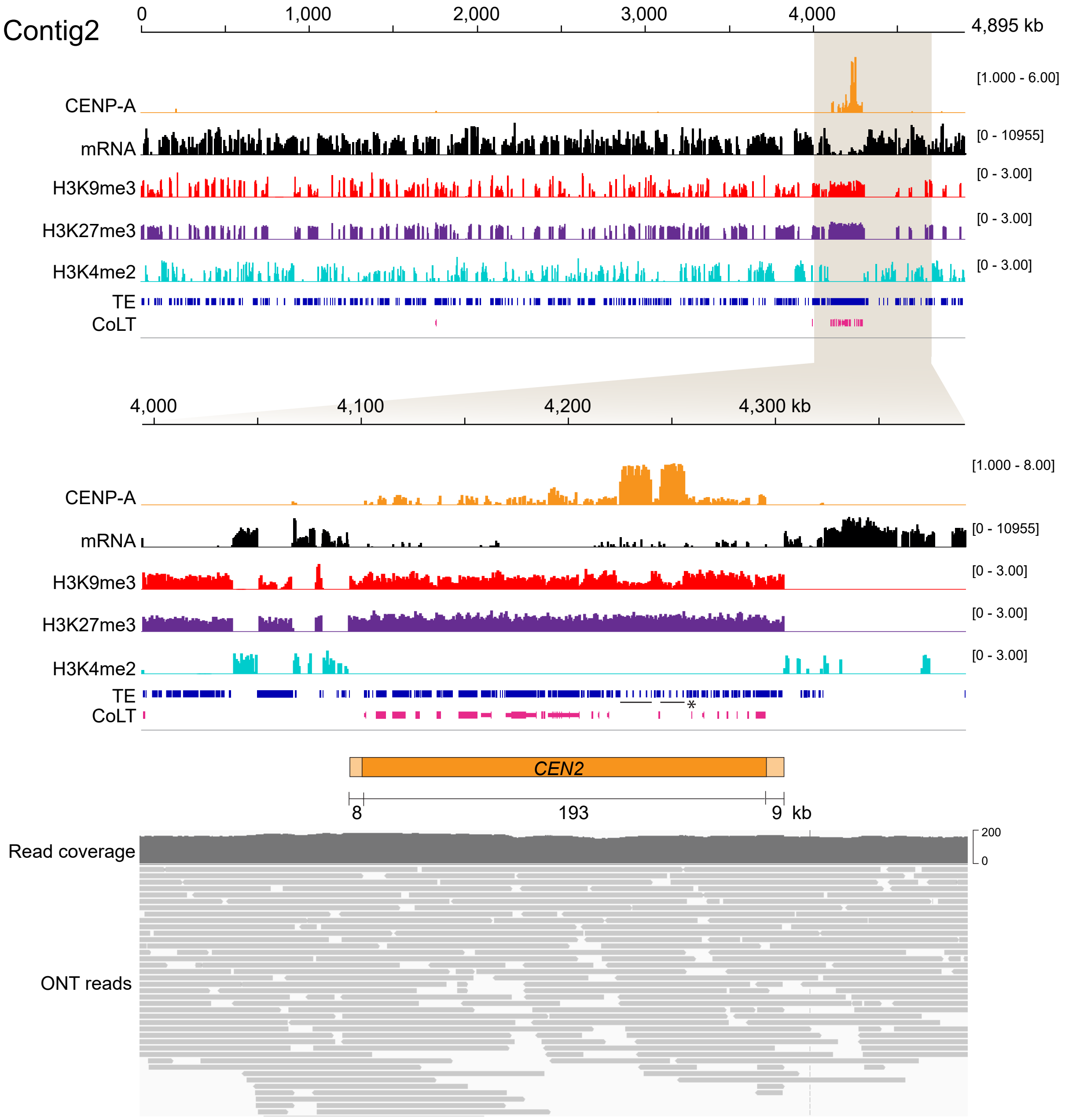

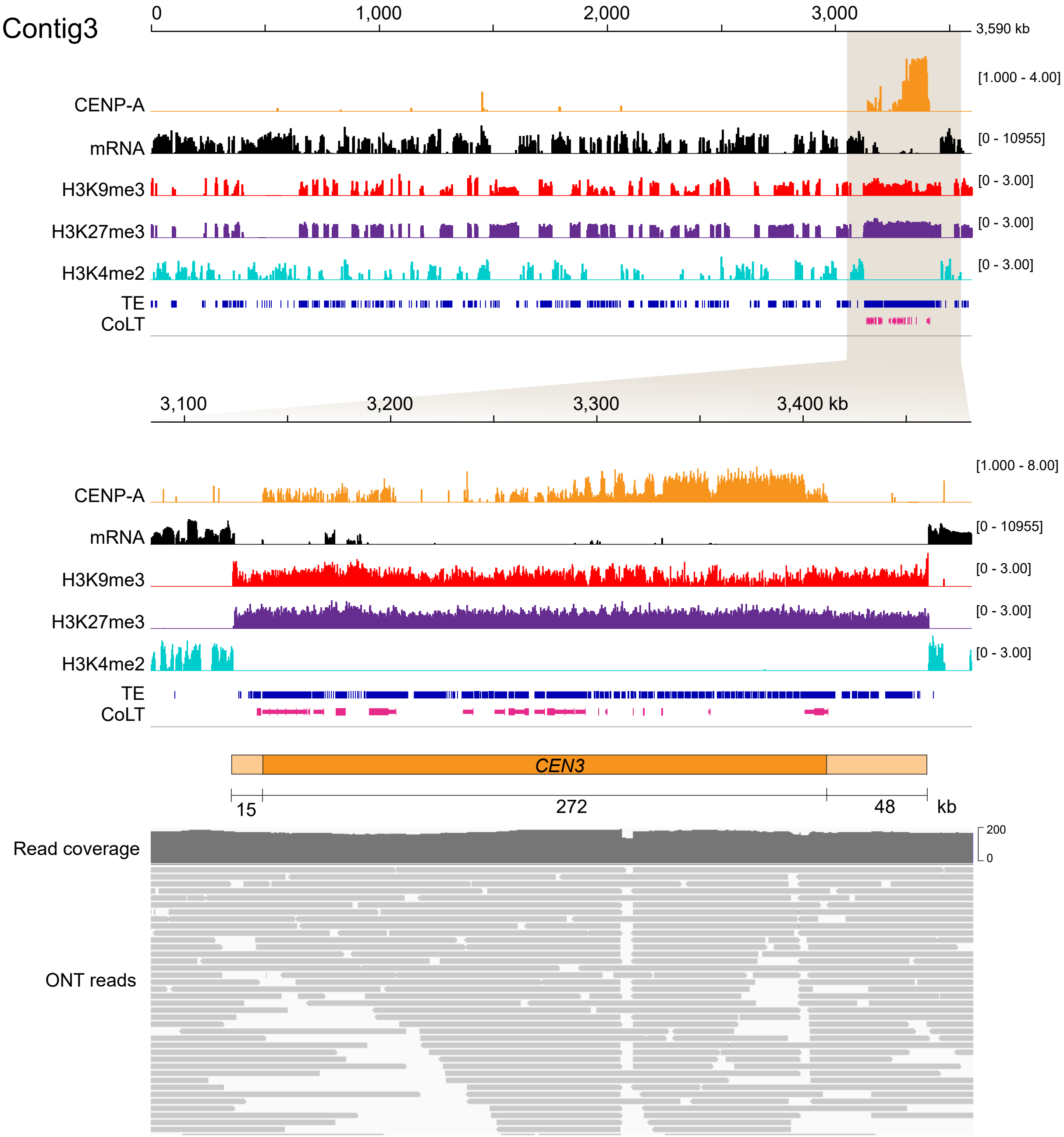

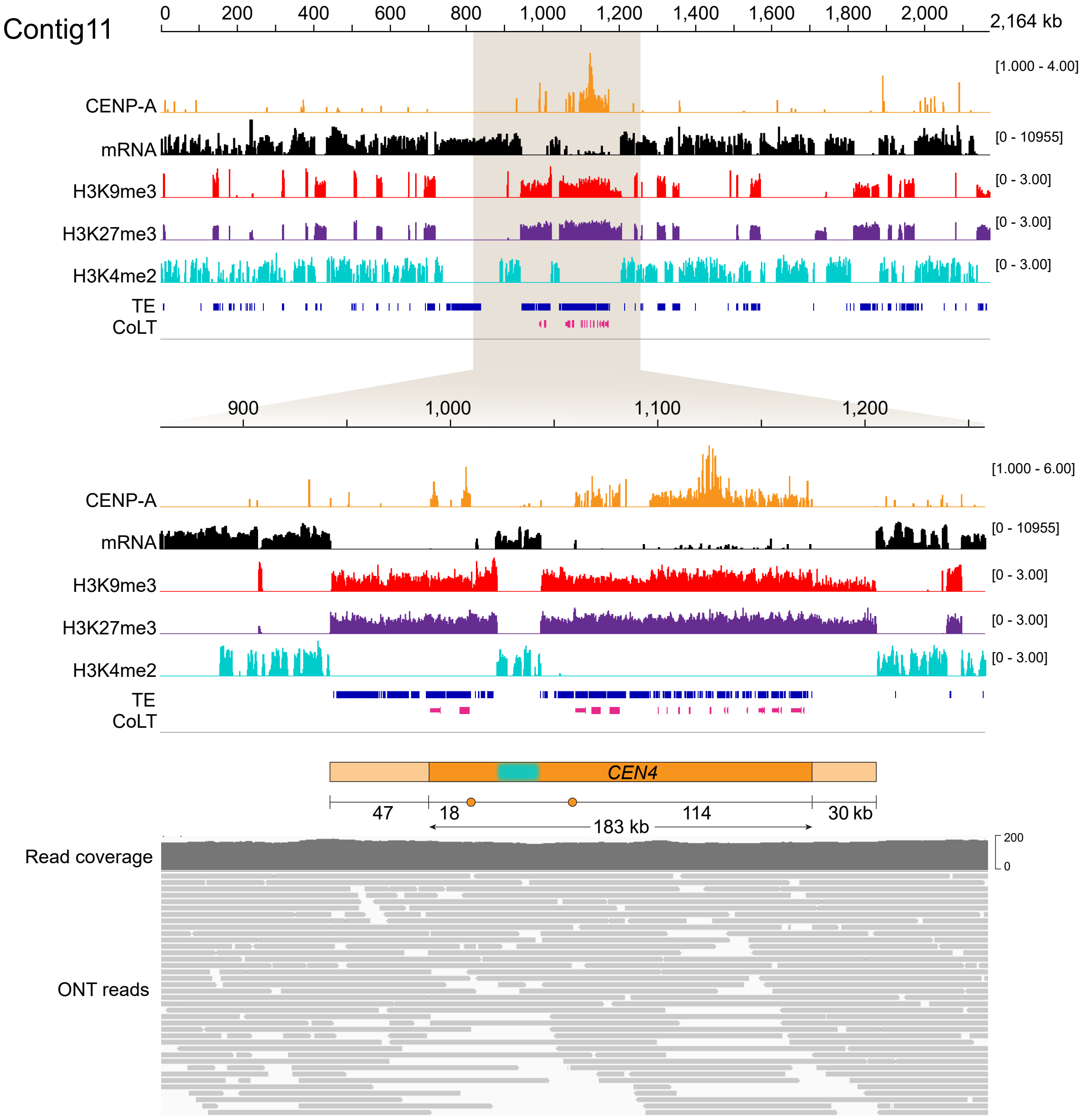

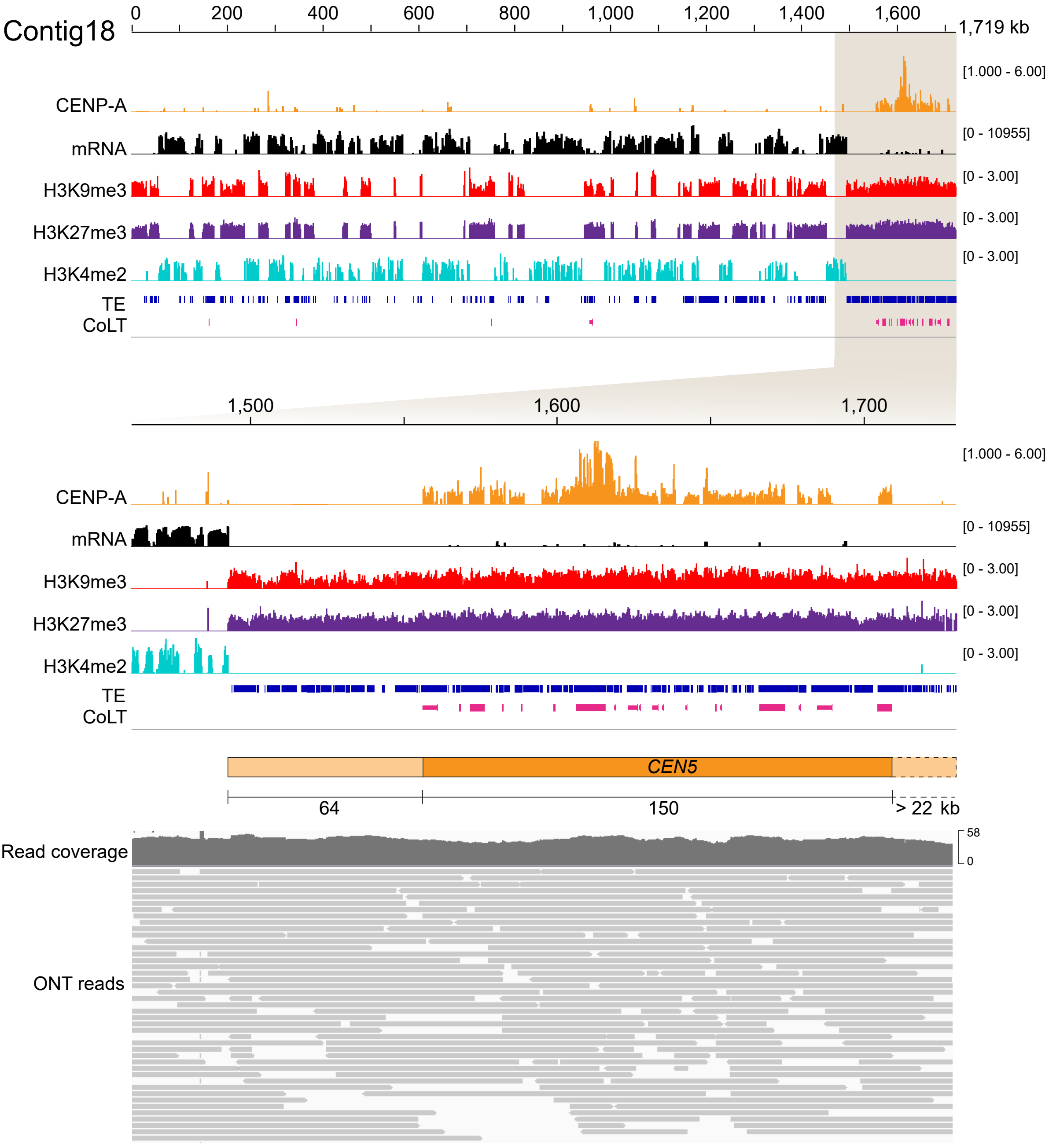

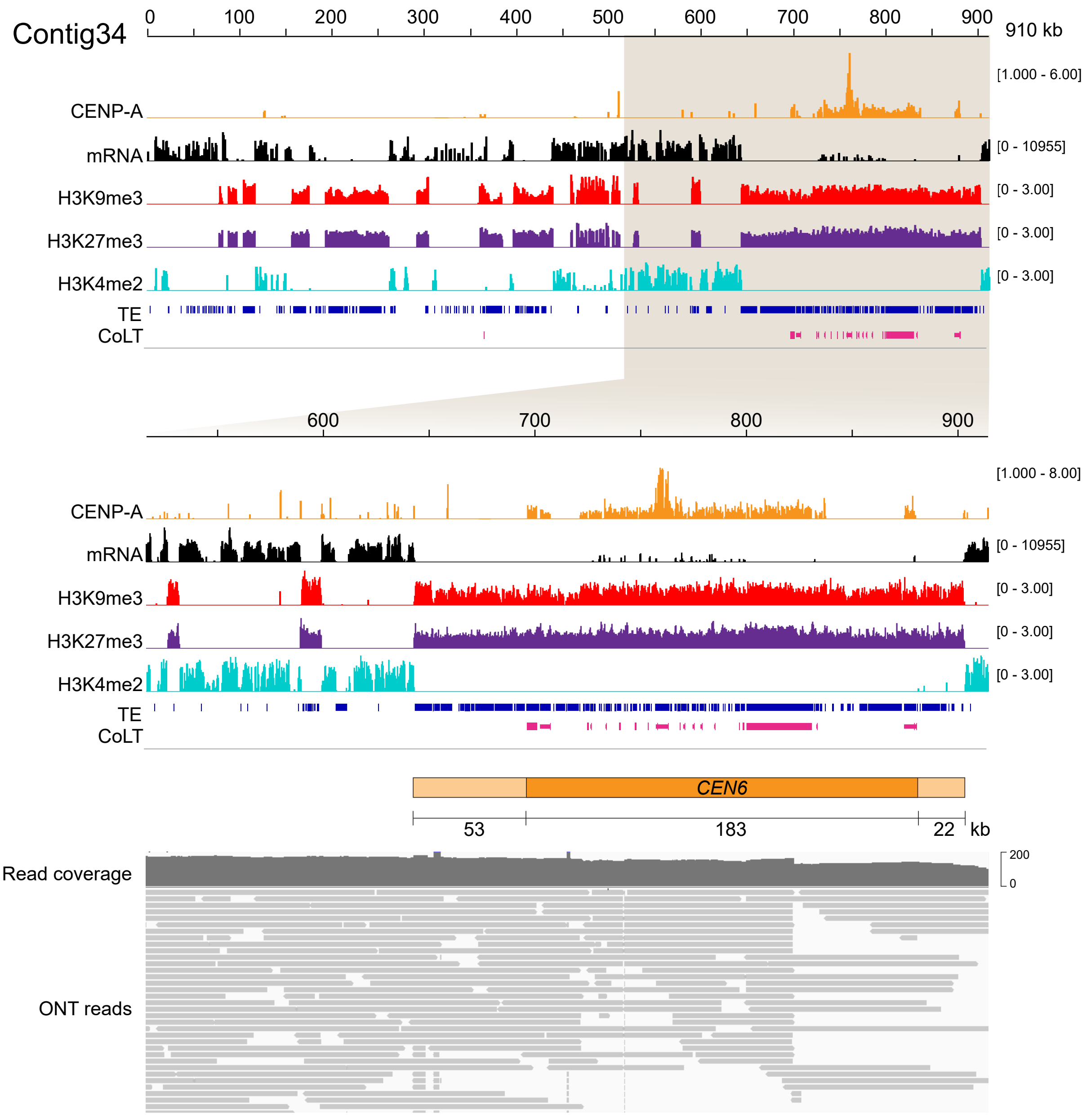

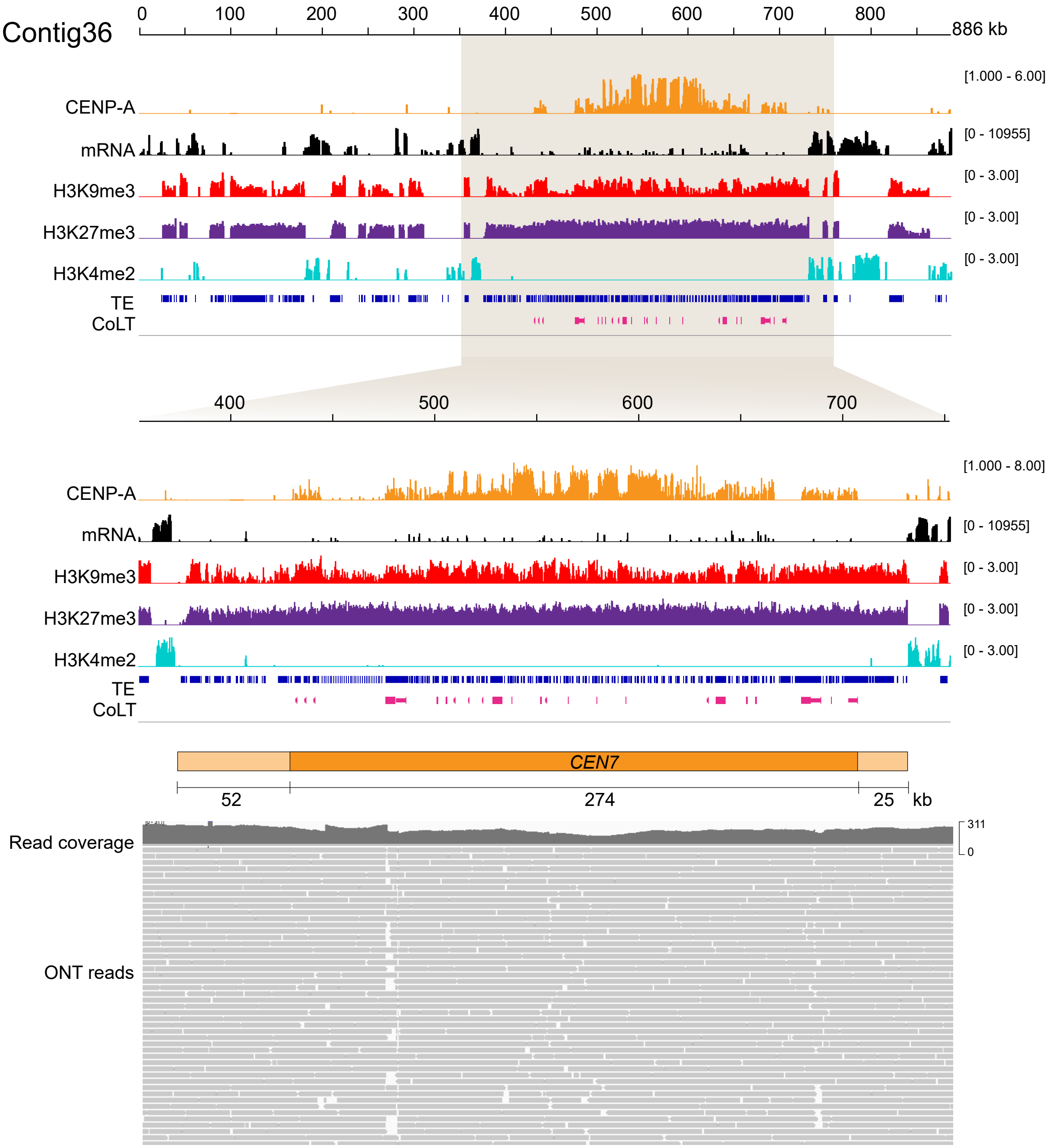

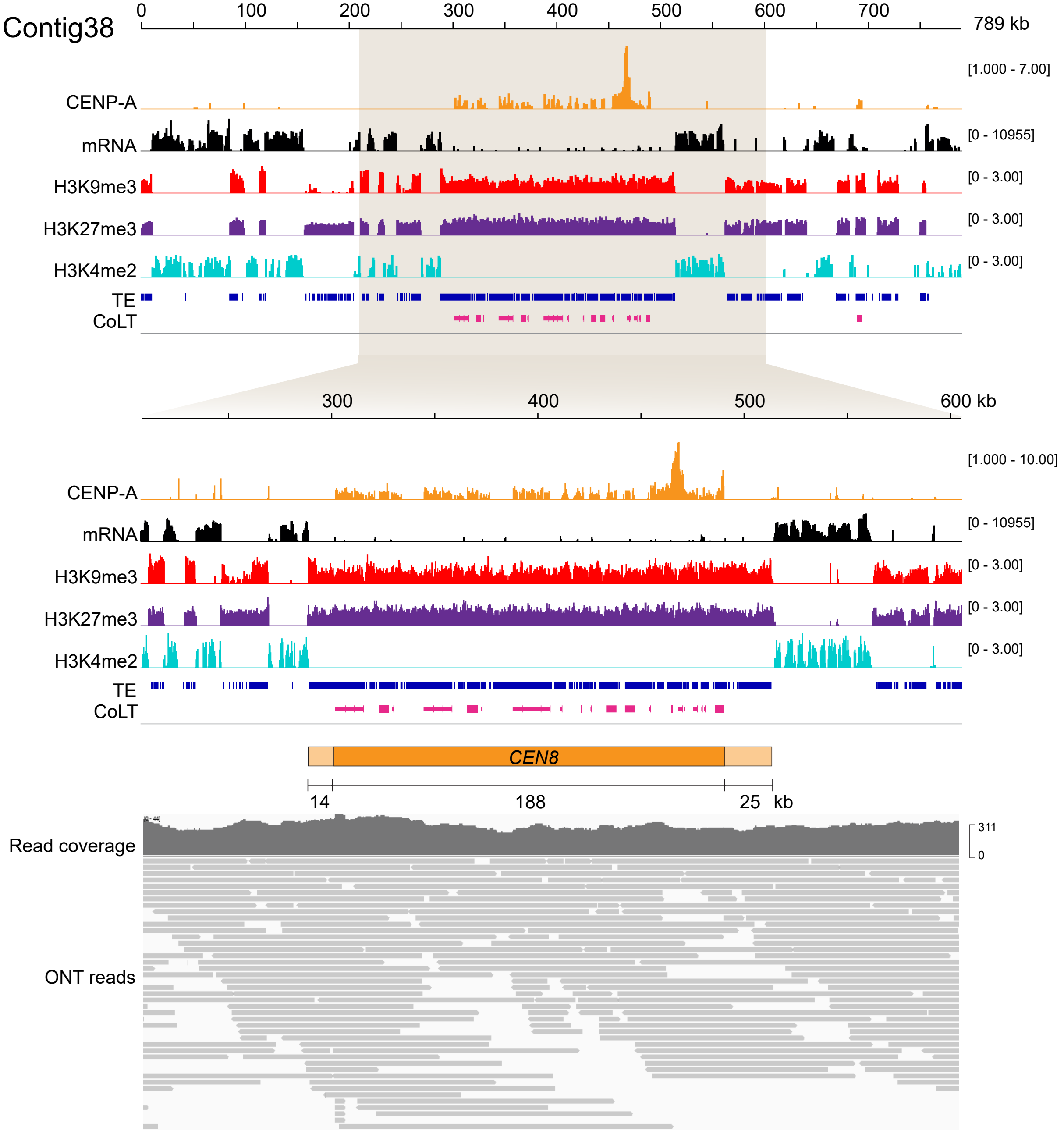

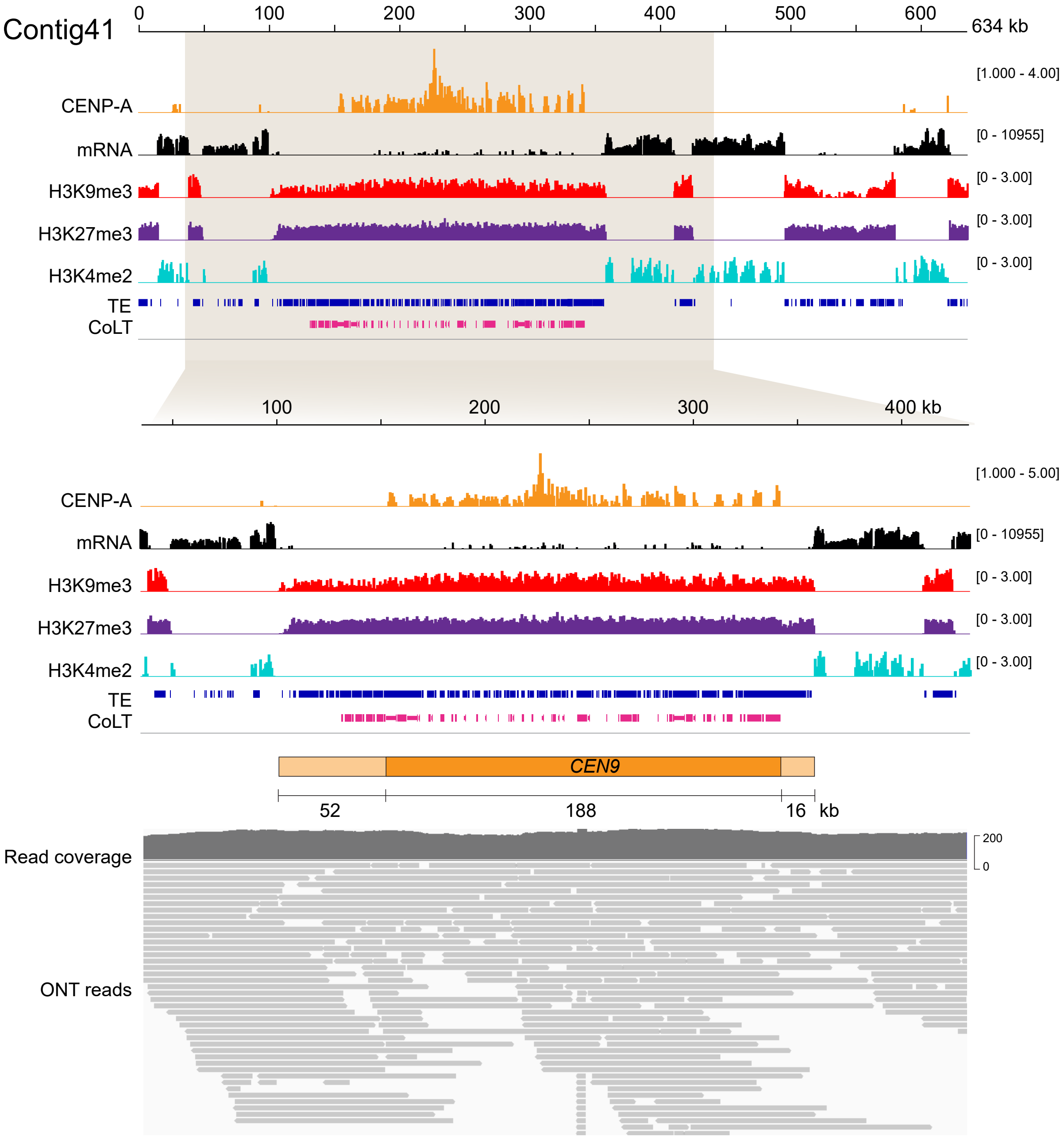

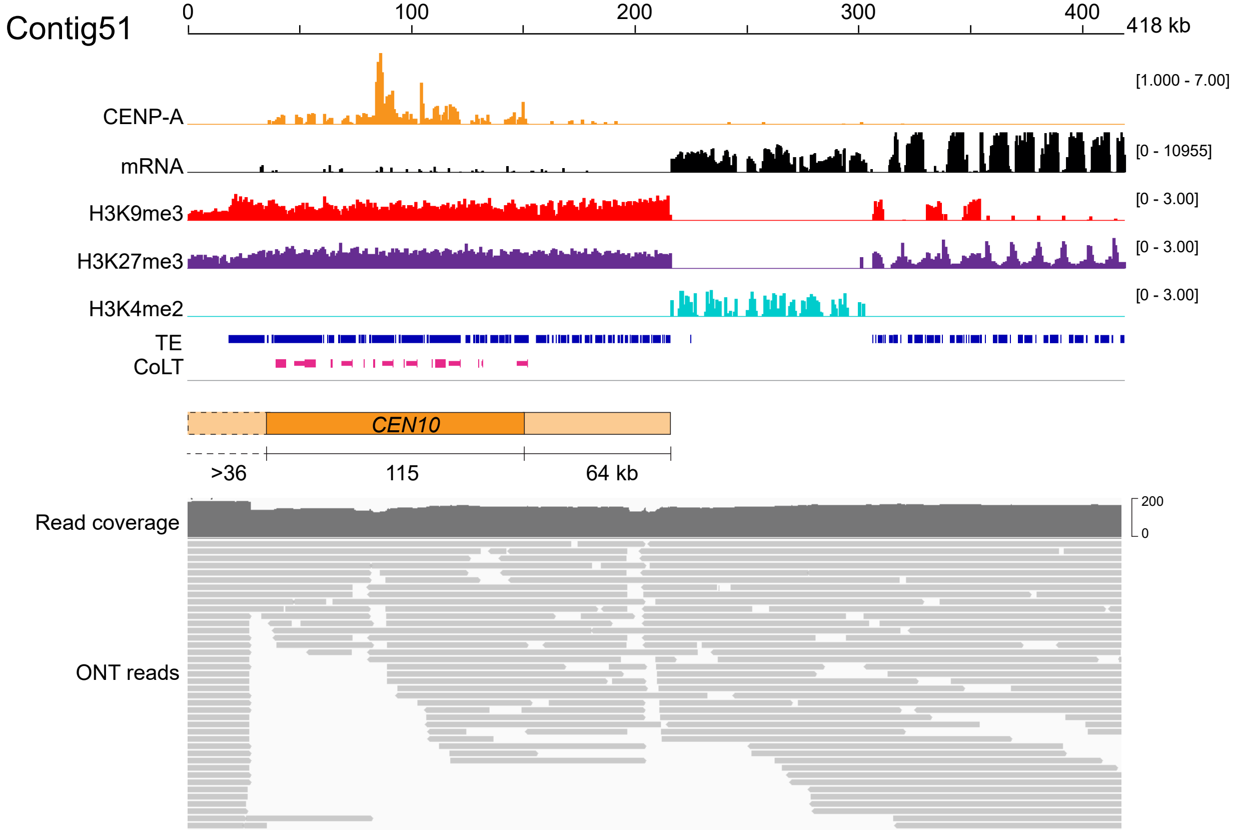

### S8 Fig

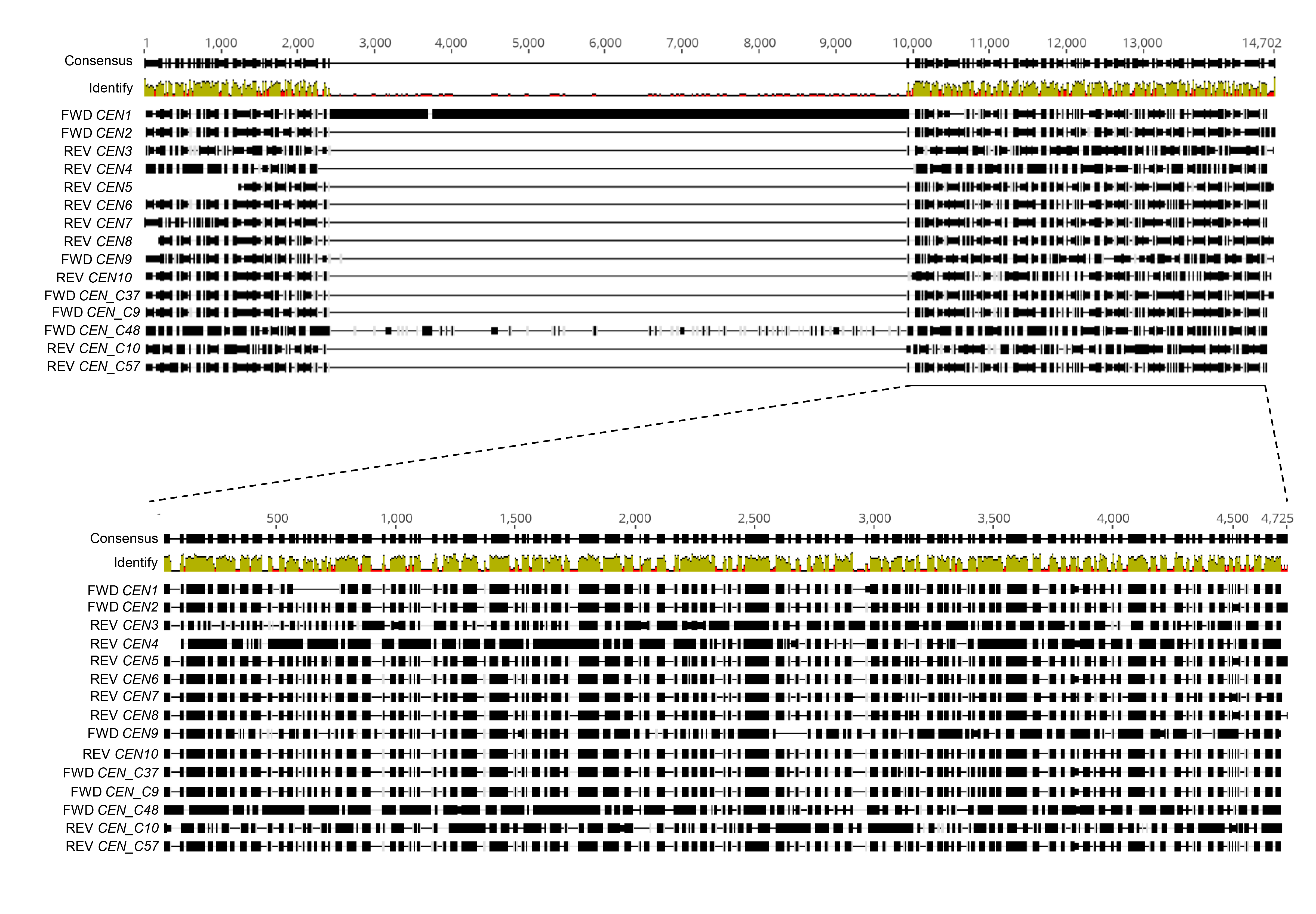

### S9 Fig

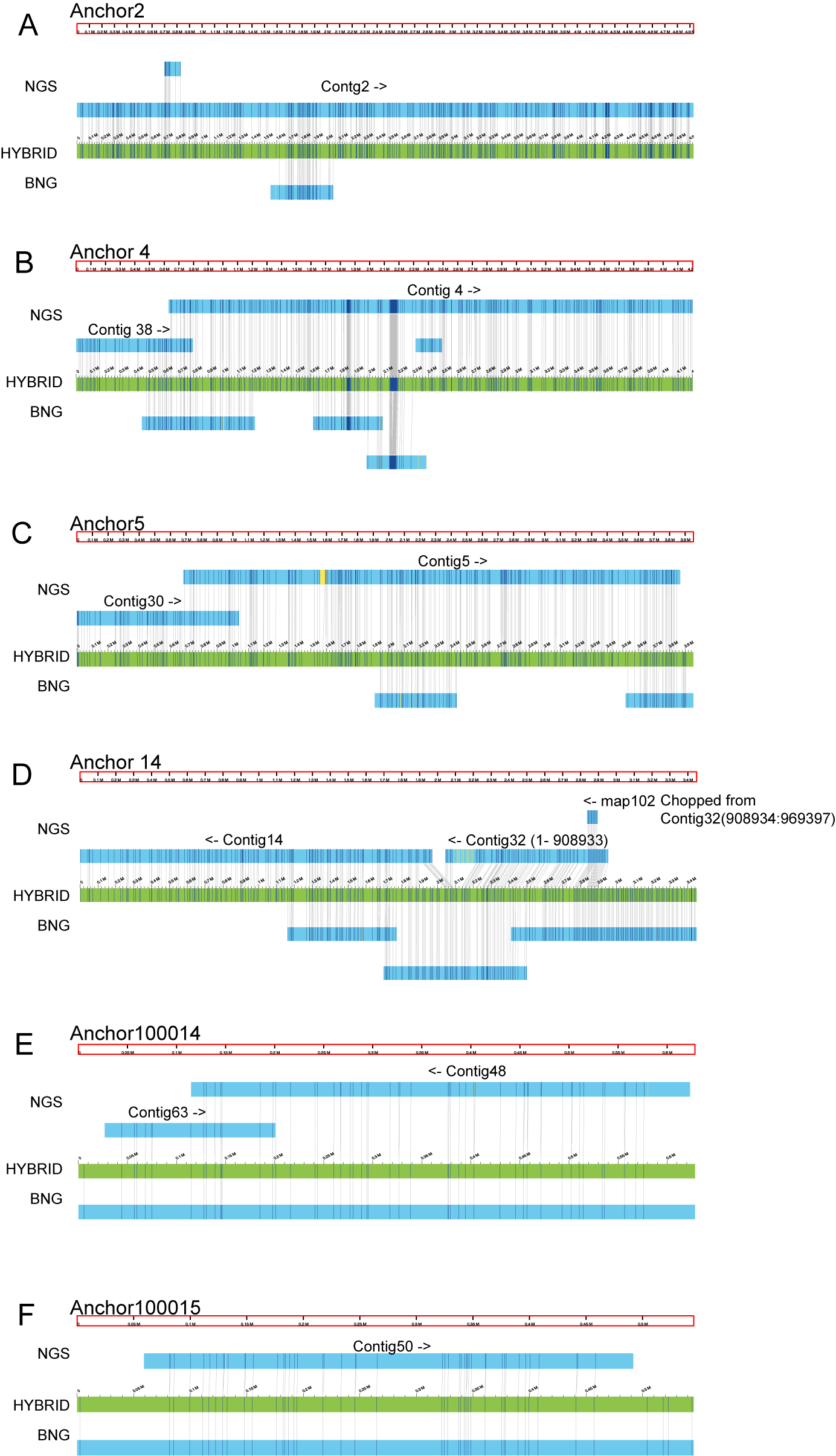

### S10 Fig

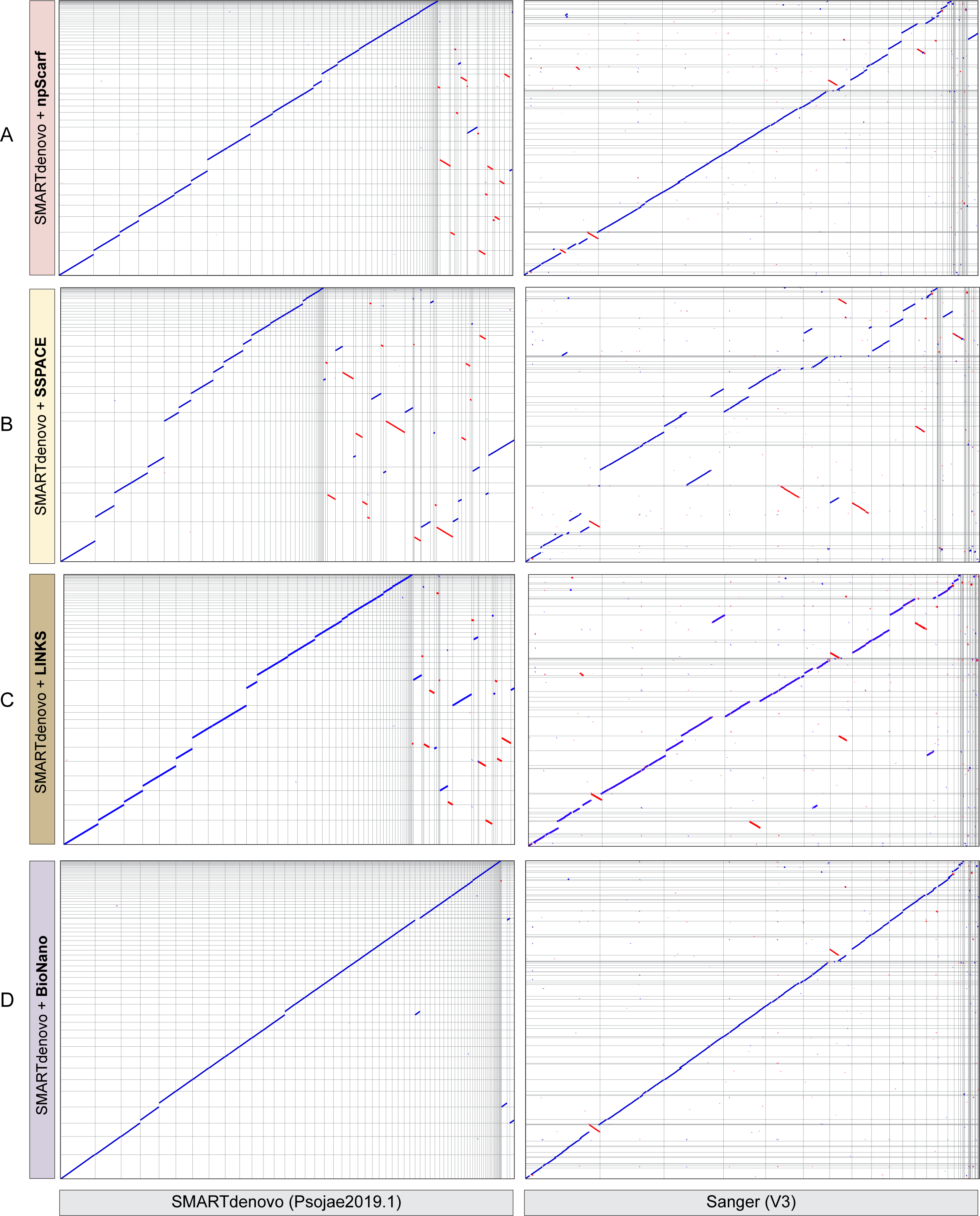
