## Supplementary material for "Long transposon-rich centromeres in an oomycete reveal divergence of centromere features in Stramenopila-Alveolata-Rhizaria lineages": S1 Table

**S1 Table. Metrics of ONT sequencing**

|  |  | Total length (bp) | Coverage (x) * | Read No. | Mean length (bp) | Longest (bp) | N50 | N70 | N90 |
| --- | --- | --- | --- | --- | --- | --- | --- | --- | --- |
| Flow cell 1 (R9.4) | >100 kb | 30,541,569 | 0.3 | 263 | 116,128 | 252,971 | 112,270 | 106,586 | 101,622 |
|  | >50 kb | 327,521,221 | 3.4 | 4,826 | 67,866 |  | 66,604 | 58,878 | 52,691 |
|  | >10 kb | 997,779,784 | 11 | 33,067 | 30,174 |  | 37,700 | 25,875 | 15,354 |
|  | All reads | 1,142,633,179 | 12 | 70,736 | 16,153 |  | 13,134 | 20,497 | 8,277 |
| Flow cell 2 (R9.4.1) | >100 kb | 934,328,836 | 10 | 7,448 | 125,447 | 340,269 | 122,282 | 111,892 | 103,698 |
|  | >50 kb | 3,044,872,424 | 32 | 37,740 | 80,680 |  | 82,584 | 68,058 | 55,763 |
|  | >10 kb | 5,716,680,752 | 60 | 154,400 | 37,025 |  | 53,463 | 33,290 | 16,416 |
|  | All reads | 7,026,276,360 | 74 | 552,717 | 12,712 |  | 41,526 | 19,392 | 4,880 |
| Flow cell 3 (R9.4.1) | >100 kb | 449,302,343 | 5 | 3,866 | 116,219 | 294,442 | 112,436 | 106,506 | 101,956 |
|  | >50 kb | 3,181,936,478 | 33 | 44,058 | 72,222 |  | 72,501 | 62,507 | 53,958 |
|  | >10 kb | 7,528,224,093 | 79 | 237,702 | 31,671 |  | 43,037 | 27,275 | 14,697 |
|  | All reads | 10,233,317,432 | 108 | 1,112,000 | 9,203 |  | 28,740 | 12,126 | 3,539 |
| Flow cell 4 (R9.4.1) | >100 kb | 623,759,690 | 7 | 5,124 | 121,733 | 348,187 | 118,765 | 109,937 | 102,983 |
|  | >50 kb | 2,575,115,320 | 27 | 33,388 | 77,127 |  | 77,908 | 65,606 | 55,020 |
|  | >10 kb | 4,723,908,946 | 50 | 120,100 | 39,333 |  | 54,134 | 36,469 | 18,702 |
|  | All reads | 5,435,861,752 | 57 | 375,655 | 14,470 |  | 47,299 | 27,140 | 7,215 |
| All flow cells | >100 kb | 2,037,932,438 | 22 | 16,701 | 122,025 | 348,187 | 118,361 | 109,628 | 102,852 |
|  | >50 kb | 9,129,445,443 | 96 | 120,012 | 76,071 |  | 76,505 | 64,751 | 54,625 |
|  | >10 kb | 18,966,593,575 | 200 | 545,269 | 34,784 |  | 48,234 | 30,893 | 15,937 |
|  | All reads | 23,838,088,723 | 251 | 2,111,108 | 11,292 |  | 36,856 | 17,302 | 4,296 |

* Coverage was calculated based on the genome size of 95 Mb [1]

1.	Tyler BM, Tripathy S, Zhang X, Dehal P, Jiang RH, Aerts A, et al. *Phytophthora* genome sequences uncover evolutionary origins and mechanisms of pathogenesis. Science. 2006;313(5791):1261-6. Epub 2006/09/02. doi: 10.1126/science.1128796. PubMed PMID: 16946064.
