## Supplementary material for "Long transposon-rich centromeres in an oomycete reveal divergence of centromere features in Stramenopila-Alveolata-Rhizaria lineages": S2 Table

**S2 Table. Statistics of ChIP-seq samples**

| Sample Name | Accession (NCBI) | Total reads | Reads mapped | % |
| --- | --- | --- | --- | --- |
| GFP-CENP-A_ Input | SRX6813696 | 45,803,908 | 38,696,722‬‬‬‬‬‬‬‬‬‬‬‬‬‬ | 84.48 |
| GFP-CENP-A_IP | SRX6813697 | 40,301,524 | 32,785,326‬‬‬‬‬‬‬‬‬‬‬‬‬‬ | 81.35 |
| H3K27_Input | SRX6813696 | 45,803,908 | 38,696,722‬‬‬‬‬‬‬‬‬‬‬‬‬‬ | 84.48 |
| H3K27_IP | SRX6813698 | 30,980,430 | 24,922,553‬‬‬‬‬‬‬‬‬‬‬‬‬‬ | 80.45 |
| H3K9me3_Input | SRR10206162 | 33,032,101 | 32,234,974 | 97.59 |
| H3K9me3_IP | SRR10828930 | 33,510,407 | 33,053,755 | 98.64 |
| H3K4me2_Input | SRR8633151 | 34,292,628 | 33,656,800 | 98.15 |
| H3K4me2_IP | SRR10828931 | 30,278,995 | 29,666,330 | 97.98 |
