## Supplementary material for "Long transposon-rich centromeres in an oomycete reveal divergence of centromere features in Stramenopila-Alveolata-Rhizaria lineages": S3 Table

**S3 Table. Metrics of scaffolded assemblies and their comparison to the Sanger V3 and Psojae2019.1 assembly.**

| Assembly* | Sanger V3 | Psojae2019.1 | npScarf | SSPACE | LINKS | Bionano |
| --- | --- | --- | --- | --- | --- | --- |
| Genome size | 82.60 Mb | 85.88 Mb | 86.38 Mb | 86.56 Mb | 85. 94 Mb | 84.47 Mb |
| Scaffold No. | 82 | 70 (contigs) | 49 | 35 | 45 | 66 |
| Gaps | 3.3 Mb | 0 | 0 | 676,659 bp | 68,409 bp | 65 bp |
| N50 | 7.6 Mb | 2.0 Mb | 3.6 Mb | 5.0 Mb | 4.5 Mb | 2.1 Mb |
| Largest Scaffold | 13.4 Mb | 6.4 Mb | 7.8 Mb | 14.5 Mb | 8.0 Mb | 6.4 Mb |
| Small Scaffold | 2.0 kb | 11.4 kb | 89 kb | 129.1 kb | 126.1 kb | 11.3 kb |
| Telomere No. | 7 | 13 | 13 | 4 | 13 | 13 |

* All *in silico* scaffolded assemblies using the scaffolders npScarf, SSPACE and LINKS, and using BioNano mapping were based on the Psojae2019.1 assembly.
