## Supplementary material for "Long transposon-rich centromeres in an oomycete reveal divergence of centromere features in Stramenopila-Alveolata-Rhizaria lineages": S4 Table

**S4 Table. Five incompletely assembled centromeres in the Psojae2019 assembly and their corresponding CENP-A regions mapped in the Sanger assembly**

| Psojae2019.1 | | | | |  | | Sanger V3 | |
| --- | --- | --- | --- | --- | --- | --- | --- | --- |
| Name | Contig | Position of mapped CENP-A region (kb) | GC% of CEN |  | | Scaffold | | Position of CEN (kb) |
| *CEN_C9** | 9 | 1-158 | 50.76 |  | | 13 | | 1690-1879 |
| *CEN_C48** | 48 | 1-35 | 45.25 |  | |  |  |  |
| *CEN_C10* | 10 | 2248-2312 | 57.23 |  | | 11 | | 118 -143 |
| *CEN_C37* | 37 | 721-771 | 56.26 |  | | 21 | | 61-103 |
| *CEN_C57* | 57 | 229-273 | 56.51 |  | | 14 | | 1716-1806† |

**CEN_C9* and *CEN_C48*, Sequences surrounding the two incompletely assembled centromeres were colinear with Sanger Scaffold13, indicating that *CEN_C9* and *CEN_C48* may be parts of the same centromere.

†The disrupted centromere in Sanger Scaffold 14 was identified by synteny analysis. The start of the centromere position (i.e. 1716 kb) is a transcriptional region that is syntenic to the one next to *CEN_C57*.
