## Supplementary material for "Long transposon-rich centromeres in an oomycete reveal divergence of centromere features in Stramenopila-Alveolata-Rhizaria lineages": S5 Table

**S5 Table. *P. sojae* strains used in the study**

| Strain Name | Genotype | Source |
| --- | --- | --- |
| P6497 | wild type |  |
| YFP09 | *P. sojae* P6497::pYF3-GFP-CENP-A | This study |
| YFP10a1 | *P. sojae* P6497 *cenpa*Δ::*GFP-CENP-A* | This study |
| YFP10b1 | *P. sojae* P6497 *cenpa*Δ::*GFP-CENP-A* | This study |
