## Supplementary material for "Long transposon-rich centromeres in an oomycete reveal divergence of centromere features in Stramenopila-Alveolata-Rhizaria lineages": S6 Table

**S6 Table. Primers used in this study.**

| Name | Sequence | Usage |
| --- | --- | --- |
| JOHE45057 | CACAGCTCCCAGACGCAAGC | 3' RACE of CENP-A |
| JOHE45486 | ACACTGGCGGCCGTTACTAGTGCGCTGTCCAAGCGCGCG | Clone GFP-CENP-A HDR template using NEBuilder® HiFi DNA Assembly |
| JOHE45487 | CCTTGCCCATCGCTGCCTGCCTGACTGC |  |
| JOHE45488 | GCAGGCAGCGATGGGCAAGGGCGAGGAA |  |
| JOHE45489 | GCGATGCCATCTTGTAGAGTTCATCCATGCCATGC |  |
| JOHE45490 | ACTCTACAAGATGGCATCGCCGCGTCCA |  |
| JOHE45491 | GACCATGATTACGCCAAGCTTGCGCTATACTACGCCGACCA |  |
| JOHE50156 | CTAGCACTGCTCTGATGAGTCCGTGAGGACGAAACGAGTAAGCTCGTCAGCAGTCAGGCAGGCAGCGA | sgRNA_PsCENPA_182 forward |
| JOHE50157 | AAACTCGCTGCCTGCCTGACTGCTGACGAGCTTACTCGTTTCGTCCTCACGGACTCATCAGAGCAGTG | sgRNA_PsCENPA_182 reverse |
| JOHE50062 | 5' GTTCATTTAGGGAGGTGCCACTGTA 3' | CENP-A_Junction_5' |
| JOHE50063 | 5' TCCTGCTGAAGGGGAAATAGATG 3' | CENP-A_Junction_3' |
| JOHE45358 | CGTTCACATCACCATCCAGTTCCAC | GFP_Seq_R |
| JOHE45420 | GACGGCTGCTGGGATCACGC | GFP_Seq_F |
