## Supplementary material for "Long transposon-rich centromeres in an oomycete reveal divergence of centromere features in Stramenopila-Alveolata-Rhizaria lineages": S1 Text

### S1 Text Nanopore sequencing and *de novo* assembly of the reference *P. sojae* genome

#### Nanopore sequence statistics for *P. sojae* reads

To improve the genome assembly of the *P. sojae* reference strain (P6497), the long-read Oxford Nanopore Technology (ONT) was applied. In total, four flow cells were used (one for MinION, three for GridION), which produced high quality sequence data set: Approximately 239x, 198x, and 21x coverage of the genome (95 Mb, Tyler, 2006) was obtained based on all reads, reads > 10 kb, and reads > 100 kb respectively (S1 Table). One flow cell (Flow cell 1) accompanying with MinION produced much fewer reads (S1 Table). This was probably because the flow cell used was an earlier version, and the DNA quality for the second batch (Flow cell 2-4) that was associated with GridION may be better.

#### Whole genome assembly and quality estimation

To assemble the entire genome, ONT reads ≥10 kb from all four flow cells were used for an initial assembly with SMARTdenovo (https://github.com/ruanjue/smartdenovo) (S5A Fig). The resulting contigs were corrected with two round of Racon [2] based on reads ≥10 kb derived from the ONT and the available PacBio sequencing data (SRA: SRR10759964) that were mapped to the assembly using GraphMap [3]. Afterwards, three rounds of Pilon (v1.22) [4] correction were performed using "pseudo-Illumina" read pairs generated from existing Sanger sequences [1] (ftp://ftp-private.ncbi.nlm.nih.gov/pub/TraceDB/phytophthora_sojae/) with a custom-made Perl script, which generated a read pair (read length 150 bp, insert length 500 bp) from every 35$^{th}$ base along the Sanger reads, resulting in approximately 20 million "pseudo-Illumina" read pairs with a nominal coverage of about 70 x. After getting the corrected assembly, mitochondrial sequences were removed based on the GC content and BLAST search results. The assembly was finalized as Psojae2019.1.

To further increase the contiguity of the assembly, different scaffolding programs such as npScarf [5], SSPACE [6], and LINKS [7] were implemented. All scaffolding programs greatly increased the continuity, forming assemblies having 35-49 scaffolds, with N50 3.6-5Mb. (S3 Table).

Notably, we also attempted to link the contigs using the next generation optical mapping Bionano Saphyr system, which is based on a direct label stain (DLS). In total, 30 contigs were anchored, but most could only be partially covered by the BioNano molecules (S10A Fig and S5 File), except 3 contigs (contigs 48, 63 and 50) that were fully covered (S10 E and S10F Figs). 8 contigs were suggested to join to form an assembly having 66 scaffolds (S10 B-E Figs and S5 File). In addition, one sequence contig (Contig 32) was identified as incorrectly assembled and the mis-assembly was resolved by breaking the contig at the divergent position (S10D Fig).

The low coverage obtained in Bionano mapping may in part be due to the gDNA extraction method (agarose embedding and releasing) adopted for Bionano, which was developed for mammalian systems and has not been well established for filamentous species. In addition, the DLE-1 enzyme used for labeling in the Bionano Saphyr system recommends a label density of 8 to 25 labels per 100 kb, while the density is about 9 in the *P. sojae* genome. This probably is a constraint of applying Bionano mapping to assemble “small” genomes or small contigs that do not have sufficient label density.

To evaluate the quality of the assemblies, we generated dot plot maps comparing the new genome assemblies with the existing Sanger assembly (S11 Fig). The contig-level assembly Psojae2019.1 is mostly colinear with the Sanger assembly, except two major regions which are very repetitive (5B Fig). In contrast, all *in silico* scaffolded assemblies demonstrated more conflicts when they were compared to the Sanger assembly (S6 A-C Figs). As Bionano mapping only scaffolded limited numbers of contigs, it generated an assembly showing similar collinearity as that of the P2019.1 assembly (S6D Fig). Remarkably, contigs that were joined by Bionano were also combined by the Sanger assembly.

Further examination of the scaffolded assemblies revealed that the scaffolding program SSPACE engulfed 9 (out of 13) telomeres in the assembly (S3 Table). Later centromere mapping experiments showed that npScarf linked contigs that had centromeres. These indicate that SSPACE and npScarf generated substantial errors in the scaffolded assemblies. One aspect that we observed from Bionano mapping is that several regions were duplicated in the new genome (S10 B, S10C and S10E Figs). This could be caused by higher heterozygosity of those regions, given that *P. sojae* is diploid and the genome is not 100% homozygous [8]. Taken together, despite various scaffolders stitched contigs and generated assemblies with higher contiguity, they also generated gaps or other structural errors. Without coverage support from long-reads for the joints, we have opted to retain contig-level assembly in our study.

**Metric of the Psojae2019.1 assembly**

Statistically, the resulting assembly of the nuclear genome (Psojae2019.1) has a size of 86 Mb contained in 70 gap-free contigs, with a contig N50 of 2 Mb (S5B Fig). Many contigs proved to contain long tandem tRNA repeats (Fig 3 and S6 Fig), implying that tRNA repeats are one of the main obstacles challenging the genome assembly. Interestingly, most of the tRNA repeats are homogenous. Based on searching for the motif (TTTAGGG) that was proposed for oomycete telomere repeats [9], telomeric sequences were identified at single ends of 13 contigs (S2 File). Analysis of the new assembly revealed ~31 % of repeat sequences (vs. ~27% of Sanger), and 24,415 protein-coding genes were predicted for the repeat-masked assembly (S5C Fig).
