## Supplementary material for "Long transposon-rich centromeres in an oomycete reveal divergence of centromere features in Stramenopila-Alveolata-Rhizaria lineages": S3 File

**S3 File.** **The** **5 kb consensus sequence shared by *P. sojae* centromeres, and representative full-length CoLT homologs in *P. sojae*, *B. lactucae* and *P. citricola*.**

>5kb_common_sequence_consensus

CGGATCGGAGCGTGATGGAAGGTATGGATCTTGGACACGAAGCTCGGACGAAGCGGAGGGAGGGACGTCGCAGACGGCGGGACCGGAGCGAAGGCACTTGAGGGGTAGAATGAAGATTCGAAAGTGATTGAATTGTTGGAGCTTACACCCTACTGTTATAGACACGCGACCCTGCGTCCTATCACCTAGTGCATCTAATCAACCATCACTCACCGGTGTTCATCCAATCATCCTACTCGTCTAATCATCGGTGGCGTAAACAACTTACCGTAATCACCATAATCAACGTGGGTTCGGATTGGACCGGTTCGGGTTAACCGGTTAATGGGTACAACTGGTACACAGTACCCAAGACCAGCAACCCTCAATACGCCCCCTCAATGCGAAGTCCCACGAATGCCGAGAGCCACCGTCAGATCCGTGTGGCGTGGTCCCGCAAGCGCCTTCGTCATGATGTCCGCTAGCATCACAGAAGTCTCGCAGTACTGGAGCTTGACTTCGCCGCGCTTGACTTCATCGCGGATGTGGTGGTACTTGATGTCGATGTGCTTGGCGCGACCGTGGTTCACTGGGTTCTTCGTCATCTTGATGCACGACTGGTTGTCTTCGCGGATCATGAGTTCGGGTCCAGTCTCATTAGCGGCCGCCAGAATCTCGCATAGCAGGCGATGAATCCACTTGCCTTCTTGGATAGCCAGACTGAGCGCGATGTACTCCGCTTCGCTCGTGGACAGCGACACACTTGATTGCTTCTTGCTTCCCCAGCTCACAGGCGCACTCATGAGCATGAACGTGTAGCCTGAAGTGGACTTGCGGTCGGCGAGGTCGCCGGCCCAGTCCGCGTCCGAGTAGCCACGAAAGTCGATCTTGTCGCCCGGCTTGAAGCAGATCCCATGCGTCTTGGTGCCTTGCAGGTAGCGAAAGATACGCTTGACCGCAACCCAGTGTTCCTCTTGGGGGTTCTCCATGAAGCGCGATACATAGCCAACGGCGTACGCGATGTCTGGGCGCGTGGCGGTCATCAGGTGCATCAGAGCACCAACAGCTTCGCGAAACGGAGCGTTCACCTTGGTCGCCGCGTCGCTTGGAACCAGTCGAGTGCTCATGTCCACCGGACTCACCACGGCCTTGCACTCGTCCATGGCGAAGCGCTTCAGGATGTCGTCCACGTATCGCCGCTGGCACATGGTCACGCTGCCGTCCGGGCCGTCCACCAACTCGATCCCAAGAACAAACGCGCACTTGCCGCTGTCGGTCATCTCGAAGCGCGTCTTCAGGTCGGTCTTCGTGCGTGCGATCAGCTCGGGCGAACTCCCGGTAATCAGTACGTCATCCACGTACACCAGTACAAGCACGCAGTGACCGTCCACCACCTTGATGTAGAGGCACGGATCGAAGGCCGACACTTGGAAGCCGATGGAGCACACGAACTCGTCGAAGGTCTCGTTCCACACTCGCGAAGCTTGCTTGAGACCGTAGATGGCCTTCACCAGTTCCAGGCAGTCGAAGTCACCATCCAGTTCGACGCCTTCCGGAACCACGCAGAAGACCACCTCCTTCATTATCCCGTACAGGAATGCTGTCACCACGTCGAGTTGGTCGAGGGGCCAGCCAAAATACTTAGCAATCGCGATCACCATGCGCAGCGTCACATACTTGACCACGGGCGAGAACGTCTCGGTGTAGTCAATGCCGTACTTCTGCTTGAAGCCCTTGGCCACGAGGCGCGCCTTGTACTTCTCGATCGAGCCGTCCGCTTTGCGCTTGATCTTGAACACCCACTTGGTATCGATGGCGCGTTGTCCGTTAGGCAACTTGGCAGCTCGAAACACACCACGCAGTCGCATCGACTTGAGCTCCGCGCGGATCGCTTTGCGCCAGTGCACTTGATCAGGACCGTTCACAGCCTCTTGGAACGTCGTCGGCTCCGATAGATCGGCTGCTGCTTCCACAGCGTTGGCACTGGCACGCCAAAACACTGGAGGCGTCGAGTCGTCGTCTTCATCTTGGTCCCCAGACTTTTCTTCTTCGTCGGGGTCGGCTCGGCGCGAGGAGGATACGTCCGGCGCACTTGCTTCTTCTAAGCCAGCTCGGTGGCGAACGGGGCGCGATCTTCTGGCAGTGTTGTCGTCGCCACTCGGTCGGTACTTGCGCTTCCCCGCCTGCTTGAAGCTCGTTTGGCGCGGTCCATCGTTGTCGTCCAGGTCCAGCGCGTCCAAGTCCAGCACAGCATCATCGACGTCTTCATCCGAAGTATGCGTCGAAAGTCCAAAGGTCGACTCATCGAAGTTGACATCGCGACTGATAACCACCTGCCCCGCCTCGATGTCATAAACTCGATACGCCTTGGACACTTCTTCGTAGCCGAGGAACAGCCCGGCGCGAGCCTTGGGGTCCCACTTGCGCCGTCTTTCCTTCGGCGTCAGGATGTACGTCTGGCATCCGAACACACGCATGTGCTTCACACTCGGCTTGGACTTGTAAACGATCTCGAACGGAGTCTTGTGTTCGATCTTGGGTGATGGCAGACGATTCTTCACGTAGATGGCTGTCATGGCCGCCTCAGCCCAGAAGCTCTTGTCCAGCTTGGCGTGATGCAGCATGCTGCGGCCAATCGTGACGATGGTCCGGATGGCGCGTTCTGCCGTGCCGTTGGTCTGGTGCGCGTACGGAACCGTGGTTTGCTGCTCGATGCCTTCGACTTCGTAGAAGGCCTTGAGCGAGTTGGTCGCGAACTCACGGGCTCCGTCGTGCCGCACAAACTTGACCTTCTTGCCGAACTGCGTCTGCACCTTCGTGACGTACATCTTGATGCAGTCCTCGCTCTCCGACTTGGCACGCAGGCAGAACCCCTTCATGCATCCGCTGGCTTCGTCCACGATCAGCAGCAGATACTTGCTGCCACCCAGTGACTCTTCTTCCATGGGGCCGCACACGTCGAAGTGGAGCAGCTCGAACGTGTCGTAGTGACGCTTGTCACGGTTGCTTGGGAACGGCTTCTGGACCATTTTCCCTTGCTGGCAGCCGCGACACACGCTTGGTCCACTCGACTTGGAAGGTGCCTTCGCGTCCTTGATCATGTTGTTGTCGACCATCTTGCGAAGCACCTCCACCGGAGCGTGACCCATACGCGCGTGTAAGTCGACCGTCTTCCCACTGGTCGCCGCATTCGCCGATCGCTGAGGCGTCTGGAGCCAGTAGAGTCCATCCACGAGGTTTGCCGTCGCCACCACTTGCTTGGAGCCCTTGTGCGCAATCTGCATCTTGGTGCTGTCGAACACGACTTGGAACAGACCACTCTTGTTGATCTGTGGCACCGAAAGCAGATTCTTGCTCATGCTTGGGACGTACAGCGCGTTCTTGATCTCGATGTCGCGCTCGTCGCCGTTGGGCAGAACCACTCGTTCCACGATGGTCCCCACACCCATGATGGCCGCCTTGTTGCCGTCGGCCACCGAAAGTTCGCCTTCATTCCGCTCAATCAAATGAGCGAACTTGGCCTTGTCGTTGCAGATGTGGTGCGTTGCGCCGCTGTCGACCGCCCACAACCCGGACACATTCTTGCCCGTCGAAATGCCACACTCCAGTGAGACCGCGAACGCCACTCGATCATAGCCGTAGTCGTAGTTGTAGCTGGCGTCGTCGTTCCTCCACTGGGCATTGTTGGCCCCGCGTCCACGGCCATTGCCATTGCCGCCGCGTCTTGGGCCCCGATTCTCGTCCTTCTGCTTGGTCCAGCACTTGTCAATCGTGTGCCCCAATTTCCCGCAGTACGTACACTGGCGAGACTCTCGGTCGGTGTTGAACGCCTTGGCCGCATTCTCCGTCTTCACCGTCGCCGTCTTCTCGCCTTGTCTCTTGATGTGCTCATTCGTCAGCACCTTCACGACGTCTTGCGATCGTAGTTCAGCGCTGCTCATCTCCAAGTTCAGCACGACGTTCTCGTAGCTCTTGGGGAGGCTGCGCAACAGGCAGATCGCCACGTCCTCATCCTCCATCTTGGCGCCGATGCTGCTGAGCTTGGCGCTGATGTTGAGGACCTCGTTGCAATGATGCAAGACGTTGCCGCCTTCCGACATCTCCATGCTGAAGAGTTGGCGCTTCAGGTAGATGCGTCCAGCCTTCTGCGAACCACCGTACAGCGTCTTCAGGCGCGTCCACGCAACCCACGCTTCCTCGCAGTCGTCGACCACATGGTGGTAGTCCGCGTCCATGTGGAGCAACATCAGTCCGAACGCGATGTTGCTCGACTTGACGTAGTCGTCCTTGGCCCGCGAGTCCGTGAAGCTCGGCGTCGTCTCGCGGTTCACGACATGCCACGTCGACTTGGTCAGGAACACGCCACGCATGTAGCGGTTCCACGTCGCGTAGTTGTCCCCGTTGAACTTGTCAATGGAGACCTTGGTCGAGGCGTCTTGGGCAGAGGACATGGTGCTTGGGCTCATAACCTGATGGAAGGTATGGATCTTGGACACGAAGCTCGGACGAAGCGGAGGGAGGGACGTCGCAGACGGCGGGACCGGAGCGAAGGCACTTGAGGGGTAGAATGAAGATTCGAAAGTGATTGAATTGTTGGAGCTTACACCCTACTGTTATAGACACGCGACCCTGCGTCCTATCACCTAGTGCATCTAATCAACCATCACTCACCGGTGTTCATCCAATCATCCTACTCGTCTAATCATCGGTGGCGTAAACAACTTACCGTAATCACCATAATCAACGTGGGTTCGGATTGGACCGGTTCGGGTTAACCGGTTAATGGGTACAACTGGTACACAGTACCCAAGACCAGCAACCCTCAATAGNNNGCCNTCACTNNNCTNCNNNTTACTMCATCTCAGAATGCACT

**>P6497_smrtdenovo_ONT10kb_corrected_contig_38_322956-327751(+)**

GGGTGCCTATTCCGACTGAAGAACAAGTATTGAGGGTTGCTGGTCTTGGGTACTGTGTACCAGTTGTACCCATTAACCGGTTAACCCGAACCGGTCCAATCCGAACCCACGTTGATTATGGTGATTACGGTAAGTTGTTTACGCCACCGATGATTAGACGAGTAGGATGATTGGATGAACACCGGTGAGTGATGGTTGATTAGATGCACTAGGTGATAGGACGCAGGGTCGCGTGTCTATAACAGTAGGGTGTAAGCTCCAACAATTCAATCACTTTCGAATCTTCATTCTACCCCTCAAGTGCCTTCGCTCCGGTCCCGCCGTCTGCGACGTCCCTCCCTCCGCTTCGTCCGAGCTTCGTGTCCAAGATCCATACCTTCCATCAGGTTATGAGCCCAAGCACCATGTCCTCTGCCCAAGACGCCTCGACCAAGGTCTCCATTGACAAGTTCAACGGGGACAACTACGCGACGTGGAACCGCTACATGCGTGGCGTGTTCCTGACCAAGTCGACGTGGCATGTCGTGAACCACGAGACGACGCCGAGCTTCACGGACTCGCGGGCCAAGGACGACTACGTCAAGTCGAGCAACATCGCGTTCGGACTGATGTTGCTCCACATGGACGCGGACTACCACCATGTGGTCGACGACTGCGAGGAAGCGTGGGTTGCGTGGACGCGCCTGAAGACGCTGTACGGTGGTTCGCAGAAGGCTGGACGCATCTACCTGAAGCGCCAACTCTTCAGCATGGAGATGTCGGAAGGCGGCAACGTCTTGCATCATTGCAACGAGGTCCTCAACATCAGCGCCAAGCTCAGCAGCATCGGCGCCAAGATGGAGGATGAGGACGTGGCGATCTGCCTGTTGCGCAGCCTCCCCAAGAGCTACGAGAACGTCGTGCTGAACTTGGAGATGAGCAGCGCTGAACTACGATCGCAAGACGTCGTGAAGGTGCTGACGAATGAGCACATCAAGAGACAAGGCGAGAAGACGGCGACGGTGAAGACGGAGAATGCGGCCAAGGCGTTCAACACCGACCGAGAGTCTCGCCAGTGTACGAACTGCGGGAAATTGGGGCACACGATTGACAAGTGCTGGACCAAGCAGAAGGACGAGAATCGGGGCCCAAGACGCGGCGGCAATGGCAATGGCCGTGGACGCGGGGCCAACAATGCCCAGTGGAGGAACGACGACGCCAGCTACAACTACGACTACGGCTATGATCGAGTGGCGTTCGCGGTCTCACTGGAGTGTGGCATTTCGACGGGCAAGAATGTGTCCGGGTTGTGGGCGGTCGACAGCGGCGCAACGCACCACATCTGCAACGACAAGGCCAAGTTCGCTCATTTGATTGAGCGGAATGAAGGCGAACTTTCGGTGGCCGACGGCAACAAGGCGGCCATCATGGGTGTGGGGACCATCGTGGAACGAGTGGTTCTGCCCAACGGCGACGAGCGCGACATCGAGATCAAGAACGCGCTGTACGTCCCAAGCATGAGCAAGAATCTGCTTTCGGTGCCACAGATCAACAAGAGTGGTCTGTTCCAAGTCGTGTTCGACAGCACCAAGATGCAGATTGCGCACAAGGGCTCCAAGCAAGTGGTGGCGACGGCAAACCTCGTGGATGGACTCTACTGGCTCCAGACGCCTCAGCGATCGGCGAATGCGGCGACCAGTGGGAAGACGGTCGACTTACACGCGCGTATGGGTCACGCTCCGGTGGAGGTGCTTCGCAAGATGGTCGACAACAACATGATCAAGGACGCGAAGGCACCTTCCAAGTCGAGTGGACCAAGCGTGTGTCGCGGCTGCCAGCAAGGGAAAATGGTCCAGAAGCCGTTCCCAAGCAACCGTGACAAGCGTCACTACGACACGTTCGAGCTGCTCCACTTCGACGTGTGCGTCCCCATGGAAGAAGAGTCACTGGGTGGCAGCAAGTATCTGCTGCTGATCATGGACGAAGCCAGCGGATGCATGAAGGGATTCTGCCTGCGTGCCAAGTCGGAGAGCGAGGACTGCATCAAGATGTACGTCACGAAGGTGCAGACGCAGTTCGGCAAGAAGGTCAAGTTTGTGCGGCACGACGGAGCCCGTGAGTTCGCGACCAACTCGCTCAAGGCCTTCTACGAAGTCG**AAGGCATCGAGCAGCAAACCACGGTTCCGTACGCGCACCAGACCAACGGCACGGCAGAACGCGCCATCCGGACCATCGTCACGATTGGCCGCAGCATGCTGCATCACGCCAAGCTGGACAAGAGCTTCTGGGCTGAGGCGGCCATGACAGCCATCTACGTGAAGAATCGTCTGCCATCACCCAAGATCGAACACAAGACTCCGTTCGAGATCGTTTACAAGTCCAAGCCGAGTGTGAAGCACATGCGTGTGTTCGGATGCCAGACGTACATCCTGACGCCGAAGGAAAGACGGCGCAAGTGGGACCCCAAGGCTCGCGCCGGGCTGTTCCTCGGCTACGAAGAAGTGTCCAAGGCGTATCGAGTTTATGACATCGAGGCGGGGCAGGTGGTTATCAGTCGCGATGTCAACTTCGATGAGTCGACCTTTGGACTTTCGACGCATACTTCGGATGAAGACGTCGATGATGCTGTGCTGGACTTGGACGCGCTGGACCTGGACGACAACGATGGACCGCGCCAAACGAGCTTCAAGCAGGCGGGGAAGCGCAAGTACCGACCGAGTGGCGACGACAACACTGCCAGAAGATCGCGCCCCGTTCGCCACCGAGCTGGCTTAGAAGAAGCAAGTGCGCCGGACGTATCCTCCTCGCGCCGAGCCGACCCCGACGAAGAAGAAAAGTCTGGGGACCAAGATGAAGACGACGACTCGACGCCTCCAGTGTTTTGGCGTGCCAGTGCCAACGCTGTGGAAGCAGCAGCCGATCTATCGGAGCCGACGACGTTCCAAGAGGCTGTGAACGGTCCTGATCAAGTGCACTGGCGCAAAGCGATCCGCGCGGAGCTCAAGTCGATGCGACTGCGTGGTGTGTTTCGAGCTGCCAAGTTGCCTAACGGACAACGCGCCATCGATACCAAGTGGGTGTTCAAGATCAAGCGCAAAGCGGACGGCTCGATCGAGAAGTACAAGGCGCCCCTCGTGGCCAAGGGCTTCAAGCAGAAGTACGGCATTGACTACACCGAGACGTTCTCGCCCGTGGTCAAGT**ATGTGACGCTGCGCATGGTGATCGCGATTGCTAAGTATTTTGGCTGGCCCCTCGACCAACTCGACGTGGTGACAGCATTCCTGTACGGGATAATGAAGGAGGTGGTCTTCTGCGTGGTTCCGGAAGGCGTCGAACTGGATGGTGACTTCGACTGCCTGGAACTGGTGAAGGCCATCTACGGTCTCAAGCAAGCTTCGCGAGTGTGGAACGAGACCTTCGACGAGTTCGTGTGCTCCATCGGCTTCCAAGTGTCGGCCTTCGATCCGTGCCTCTACATTAAGGTGGTGGACGGTCACTGCGTGCTTGTACTGGTGTACGTGGATGACGTACTGATTACCAGGAGTTCGCCCGAGCTGATCGCACGCACGAAGACCGACCTGAAGACGCGCTTCGAGATGACCGACAGCGGCAAGTGCGCGTTTGTTCTTGGGATCGAGTTGGTGGACGGCCCGGACGGCAGCGTGACCATGTGCCAGCGGCGATACGTGGACGACATCCTGAAGCGCTTCGCCATGGACGAGTGCAAGGCCGTGGTGAGTCCGGTGGACATGAGCACTCGACTGGTTCCAAGCGACGCGGCGACCAAGGTGAACGCTCCGTTTCGCGAAGCTGTTGGTGCTCTGATGCACCTGATGACCGCCACGCGCCCAGACATCGCGTACGCCGTTGGCTATGTATCGCGCTTCATGGAGAACCCCCAAGAGGAACACTGGGTTGCGGTCAAGCGTATCTTTCGCTACCTGCAAGGCACCAAGACGCATGGGATCTGCTTCAAGCCGGGCGACAAGATCGACTTTCGTGGCTACTCGGACGCGGACTGGGCCGGCGACCTCGCCGACCGCAAGTCCACTTCAGGCTACACGTTCATGCTCATGAGTACGCCTGTGAGCTGGGGAAGCAAGAAGCAATCAAGTGTGTCGCTGTCCACGAGCGAAGCGGAGTACATCACGCTCAGTCTGGCTATCCAAGAAGGCAAGTGGATTCATCGCCTGCCATGCGAGATTCTGGCGGCCGCTAATGAGACTGGACCCGAACTCATGATCCGCGAAGACAACCAGTCGTGCATCAAGATGACGAAGAACCCAGTGAGCCACGGTCGCGCCAAGCACATCGACATCAAGTACCACCACATCCGCGATGAAGTCAAGCGCGGCGAAGTCAAGCTCCAGTACTGCGAGACTTCTGTGATGCTAGCGGACATCATGACGAAGGCGCTTGCGGGACCACGCCACACGGATCTGACGGTGGCTCTCGGCATTCGTGGGACTTCGCATTGAGGGGGCGTATTGAGGGTTGCTGGTCTTGGGTACTGTGTACCAGTTGTACCCATTAACCGGTTAACCCGAACCGGTCCAATCCGAACCCACGTTGATTATGGTGATTACGGTAAGTTGTTTACGCCACCGATGATTAGACGAGTAGGATGATTGGATGAACACCGGTGAGTGATGGTTGATTAGATGCACTAGGTGATAGGACGCAGGGTCGCGTGTCTATAACAGTAGGGTGTAAGCTCCAACAATTCAATCACTTTCGAATCTTCATTCTACCCCTCAAGTGCCTTCGCTCTGGTCCCGCCGTCTGCGACGTCCCTCCCTCCGCTTCGTCAGAGCTTCGTGTCCAAGATCCATACCTTCCATCAGCGACAATAGATTCGCAAGCTGATCGC

ATG – predicted Start codon

TAG – predicted stop codon

The LTR marks (5’-TG..CA-3’) are underlined within the LTR regions highlighted in cyan.

**Nucleotides in bold**: The probe (GenBank: JF519821.1) used for fingerprinting of *P. sojae*

Nucleotides underlined: The probe (GU244585.1) used for fingerprinting of *P. sojae*

Alignment of the Left and Right LTRs

```
Left       TGAGGGTTGCTGGTCTTGGGTACTGTGTACCAGTTGTACCCATTAACCGGTTAACCCGAA	60
Right      TGAGGGTTGCTGGTCTTGGGTACTGTGTACCAGTTGTACCCATTAACCGGTTAACCCGAA	60
           ************************************************************

Left       CCGGTCCAATCCGAACCCACGTTGATTATGGTGATTACGGTAAGTTGTTTACGCCACCGA	120
Right      CCGGTCCAATCCGAACCCACGTTGATTATGGTGATTACGGTAAGTTGTTTACGCCACCGA	120
           ************************************************************

Left       TGATTAGACGAGTAGGATGATTGGATGAACACCGGTGAGTGATGGTTGATTAGATGCACT	180
Right      TGATTAGACGAGTAGGATGATTGGATGAACACCGGTGAGTGATGGTTGATTAGATGCACT	180
           ************************************************************

Left       AGGTGATAGGACGCAGGGTCGCGTGTCTATAACAGTAGGGTGTAAGCTCCAACAATTCAA	240
Right      AGGTGATAGGACGCAGGGTCGCGTGTCTATAACAGTAGGGTGTAAGCTCCAACAATTCAA	240
           ************************************************************

Left       TCACTTTCGAATCTTCATTCTACCCCTCAAGTGCCTTCGCTCCGGTCCCGCCGTCTGCGA	300
Right      TCACTTTCGAATCTTCATTCTACCCCTCAAGTGCCTTCGCTCTGGTCCCGCCGTCTGCGA	300
           ****************************************** *****************

Left       CGTCCCTCCCTCCGCTTCGTCCGAGCTTCGTGTCCAAGATCCATACCTTCCATCA	355
Right      CGTCCCTCCCTCCGCTTCGTCAGAGCTTCGTGTCCAAGATCCATACCTTCCATCA	355
           ********************* *********************************
```

**>P6497_smrtdenovo_ONT10kb_corrected_contig_38_322956-327751(+)**

MSPSTMSSAQDASTKVSIDKFNGDNYATWNRYMRGVFLTKSTWHVVNHETTPSFTDSRAKDDYVKSSNIAFGLMLLHMDADYHHVVDDCEEAWVAWTRLKTLYGGSQKAGRIYLKRQLFSMEMSEGGNVLHHCNEVLNISAKLSSIGAKMEDEDVAICLLRSLPKSYENVVLNLEMSSAELRSQDVVKVLTNEHIKRQGEKTATVKTENAAKAFNTDRESRQCTNCGKLGHTIDKCWTKQKDENRGPRRGGNGNGRGRGANNAQWRNDDASYNYDYGYDRVAFAVSLECGISTGKNVSGLWAVDSGATHHICNDKAKFAHLIERNEGELSVADGNKAAIMGVGTIVERVVLPNGDERDIEIKNALYVPSMSKNLLSVPQINKSGLFQVVFDSTKMQIAHKGSKQVVATANLVDGLYWLQTPQRSANAATSGKTVDLHARMGHAPVEVLRKMVDNNMIKDAKAPSKSSGPSVCRGCQQGKMVQKPFPSNRDKRHYDTFELLHFDVCVPMEEESLGGSKYLLLIMDEASGCMKGFCLRAKSESEDCIKMYVTKVQTQFGKKVKFVRHDGAREFATNSLKAFYEVEGIEQQTTVPYAHQTNGTAERAIRTIVTIGRSMLHHAKLDKSFWAEAAMTAIYVKNRLPSPKIEHKTPFEIVYKSKPSVKHMRVFGCQTYILTPKERRRKWDPKARAGLFLGYEEVSKAYRVYDIEAGQVVISRDVNFDESTFGLSTHTSDEDVDDAVLDLDALDLDDNDGPRQTSFKQAGKRKYRPSGDDNTARRSRPVRHRAGLEEASAPDVSSSRRADPDEEEKSGDQDEDDDSTPPVFWRASANAVEAAADLSEPTTFQEAVNGPDQVHWRKAIRAELKSMRLRGVFRAAKLPNGQRAIDTKWVFKIKRKADGSIEKYKAPLVAKGFKQKYGIDYTETFSPVVKYVTLRMVIAIAKYFGWPLDQLDVVTAFLYGIMKEVVFCVVPEGVELDGDFDCLELVKAIYGLKQASRVWNETFDEFVCSIGFQVSAFDPCLYIKVVDGHCVLVLVYVDDVLITRSSPELIARTKTDLKTRFEMTDSGKCAFVLGIELVDGPDGSVTMCQRRYVDDILKRFAMDECKAVVSPVDMSTRLVPSDAATKVNAPFREAVGALMHLMTATRPDIAYAVGYVSRFMENPQEEHWVAVKRIFRYLQGTKTHGICFKPGDKIDFRGYSDADWAGDLADRKSTSGYTFMLMSTPVSWGSKKQSSVSLSTSEAEYITLSLAIQEGKWIHRLPCEILAAANETGPELMIREDNQSCIKMTKNPVSHGRAKHIDIKYHHIRDEVKRGEVKLQYCETSVMLADIMTKALAGPRHTDLTVALGIRGTSH*

GAG, Capsid protein (pfam14223)

PR, GAG-pre-integrase (pfam13976)

INT, Integrase (pfam00665)

RT, Reverse transcriptase (pfam07727)

RH, RNase H (cd09272)

Domains were annotated based on CD-search, hereafter.

**>Bremia_lactucae_Sca7_2388-7076(+)**

ATGTCTTGCGTCGAAGAGTATTGAGGGTTATGGGTTTCAGGTAATGTACACAACCCAAACCGGTCCAATCCGAACCGCGGTTAAACATAGCATCTACTTATATTATAGTATAGTTAGATGCAGTATGTAGAAATGAGTGGTTTCGAGAAGTAAAAGTGTAAGCCCCAATAAATCAATCACTTCCAAATATTCCTTCTCCTTCACAAGTCGTCTTCACCTTTCAAGTGCAGACTTCTCTCGTGTTCTTCAAGCCTCCTTCTCTCTTCCAGTGCTCGGTACCCTTTAACAGGTTATGAGCCCAAGCACGATGTCCTCTGCTCAAGATTCCGCCACCAAGATCTCCATCGACAAATTCGATGGCGACAACTATGCGACGTGGAGCCGCTACATGCGTGGCGTGTTCCTGACCAAGTCGACTTGGCATGTGGTCAACCGGGAGACTACTCCAACCTTCGCCGATCCTCGCGCCATGGACGAATATGTCAAGGCGAGCAATATCGCCTTCGGCTTGATGTTGCTGTACATAAGCGCGGATTACCATCACGTGGTTGATGACTGCGAAGAAGCATGGGTTGCGTGGGCAAAGCTGAAGACCCTCTACGGCGGATCGCAGAAGGCAGGACGGATCTTCTTGAAGCGACAGCTCTTCAGCATGGAGATGGCGGAAGGCGGTAATGTACTTCATCATTGCAATGAAGTCCTCAACATCAGCGCAAAGCTCAAGAGCATCGGCGCCAAGATGGAGGACGAGGACGTGGCGATTTGCTTGTTGCGCAGCCTCCCCAAGAGTTACGAGAACGTCGTTCTCAACTTGGAGATGAGCAGCGCGGAATTGCGATCGCGGGACGTCGTGAAAGTGCTGACAAATGAGCACATCAAGAGACAAGGAAACAAGACGACATCGGTGAAGACTGAAGAAACAACCAAGGCTTTCAGTACCGAACGCGAGTTTCGTCAGTGTACATTCTGCGGAAAATTGGGGCACACTGTGGAGAAGTGCTGGACCAAGCAGAAGGAAGACAATCGGGGAGCTCGACGCGGCGGCAATTATCGTGGACGCGGAGCGAATCAAGTTCAGTGGCAAGGCTACAGTGGCAACAGCAACGGTAACTACGATCGTGTGGCGTTTGCTGTGTCAATGAAATGCGGACTTTCAACGACCAAGAACATGTCCGGGATGTGGGCGATTGATAGCGGTGCTACGCATCACATCTGCTACGAAAAATCCAAGTTCGAAGTTCTGGACGAACACAATGAAGGTGAAGTTCTGGTCGCAGATGGAAATAAAGCCGCTATCAAAGGCATCGGGACTATCATCGAAAAGGTACTACTGCCTAACGGTGAAGAACGTGAAGTCGAGATCAAAAACGCGCTTTTCGTCCCAGATATGAACAAGAACCTACTATCAATTCCGCAAATCAACAAGAGCGGCAAGTTTCAAGTGGTGTTTGATGGTACCAAAATGGAGATCTCGCGTAAAGACCCTGAGCAAGTGGTGGCGATGGCAGATCTTGCGGATGGACTCTATTGGCTTCGCACATCTCACCGATCGATAAATACTACATCAAGGTCAAGAGTCGAAGATCTTCATGCACGCATGGGCCACGCACCGTTTGATGTCTTGTGCAAAATGGTTTCCAAAGGGATGATCAAGGATGCTAAAATGCCAGAAAAGGCAAGCAGATCTAGCATATGCCACGGGTGTCAAGAAGGAAAAATGGTACAGCGACCGTTTCCAAGTAACCCGAACAAGCAGCACTACGACCCGTTCGAGCTTCTACACATCGACACTTGCGGACCCATGGAAGTTGAATCGCTGGGTGGAAGCAAGTACTTGCTACTAATCGTCGATGAAGGCAGCGGATGCATGAAGGGCTTCGGTCTGCGCGCCAAGTCTGATAGTGAAGAGTGTATCAAGAGATACATCAATGCGGTGCAGACGCAGTTTGACTTCAAGGTCAAGTTCGTTCGACATGATGGTGCACGAGAATTCGCGACAACTTCGATCAAGGCATTTTACGATGATCAAGGAATCGAGCAGCAAGTCACTGTTCCGTATGCGCACCAGACGAATGGCACGGCAGAACGGGCAATCCGGACTATCGTAACGATTGGCCGCAGCATGCTCCACTACGCAAAATTAGACAAGTGTTTCTGGGCTGAGGCAGCAATGACAGCAATATACATCAAGAATCGACTGCCATCGCCCAAGTACCAAGATCAGACTCCGTTCGAGATCATCAACGGATCTCGACCAAGTGTCAAACACATGCGAGTTTTTGGCTGCCGTGCGTTCGTGCTGACTCCAAAAGAAAAGAGATCGAAGTGGGATCCGAAGTCCCGTGAAGGACTGTTTATGGGGTACGAAGAAGCATCAAAGGCATACCGCATTTACGACATTGAAACGGATCAGGTGGTGATATCTCGCGATGTCACGTTCGACGAATCGACGTTTGGGTTCGCACAGACGCCTCTACAAGATGTTGTTGATGATACCTTATTGGATTTTGACACGATGAGCATCAGTAGTGAACCTGCCACCACGGAGTTCAAGCAAACAGGCAAGCGCAAGAACCGTTCAAACAGCCAAGAACAAGTGTTTAAACGACCAGCTTGCCGCGGAGCCGGGTTAGAAGAAGCAAGTGCGCCGGATGACTTCGAGTTGCGCCACCAAAAACGACGTTCGAATGTTCGAGCTAACCGGGACGAAGAGCAAAAGGACAATGATATGGATGATGATGACGACACCCACCACGGTTTTTGGCAAGCAAGCGTGAATGCAGTTGAGGGCGCTGATTTATCCGAACCAACAACGTTTGAAGACGCGATTAATGGGCCAGATCGAGTACACTGGCGTAAGGCGATCTGTGCGGAGCTAGACTCAATGAAACTCCGTAGCGTATTTCGAGCAACGAAGCTGCCAGCTGGGCAGCACACCATTGGGACCAAGTGGGTATTCAAGATCAAGCGCAAGGCTGACGGAAACATCGATAAATACAAGGCACGTCTCGTTGCCAAGGGTTTCAAGCAAAAGTATGGCATCGACTACACGGAGACGTTTTCACCAGTAGTCAAGTGCGTGACGCTGCGCATGGTGATTGCACTCACAAAGTACTTCGACTGGACACTGGACCAACTGGACGTGGTGACAGCTTTTCTGTATGGAGAAATGAAGGAAAGAGTATTCTGTGCGGTCCCAGAAGGAGTCGACGTCAATAAAGATTTTGATTGCTTTGAACTGGTTAAAGCAATCTACGGACTAAAACAAGCTTCACGGGTATGGAATGACACTTTTCATGAGTTTGTTTGCTCCATCGGATTTCAAGTCTCCGATTTTGATCCGTGCCTTTATCTCAAGATTGCAAACGGAGAGTGCGTGCTATTGTTGGTCTATGTTGATGATGTACTGGTGACTGGGAGTTCAATTGAAATGATTGCGCACACCAAGAAAGACTTAAAGTCACGTTTTGAGATGACTGATAGTGGCAAGTGCGCTTTTGTCCTCGGCATCGAGCTGGTAGACAACAATGACGGAAGCGTGACGATGTGCCAGCGACGCTACGTGGACGATGTTCTCAAGCGTTTCGGCATGAGTGACTGCAAAGCTGTCATCAGCCCAATGGACATCAGCTCTCGACTAACGTCAAGCAACGCAGCAACTAAAATCAACGTTCCGTTTCGTGAAGCAGTGGGTGCTTTAATGCACTTGATGACCGCGACACGACCAGACATTGCCTTCGCCGTGGGTTACGTGTCGCGTTTTATGGAAAACCCACAAGTCGAGCATTGGATGGCGGTGAAGCGCATCCTTCGATACTTGCAAGGAACCAAGTCGCATGGAATCTGCTTTAAGCCTGGCAACAATGTGGATTTTTGCGGCTACTCGGATGCAGACTGGGCCGGTGATCATGCAGACCGCAAGTCGACTTCAGGATACGCGTTTATTCTCATGGGCGCTCCTGTAAGCTGGGGGAGCAAGAAACAATCAAGTGTGTCGCTGTCAACAAGTGAAGCCGAATATATCGCGCTAAGTTTAGCGATTCAAGAAGGCAAGTGGGTGCATCGACTACTGCGCGAAATCCTAGACGCGGCAGCAAACAAGTCTGAACCGGTGCTTAAGATCATGGAAGACAACCAGTCGTGTATCAAGATGACGAAGAATCCGGTAAACCATGGTCGTGCGAAACATATCGACATCAAGTATCACCACATTCGTGATGAAGTAAAGCGTGGTGATGTCAAGTTGGAATACTGCGAGACTTCGAAGATGTTGGCGGATATCATGACGAAAGGACTGCCAGGACCGCGTCACAAGGAATTAACGACAGCGCTTGGCATACACGCGTGTTAGCATTGAGGGGGCGTATTGAGGGTTATGGGTTTCAGGTAATGTACACAACCCAAACCGGTCCAATCCGAACCGCGGTTAAACATAGCATCTACTTATATTATAGTATAGTTAGATGCAGTATGTAGAAATGAGTGGTTTCGAGAAGTAAAAGTGTAAGCCCCAATAAATCAATCACTTCCAAATATTCCTTCTCCTTCACAAGTCGTCTTCACCTTTCAAGTGCAGACTTCTCTCGTGTTCTTCAAGCCTCCTTCTCTCTTCCAGTGCTCGGTACCCTTTAACAAAGCCCCGTCATGCCTCACAAACTTGACTTTTTCCGAACTGCGTTAGGACCTTCA

ATG – predicted Start codon

TAG – predicted stop codon

TGA – The corresponding predicted stop codon in *P. sojae*

The LTR marks (5’-TG..CA-3’) are underlined within the LTR regions highlighted in cyan.

**>P_citricola_Sca60_26953-31785(-) (predicted coding region is truncated)**

GGGTGAACTTCCGGTAATCAGTATTGAGGGTTGCTGGTCTTGGGTACTGTGTACCAGTTGTACCCATTAACCGGTCAACCCGAACCGGTCCAATACGAACCCACGTTGATTATGGTGATTACGGTAAGTTGTTTACGCCACCGATGATTAGACGAGTAGGATGATTGGACGAACACCGATGAGTGATGGTTGATTAGATGCACTAGGTGATAGGACGCAGGGTCGCGTGTCTATAACAGTAGGGTGTAAGCTCCAACAATTCAATCACTTTCGAATCTTCATTCTACCCCTCAAGTGTCTTCGCTCCGGTCCCGCCGTCTGCGACGTCCCTCCCTCCGCTTCGTCCAAGCTTCGTGTCCAAGATCTATACCTTCCATCAGGTTATGAGCCCAAGCACCATGTCCTCTGCGCAAGACGCCTCGACCAAGGTCTCCATTGACAAGTTCAACGGGGACAACTACGCGACGTGGAACCGCTACATGCGTGGCGTGTTCCTGACCAAGTCGACGTGGCATGTGGTGAACCGCGAGACGACGCTGAGCTTCACGGACTCGCGGGCCAAGGACGAGTACATCAAGTCGAGCAACATCGCGTTCGGACTGATGTTGCTCCACATGGACGCGGACTACCACCACGTGGTCGACGACTGCGAGGAAGCGTGGGTTGCGTGGACGCGCCTGAAGACGCTGTACGGTGGTTCGCAGAAGGCTGGTCGCATCTACCTGAAGCGCCAACTCTTCAGCATGGAGATGTCGGAAGGCGGCAATGTGTTGCATCATTGCAATGAGGTCTCAACATCAGCGCCAAGCTCAGCAGCATCGGCGCCAAGATGGAGGACGAGGACGTGGCGATCTGCCTGTTGCGCAGTCTCCCCAAGAGCTACGAGAACGTCGTGCTGAACTTGGAGATGAGCAGCGCTGAACTGCGGTCGCAAGACGTCGTGAAGATGCTCACGAATGAGCACATCAAGCGGCAAGGCGAGAAGACGGCGACGGTGAAGACGGAGAACGCGGCCAAGGCGTTCAACACCGACCGAGAGTCTCGCCAGTGCACGTACTGCGGTAAATTAGGGCAAACGGTGGACAAGTGCTGGACCAAGCAGAAGGACGAGAATCGGGGCCCAAGACGCGGCGGCAATGGTAATGGCCGTGGACGCGGGGCCAACAACATCCAGTGGAGAAACGACAACGACGGCTACGACTACGACTACGACTACGATCGAGTGGCGTTCGCGGTTTCCATGGAATGTGGTATTCCGACGGGCAAGAATGTGTCCGGGATGTGGGCGGTCGACAGCGGCGCAACGCACCACATCTGCAACGACAAGGCCAAGTTCGCCTATTTGATCGAGCGAAACGAGGGCGAACTTTCGGTGGCCGACGGCAACAAGGCGGCCATCATGGGTGTGGGGACCATCGTGGAACGAGTGGTTCTGCCCAACGGCGACGAGCGCGAGATCGAGATCAAGAACGCGCTGTACGTGCCAAGCATGAGCAAGAATCTGCTTTCGGTACCGCAGATCAACAAAAGTGGCAAGTTCCAAGTGGTGTTTGACGGCGCCAAGATGCAGATTGCGCGCAAGGACTCCAAGCAAGTAGTGGCGACGGCAAACCTCGTGGATGGACTCTACTGGCTGATGACGCCTCAGCGATCGGCGAATGCGGCGACGAGTGGCAAGATTGTCGACCTGCACGCGCGCATGGGTCACGCTCCGGTTGAGGTGCTTCGCAAGATGGTCGACAACAACATGATCAAGACGCGAAGGCACCTTCCAAGTCGAGCGGACCAAGTCTGTGTCGTGGTTGCCAGCAAGGGAAGATGGTCCAAAACCGTTCCCAAGCAACCGTGACAAGCGCCACTACGACACCTTCGAGTTGCTCCACTTCGACATCTGCGGACCCATGGAGAAAAACTCGCTTGGCGGCAGCAAGTACCTGCTGCTGATCGTGGACGAAGCCAGCGGATGCATGAAGGGTTTCTGCCTGCGTGCCAAGTCGGAGAGCGAAGACTGCATCAAGACGTACATCATGAAGGTGCAGAAGCAGTTCGGCAAGAAGGTCAAGTTTGTGCGGCACGACGGAGCTCGTGAGTTTGCGACTAACTCGCTCAAGGATTTCTACGAAGACGAAGGCATCGAGCAGCAAACCACGGTCCCGTACGCGCACCAGACCAACGGCACGGCAGAGCGAGCTATCCGGACTATCGTCACGATCGGCCGCAGCATGCTGCATCACGCTAAGCTGGACAAGTGTTTCTGGGCTGAGGCGGCAATGACGGCCATCTACGTGAAGAACCGTCTGCCGTCACCCAAGATCGAGCACAAGACTCCGTTCGAGATCGTGTACAAGTCCAAGCCAAGTGTGAAGCACATGCGTGTGTTCGGCTGTCGGACGTACATTCTGACTCCGAAGGAGAAGCGGCTCAAGTGGGATCCCAAGGCTCGCGCCGGATTGTTTCTCGGCTACGAAGAAGTGTCCAAGGCGTATCGAGTTTATGACATCGAGGCGGGGCAGGTGGTCATCAGTCGCGATGTCAACTTCGATGAGTCGGCCTTTGGACTTTCGCCGCAGATTTCCGATGAAGATGTCGATGACCTGGACTTCGACTCGCTCGACATCGACGATGACGGTCCTCGCCAGACGGAATACAAGCAGGCGGGGAAACGCAAGAGTCGCCCAAGTGACGAAGATGAAGGTACCAGAAGACCGCGCACTGTGCGTCATCGGCCTGGATTGGAAGAAGCAAGTGCGCCGGACTCCTCTTCGCACCGAGTGGACACCAACGAAGAAGAAAAGTCTGGTGACCAAGATGAAGAATCGACTTCTCCTGTGTTTTGGCGTGCCAGTGCCAACGCTGTCGAAGCAGCAGCCGATCTATCGGACCCGTCGACATTCCAAGAAGCGGTGAACGGTCTGATCAAGTGCACTGGCGCAAGGCTATTCGTGCGGAGCTCAAGTCGATGCGACTTCGTGGTGTGTTTCGTGCTGCCAAGTTGCCTAACGGACAACGCGCCATCGGCACCAAGTGGGTGTTCAAGATCAAGCGCAAAGCGGACGGATCGATCGAGAAGTACAAGGCGCGCCTCGTGGCCAAGGGGTTCAAGCAGAAGTACGGCATCGACTACACGGAGACGTTCTCGCCCGTCGTCAAGTACGTGACGCTGCGCATGGTGATCGCGATTGCCAAGTATTTTGGCTGGCCCCTCGACCAACTCGACGTGGTGACTGCATTCCTGTACGGGATAATGAAGGAGCTGGTGTTCTGCGCGGTCCCCGAAGGTGTTGATCTCGACGGTGACTTCGACTGCCTGGAATTGGTGAAGGCGATCTACGGTCTCAAGCAAGCTTCGCGCGTGTGGAACGAGACCTTCGACGAGTTCGTGTGCTCCATCGGCTTCCAAGTATCGGCGTTCGACCCATGCCTCTACATCAAGATCGTTGACGGTCACTGCGTGCTCGTGCTGGTCTACGTGGATGACGTGCTGATTACCGGAAGTTCACCCGAACTGATCTCACGCACGAAGACCGACCTCAAGACACGCTTCGAGATGACCGACAGCGGCAAGTGCGCGTTCGTTCTTGGGATCGAGTTGGTGGACGGCCCGGACGGCAGCGTGACGATGTGTCAGCGCCGCTATGTGGATGACATTCTCAAGCGTTTTGGTATGGACGAGTGCAAGGCCGTGGTGAGTCCCGTCGACATAAGCACTCGACTGATTTCAAGTGACGCAGCGACCAAGGTGAACGCTCCGTTTCGCGAAGCTGTTGGTGCTCTGATGCACTTGATGACTGCCACGCGCCCGGACATCGCGTACGCCGTTGGCTATGTGTCGCGCTTCATGGAGAACCCTCAAGAAGAACACTGGGTTGCGGTCAAGCGTATCTTTCGCTACCTGCAAGGCACCAAGACGCATGGCATCTGCTTCAAGCCGGGTGACAACATCGACTTTCGTGGATACTCGGACGCTGACTGGGCTGGCGACCTCGCGGACCGCAAGTCGACTTCCGGCTACACGTTCATGCTGATGAGTGCGCCTGTGAGCTGGGGAAGCAAGAAGCAGTCAAGTGTGTCGCTGTCTACGAGCGAGGCGGAGTACATCGCGCTCAGTCTGGCTATCCAAGAGGGCAAGTGGATCCATCGCCTACTGTGCGAGATTTTGGCGGCTACCAACGAGACGGGACCCGAGCTCATGATCCGCGAAGACAACCAGTCGTGCATCAAGATGACGAAGAACCCAGTGAACCACGGTCGCGCCAAGCACATCGACATCAAGTACCACCACATCCGCGACGAAGTGAAACGCGGTGAAGTGAAGCTCGAGTATTGCGAGACTTCTATGATGCTGGCGGACATCATGACGAAGGCGCTCCGGGACCTCGTCACACGGACCTGACGGCTGCTCTCGGCATCCACGCGTGTTCGCATTGAGGGGCGTATTGAGGGTTGCTGGTCTTGGGTACTGTGTACCAGTTGTACCCATTAACCGGTCAACCCGAACCGGTCCAATACGAACCCACGTTGATTATGGTGATTACGGTAAGTTGTTTACGCCACCGATGATTAGACGAGTAGGATGATTGGACGAACACCGATGAGTGATGGTTGATTAGATGCACTAGGTGATAGGACGCAGGGTCGCGTGTCTATAACAGTAGGGTGTAAGCTCCAACAATTCAATCACTTTCGAATCTTCATTCTACCCCTCAAGTGTCTTCGCTCCGGTCCCGCCGTCTGCGACGTCCCTCCCTCCGCTTCGTCCAAGCTTCGTGTCCAAGATCTATACCTTCCATCAATCAGCACGTCATCCACGTAGACCAGCACGAGCACACAGTG

ATG – predicted start codon

TAG – predicted stop codon

The LTR marks (5’-TG..CA-3’) are underlined within the LTR regions highlighted in cyan.
